# The eutherian-specific histone H3.4 promotes germ cell development and reproductive fitness

**DOI:** 10.1101/2025.06.19.660366

**Authors:** Pavel A. Komarov, Philipp Bammer, Ching-Yeu Liang, Hans-Rudolf Hotz, Grigorios Fanourgakis, Sunwoo Chun, Hubertus Kohler, Tim-Oliver Buchholz, Jean-Francois Spetz, Antoine H.F.M Peters

**Author notes:** Corresponding author: Antoine H.F.M. Peters. shared authorship (Philipp Bammer, Ching-Yeu Liang).

## Abstract

Many genes encoding chromatin proteins are subject to evolutionary selection driving reproductive fitness. In mice and men, the histone H3.4 variant is essential to spermatogenesis. Here we define the evolutionary origin and molecular-physiological roles of sequence variation in *H3f4* for male germ cell development in mice. Our phylogenetic analyses indicate that eutherian *H3f4* orthologs originate from an ancestral *H3.2* gene existing prior to the divergence of eutherian and marsupial mammals over 100 million years ago. Positioned in truncated histone gene clusters, eutherian *H3f4* orthologs show increased non-synonymous and synonymous substitution rates compared to orthologous marsupial *H3.2* loci located in typically large histone clusters. To determine the impact of sequence divergence on reproductive fitness, we reverted non-synonymously substituted residues in *H3f4* to those present in canonical H3.1 (*H3f4^V24A^, H3f4^H42R^, H3f4^S98A^*). Hemizygous expression of such triply reverted *H3f4^H3.1^* allele on a *H3f4*-deficiency background caused an >40% reduction in testis weight associated with impaired meiotic DNA double strand break repair, death of pachytene spermatocytes, impaired differentiation of spermatids and aberrant expression of thousands of genes during spermatid elongation. Hemizygous expression of individual residue substitution alleles revealed residues V24 and H42 of H3.4 to promote spermatogenesis, while residue S98 is neutral. Together, our study shows that *H3f4* has been subject to positive evolutionary selection, promoting male reproductive fitness.

## Introduction

Spermatogenesis is a complex developmental process that yields to the daily formation of millions of haploid spermatozoa competent for supporting embryonic development. While the genome is packaged by histones during most of this process, histones become replaced by protamines during spermatid elongation, thereby compacting their chromatin which is required for fertility^1,2^. Contrasting somatic cells, germ cells express many histone variants, some serving prominent chromatin remodeling roles complementary to canonical histones^1,3,4^. In mammals, testicular isoforms exist within all histone families except for histone H4^5,6^. Multiple *H1, H2A, H2B* and *H3* genes are subject to rapid evolution including pseudogenization^7–10^.

Among H2A and H2B proteins, the TH2A and TH2B histones are highly expressed during male meiosis and replace over 60% of canonical H2A and H2B^11,12^. TH2A is required for meiotic progression^13,14^ and TH2B was shown to prime chromatin for histone-to-protamine replacement during spermatid elongation^11^. In accordance, TH2A/TH2B-containing nucleosomes make fewer contacts with DNA and exhibit reduced stability, compared to canonical nucleosomes^15,16^. Both proteins are also maternally expressed in oocytes and pre-implantation embryos where they contribute to transcriptional activation of the paternal genome^17^. Further, specialized short H2A isoforms lacking the C-terminal part (H2A.B and H2A.L) are expressed almost exclusively in the testis^7,18^. In mouse, histone H2A.B.3 is the most highly expressed H2A.B histone that is incorporated at transcription start sites (TSS) of active genes in spermatocytes and round spermatids^19,20^. However, all three *H2A.B* paralogs serve biparental contributions to embryonic development of mice^21^. The H2A.L.2 histone is incorporated during spermatid elongation and is required for the loading of transition proteins (TNPs) onto chromatin^22,23^. The short H2A isoforms (H2A.B, H2A.L, H2A.Q and H2A.P) have been identified only in eutherians^7^. They likely originate from H2A.R, a regular H2A isoform present only in marsupials and monotremes^7^. Importantly, short H2A genes underwent pseudogenization, or further duplication during eutherian evolution^7^.

Next to TH2B, H2B.L is expressed during spermatid elongation and plays a role in pericentric heterochromatin organization^22^. Another isoform, H2B.W.1, is expressed in human spermatogonia^24^, and has been implicated in male fertility^25,26^. Like H2A variants, the testis expressed eutherian-specific TH2B, H2B.W and H2B.L-encoding genes, present in all mammals, also underwent duplication and/or pseudogenization during evolution^8^. For example, *H2B.L* exists only as a pseudogene in the human genome^8^.

For H1 histone variants, H1T (H1.6) is expressed from pachytene spermatocytes until late spermatids^27^ in which it controls chromatin de-condensation at genic regions^28^ and heterochromatin maintenance at repeat elements^29^. The H1T2 (H1.7) and HILS1 (H1.9) isoforms are highly expressed during spermatid elongation^30,31^. H1T2 is required for fertility in mouse^31,32^ and human^33^. In human, *HILS1* is retained only as a pseudogene^34^.

Within the *H3* family, primates express novel *H3* isoforms, mostly originating from replication-independent (RI) H3.3-encoding genes and accumulating non-synonymous mutations. For example, H3.X and H3.Y are specific to primates^35,36^, while H3.5 exists in hominids^37^. H3.5 is expressed in the spermatogonia and primary spermatocytes^38,39^, and reduced *H3-5* transcripts in sperm were associated with non-obstructive azoospermia^39^. In mice, thirteen *H3* genes were predicted to have evolved from *H3.3*-encoding genes, with some being specifically expressed in the testis^10^.

Next to *H3.3* variant-related genes, the H3.4 variant (also known as *H3t*)^34^ is mainly expressed in spermatogenic cells^10,22,40–44^. H3.4 is encoded by a single *H3f4* gene possessing a 3’-UTR stem loop^10,45^, and is incorporated into chromatin in a replication-coupled (RC) manner^10^. The nuclei of c-KIT positive differentiating spermatogonia, meiotic spermatocytes and haploid round spermatids are widely labeled by H3.4^42^. However, H3.4 is removed from the sex chromosomes during meiotic sex chromosome inactivation (MSCI) process^42,43,46^. The precise developmental timing of H3.4 deposition and eviction, and its genomic distribution remain unexplored. In mouse, *H3f4* is essential to spermatogenesis, with spermatogonial differentiation failing in its absence^42^. In human, the c190C>T single nucleotide polymorphism (SNP) encoding the R64C substitution in *H3-4* has been associated with a Sertoli cell only syndrome (SCOS)^47^.

Given the pivotal role of *H3f4* and *H3-4* in male fertility, it is intriguing why spermatogenesis became dependent on the gene, despite the presence of multiple canonical RC *H3* genes in eutherian genomes. In human, *H3-4* is a member of the small *HIST3* gene cluster that also contains the *H2AC25, H2BC26* and *H2BC27* histone genes and which is flanked by the *TRIM11, TRIM17* and *RNF187* genes (Fig. 1b). The composition of *HIST3* cluster is largely conserved in other eutherian species, including apes, dog, cat, pig, horse and mouse^34^. The origin of *H3-4* and its conservation among other mammals are, however, unknown.

**Figure 1.**
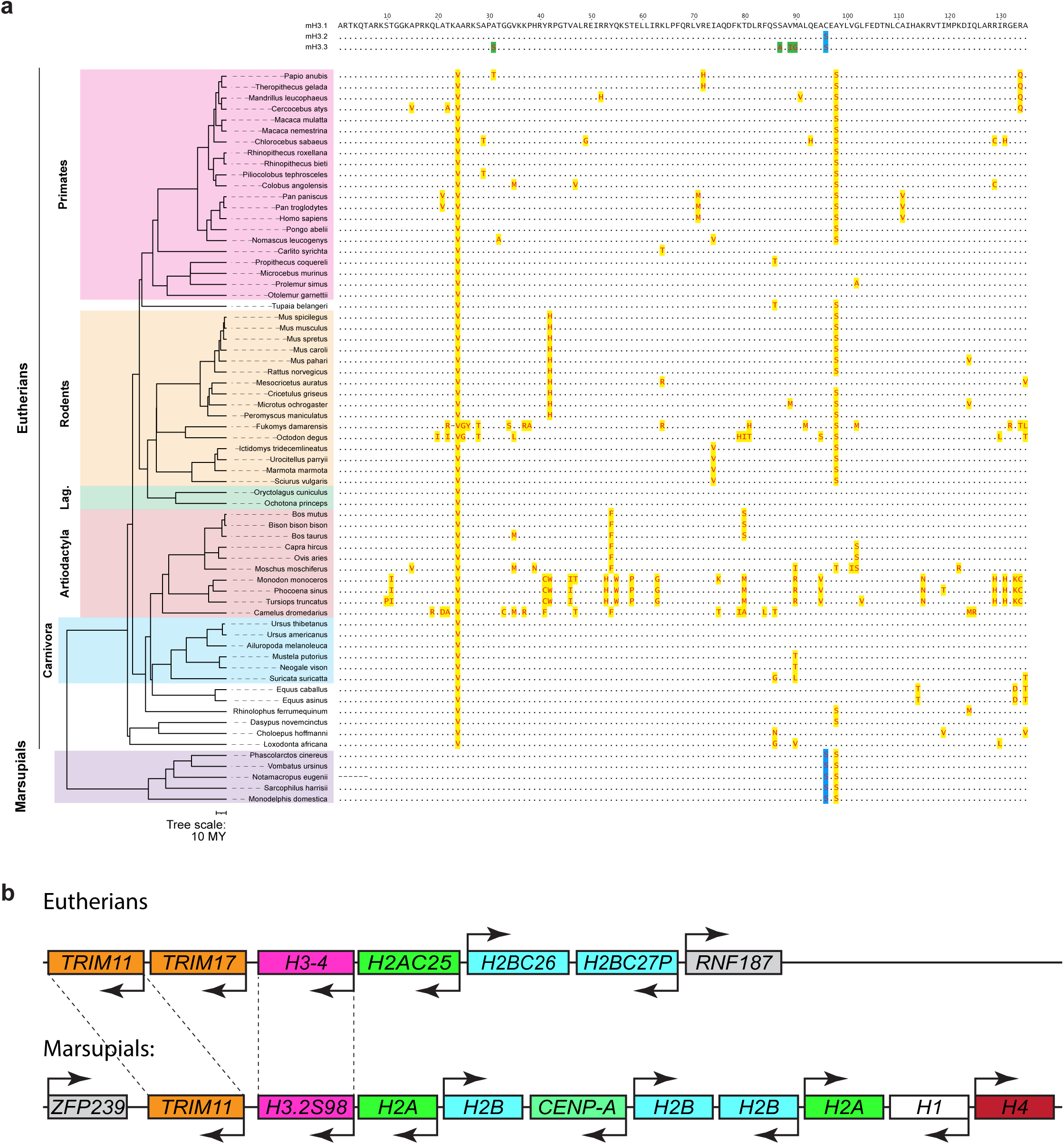
Sequence divergence of histone H3.4 in the mammalian lineage. **a**, Alignment of amino acid sequences of histones H3.1, H3.2 and H3.3 (mH3.1, mH3.2, mH3.3) in mouse and H3.4 in the indicated species. Conserved residues relative to mH3.1 are represented as dots while amino acid sequence variations are highlighted in yellow. H3.2 and H3.3 specific substitutions are highlighted in blue and green, respectively. Species are ordered according to their position in the phylogenetic tree representing over 100 million years of divergence. **b**, Schematic representation of the *H3-4* locus and neighbouring genes in placental (human, top) and marsupial (opossum, bottom) species. Scaled representation is shown in Supplementary Fig. 1a.

In human, four amino acids differ between H3.4 and canonical H3.1 (V24A, M71V, S98A and V111A). *In vitro*, H3.4-specific residues M71 and V111 reduce the stability of nucleosomes^48,49^. Moreover, H3.4 interacts preferentially via its residue V111 with the histone chaperone hNAP2 over hNAP1 thereby promoting nucleosome formation *in vitro*^50^. The Tudor domains of PHF1 and PHF19, two accessory components of the Polycomb Repressive Complex (PRC) 2 complex, preferentially bind H3.4K27me3 over canonical H3.1K27me3 via residue V24^44,51^ suggesting a role for H3.4 in PcG-mediated gene repression. In mouse, while V24 and S98 are present, M71 and V111 residues are not (Fig. 1a). Instead, mouse H3.4 harbors a histidine rather than an arginine at position 42 which promotes open chromatin *in vitro*^42^. The *in vivo* impact of the evolutionary acquired amino acid substitutions in H3.4 for spermatogenesis and reproductive fitness are unknown.

In this study, we investigated the evolutionary origin of *H3-4* orthologous genes and assessed whether acquired amino acid substitutions may regulate fertility traits. We identify that the eutherian-specific *H3f4* evolved from a *H3.2* gene existing over 100 million years ago, prior to separation of eutherian and marsupial lineages. Following the high expression of *H3f4*, H3.4 replaces progressively canonical H3 histones during spermatogonial expansion and differentiation. In proliferating spermatogonia, RC-deposition of H3.4 occurs genome-wide, whereas in post-replicative spermatocytes and spermatids, H3.4 remains located at transcriptionally silent and intergenic regions characterized by low nucleosome turnover. In turn, at transcriptional start sites (TSSs) of active genes and on chromosome X in pachytene spermatocytes, H3.4 is gradually evicted and replaced by H3.3. Through gene editing, we generated a mouse model expressing H3.1 from the *H3f4* locus. Under conditions of reduced gene dosage, our results show that H3.4 supports spermatogenesis better than H3.1, with V24 and H42 amino acids playing key roles in the function of H3.4.

## Results

### The *H3-4* gene is conserved in the eutherian lineage

To determine the phylogeny of the *H3-4* gene and the level of its amino acid conservation, we searched for *H3-4* orthologs in a wide set of eutherian species (Fig. 1a, Supplementary Table 1). We performed TBLASTN searches using the human H3.4 protein sequence as a query against the genomes available at the Ensembl website (www.ensembl.org, release 112) and defined *H3-4* orthologs as those *H3* genes localized within the “*HIST3”* gene cluster containing *TRIM11* and/or *TRIM17, H2A* and several *H2B* genes^34^ (Supplementary Fig. 1a). The presence of a valine residue at position 24 (V24) is the distinctive feature among all H3.4 orthologs identified in 62 species analyzed, contrasting with alanine (A24) universally present in canonical H3.1, H3.2 and variant H3.3 proteins (Fig. 1a, Supplementary Fig. 2a-c). Beyond H3 genes within orthologous *HIST3* gene clusters, we were not able to identify *H3-4* paralogs encoding H3 proteins with V24. We conclude that H3.4 is encoded by a single gene localized within *TRIM11*/*TRIM17, H2A* and *H2B* containing *HIST3*-like gene clusters in eutherian mammals which diverged over 100 million years (Fig. 1b).

### The eutherian *HIST3* cluster likely originates from a large histone cluster still existing in marsupial mammals

To further identify the origin of the *H3-4* containing *HIST3* cluster, we expanded our search for putative *H3-4* orthologs in marsupial species, requiring linkage with a neighboring *TRIM* gene as a selective genome anchor. We identified a large canonical replication-dependent histone cluster consisting of one *H3*, two *H2A*, three *H2B* genes and other histone genes (*H1, H4, CENP-A*) in genomes of four marsupial species analyzed (Koala, Opossum, Tasmanian Devil and Wombat) (Fig. 1b, Supplementary Fig. 1a, Supplementary Table 2). Intriguingly, in contrast to eutherian *H3-4*, the *H3* genes linked to *TRIM* genes in marsupial genomes do not encode for H3^V24^ proteins but for regular H3^A24^ proteins (Fig. 1A). Moreover, these H3 proteins contain a serine at residue 96 (S96), characteristic of canonical H3.2 replication-dependent histones in eutherian and marsupial mammals, and not for a cysteine (C96) typical for H3.1 and H3.4 proteins (Fig. 1a and Supplementary Fig. 2). Finally, these H3 proteins harbor a serine at residue 98 (S98) which is frequently present in eutherian H3.4 orthologs suggesting common origin (Fig. 1a, Supplementary Fig. 2 and Supplementary Table 1). Importantly, no other H3.2 genes in marsupial genomes encoded S98 (Supplementary Fig. 2b). Based on these data, we conclude that the small eutherian *HIST3* clusters evolved from a large regular histone cluster found in marsupials, with *H3-4* originating from a *H3.2^S98^*gene.

### Increased sequence variation among *H3-4* orthologs

*H3-4* orthologs display a high degree of sequence variation among eutherian mammals with certain amino acid polymorphism conserved between evolutionary related species (Fig. 1a). For example, residues M71 and V111 are found in closely related human, chimp and bonobo, Q134 is restricted to drill, sooty mangabey, gelada and olive baboon and V74 is limited to squirrels. Residue H42 is specific to muroids (mice, rats and hamsters). The extensive sequence variation may reflect reduced selective pressure to maintaining the canonical H3 sequence, allowing possible functional adaptation to germ cell development.

To directly compare the degrees of sequence conservation among H3.4 and regular H3 histones, we selected several eutherian (human, lemur, mouse, rabbit, cow, camel and elephant) and marsupial (wallaby, Tasmanian devil, opossum, koala and common wombat) species across the evolutionary time axis. We retrieved sequences of all *H3.1, H3.2* and *H3.3* genes in these 12 species from the Ensembl database (release 112) (Supplementary Table 3). Among the selected species, the alignment of coding sequences (CDS) of *H3-4* orthologs showed a substantial degree of variation (Supplementary Fig. 1b). The vast majority of variations are at the third position of the triplet and thus do not encode a change in amino acid. Nonetheless, the rate of non-synonymous mutations among *H3-4* orthologs was high across the selected species (Fig. 1a). In contrast, we barely observed any variation in amino acid composition among H3.1 and H3.2 proteins of selected eutherian and marsupial species including the (Supplementary Fig. 2a,b). The genes encoding H3.3 displayed also only minor protein sequence divergence (Supplementary Fig. 2c). Therefore, contrasting other H3 isoforms, *H3-4* genes appear indeed to be subject to reduced selective evolutionary pressure for conserving their sequence.

We next investigated whether the relaxation of sequence conservation in *H3f4* adapted the function of H3.4 in spermatogenesis. To address this question, we studied H3.4 expression dynamics, cellular and genomic localization and its functional role in male germ cell development.

### Dynamic *H3f4* and *H3.3* expression during spermatogenesis

We first profiled the expression dynamics of histone transcripts in seven FACS-purified cell types covering the entire process of spermatogenesis, from undifferentiated spermatogonia to elongated spermatids^52,53^ Principle component analysis (PCA) on UCSC-annotated protein-coding genes demonstrated a clear separation of different cell types, reflecting the developmental trajectory of developing male germ cells (Supplementary Fig. 3a). In undifferentiated and differentiating spermatogonia (SgU, SgD) and in early meiotic leptotene/zygotene stage spermatocytes (ScLZ), most canonical *H2A, H2B, H3* and *H4* genes were amply expressed (Fig. 2a, Supplementary Fig. 3b). Remarkably, *H3f4* expression exceeded expression of H3.1- and H3.2-encoding genes, likely promoting H3.4 deposition during replication. Upon progression through meiotic prophase and in haploid spermatids transcript levels of all RC histones including *H3f4* decreased, while those of RI *H3.3* variants remain high^54^ thereby supporting nucleosome turn-over beyond replication^46^. Together, we measured a competitively high *H3f4* expression during the replicative phase of spermatogenesis, while *H3.3* expression takes over during meiosis and spermatid development.

**Figure 2.**
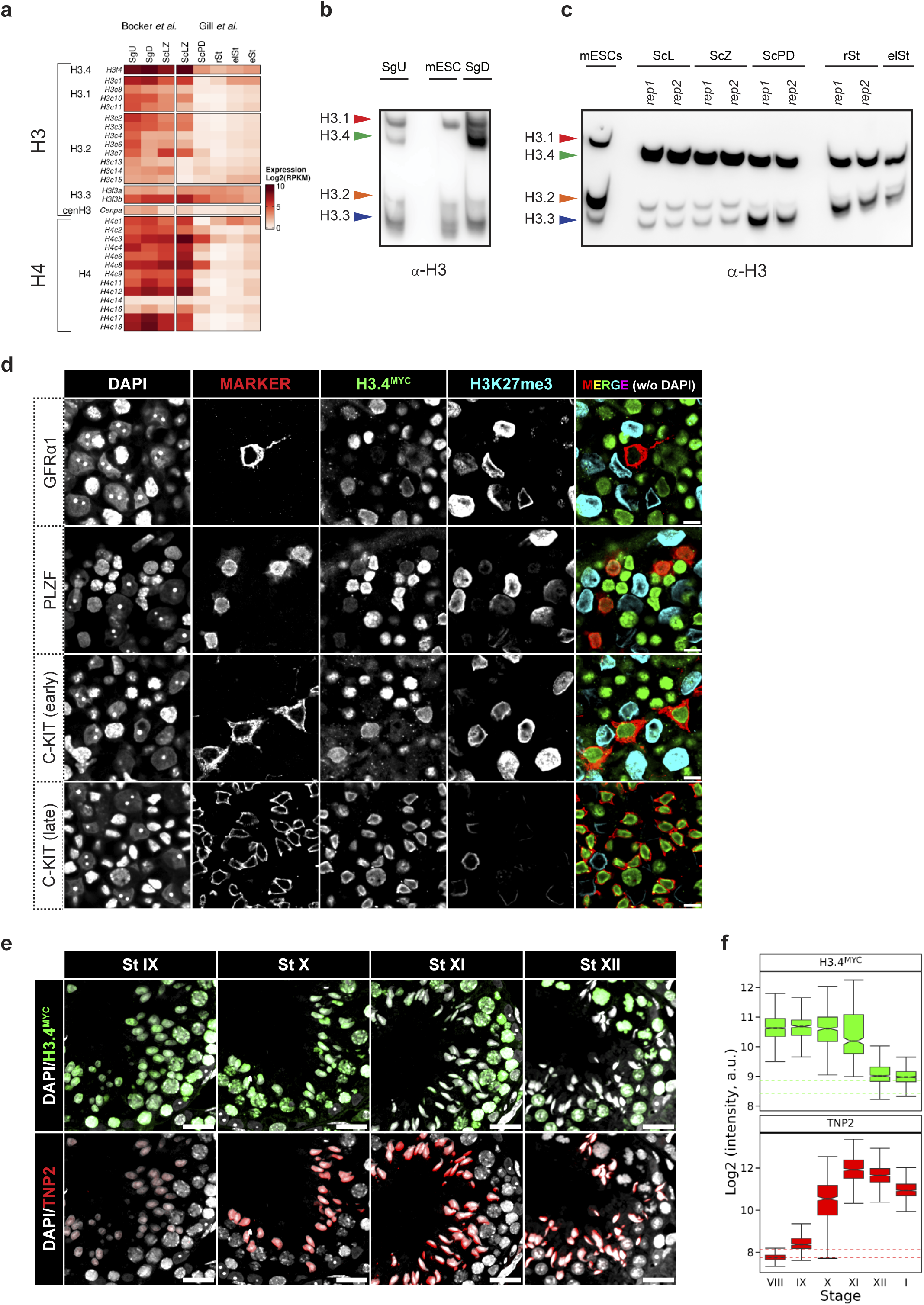
Expression dynamics of histone H3.4 during spermatogenesis. **a**, Heatmap showing mRNA expression of H3 and H4 encoding genes during spermatogenesis^52,53^. Gene names are according to the current nomenclature^34^. Abbreviations: SgU, SgD, ScLZ, ScPD, rSt, elSt and eSt refer to populations of FACS-sorted undifferentiated and differentiating spermatogonia, leptotene/zygotene and pachytene/diplotene stage spermatocytes and round, elongating and elongated spermatids. **b-c**, Western blots showing H3.1, H3.4, H3.2 and H3.3 proteins separated by TAU gel electrophoresis in FACS-purified spermatogonia (**b**) and spermatocytes and spermatids (**c**). We used mouse embryonic stem cells (ESCs) as controls and used a panH3 antibody recognizing the C-terminus of H3 to detect all H3 isoforms. Additional abbreviations: ScL: leptotene spermatocytes, ScZ: zygotene spermatocytes. **d**, Whole-mount immunofluorescence images of H3.4^MYC^ expression in different populations of spermatogonia, marked by GFRα1, PLZF or c-KIT. H3K27me3 marks Sertoli cells. Scale bar: 10 μm. **e**, Representative immunofluorescence images of stage IX to XII seminiferous tubules, co-stained for H3.4^MYC^, TNP2 and DAPI. Scale bar: 20 μm. **f**, Quantification of H3.4^MYC^ and TNP2 signals in step 8-13 spermatids (from stage VIII to I tubules). The dashed lines represent the Q25 and Q75 intensity levels of the respective signals in Sertoli cells. Staging was performed manually as described before^61^.

### H3.4 is the main H3 protein in differentiating spermatogonia

To investigate the relative protein levels of different H3 isoforms during male germ cell development we employed triton - acetic acid - urea (TAU) gel electrophoresis combined with immunoblot detection using a pan-H3 antibody (Supplementary Table 9), which enables separation and detection of different H3 isoforms based on their electric charge and hydrophobicity^55^. Probing acid-extracted histone samples obtained from testis, various somatic tissues and ESCs from mice, we observed comparable H3.1, H3.2 and H3.3 expression levels in most somatic tissues (Supplementary Fig. 3c). In ESCs, H3.1 and H3.2 were more abundant than H3.3, presumably related to their high proliferative nature. In testis, H3.1 and H3.2 levels were remarkably low compared to H3.3 and an abundant H3 protein migrating faster than H3.1. Mass spectrometry identified the faster migrating protein as H3.4, given the presence of the “KQLATKV_24_AR” peptide and absence of the “KQLATKA_24_AR” peptide diagnostic for other H3 proteins (Supplementary Fig. 3d). TAU gel electrophoresis of the H3.4 protein expressed in HEK293 cells confirmed the migration pattern of H3.4 (Supplementary Fig. 3e). These analyses revealed a leading expression of H3.4 over H3.1 and H3.2 in testis and its absence in other somatic samples, underscoring its testis related expression, as previously noted^10,22,40–42^.

We next profiled H3 protein levels in FACS-purified testicular male germ cells. While somatic H3 isoforms were prevalent, H3.4 was readily detectable in SgU cells (Fig. 2b). As spermatogonial proliferation and differentiation proceeded, H3.4 became the predominant H3 protein in c-KIT expressing SgD cells (Fig. 2b). In post-replicative meiotic prophase cells, H3.3 levels majorly increased from pachytene stage onwards, possibly reflecting increased transcription-coupled nucleosome turnover^56^, as well as nucleosome replacement at sex chromosomes during MSCI^46,57^ (Fig. 2c). Together, the TAU-WB data argue for a RC replacement of most canonical H3.1 and H3.2-containing nucleosomes by H3.4-bearing nucleosomes during spermatogonial amplification.

### Low expression of C-term. tagged H3.4^MYC^ is compatible with male germ cell development

To further study the developmental dynamics and genomic localization of H3.4, we aimed for tagging it with small exogenous tags, an approach commonly used in somatic cell studies (Supplementary Fig. 4a). Using CRISPR/Cas9 gene editing transgenesis, we successfully added a MYC and an HA tag to the N-terminus (*^MYC^H3f4* and *^HA^H3f4*) or a MYC and a 3xFLAG tag to the C-terminus of H3.4 (*H3f4^MYC^* and *H3f4^3xFLAG^*). We added a flexible linker between H3.4 and the tags to avoid disruption of protein function (Supplementary Fig. 4a). Nonetheless, except for the *H3f4^MYC^*allele, spermatogenesis and male fertility were severely impaired in all transgenic mouse strains, even when the tagged H3.4 proteins were heterozygously expressed (*H3f4^TAG/wt^*) (Supplementary Fig. 4b). In contrast, two independently generated *H3f4^MYC^* alleles showed overall normal progression through spermatogenesis when heterozygously expressed, even though testis weights were ∼20% reduced (Supplementary Fig. 4c). Homozygous *H3f4^MYC/MYC^* expression impaired spermatogenesis during meiotic prophase with a phenotype less severe than observed for the other three transgenic tagged mouse strains showing impaired spermatogonial development (Supplementary Fig. 4b, 4c). Titration experiments of FACS-isolated spermatocytes from *H3f4^MYC/wt^* mice showed that H3.4^MYC^ was expressed at a level of ∼20% of non-tagged H3.1/H3.2/H3.4 proteins (Supplementary Fig. 4d). We next used the low expressing transgenic *H3f4^MYC/wt^* mice to investigate the dynamics of H3.4 deposition, removal and its genomic distribution.

### H3.4 deposition into chromatin starts in spermatogonial stem cells and accumulates until entry into meiosis

To determine the onset of H3.4 deposition into chromatin, we recorded the presence of H3.4^MYC^ in populations of GRFα1 and PLZF positive SgU cells and c-KIT positive SgDs^58^ by immunofluorescence microscopy on whole mount testicular tubules. H3.4^MYC^ is already detectable in nuclei of about 70% of GRFα1^+^ type A_single_ (A_s_) and A_paired_ (A_pr_) SgU cells and in all clones of type A_aligned_ (A_al_) SgU cells containing 4 to 8 germ cells. The intensity of H3.4^MYC^ signals increased parallel to the clone size of GRFα1^+^ and PLZF^+^ SgU and reached a plateau in c-KIT^+^ SgD cells (Fig. 2d). In contrast, H3.4^MYC^ was not detectable in somatic Sertoli cells marked by H3K27me3 (Fig. 2d, Supplementary Fig. 5a, 5b). In summary, these data suggest a progressive incorporation of H3.4 over multiple rounds of replication into spermatogonia starting already in undifferentiated GFRα1-positive A_s_ spermatogonia^59,60^.

### H3.4 is evicted from chromatin in step 11 and 12 elongating spermatids

From c-KIT^+^ SgD onwards, nuclear H3.4 levels remained high in spermatocytes and spermatids until their nuclear elongation. To identify the timing of H3.4 eviction from chromatin, we co-stained paraffin-embedded *H3f4^MYC/wt^* testis sections for MYC and TNP2, a marker of elongating spermatids^1^. We manually staged seminiferous tubules^61^ and quantified intensities of anti-MYC and anti-TNP2 signals in step 8-13 spermatids. While TNP2 incorporation started in stage IX and gradually increased, H3.4^MYC^ signals remained high until stage XI, where H3.4 eviction began (Fig. 2e,f). At stage XII, step 12 elongating spermatids demonstrated a loss of H3.4^MYC^ signal, reaching intensities as low as measured in somatic Sertoli cells (Fig. 2e,f). We confirmed the rapid timing of H3.4 eviction using an antibody recognizing all replication dependent H3.1, H3.2 and H3.4 proteins^62^ (Supplementary Fig. 8c). Together, our data indicates that most RC-H3.4 proteins are removed at the end of nuclear elongation, in stage XII spermatids.

### H3.4 is localized throughout the genome and gets replaced by H3.3 at specific genome regions in post-mitotic germ cells

Next, we investigated the genomic distribution of H3.4^MYC^ in FACS-purified spermatogonia, meiotic spermatocytes and haloid spermatids using ultra-low input^63,64^ and regular ChIP-seq protocols (see materials and methods for details). We first studied the occupancy of H3.4^MYC^ within regions of ±5 kb around transcriptional start sites of genes that we had classified into nine k-mean clusters according to having a promoter with or without a CpG island, occupancies of H3.4^MYC^, H3.3, H3K4me3, H3K27me3, and to associated mRNA expression (Fig. 3a-c, Supplementary Fig. 6a-c). Consistent with global protein and imaging data (Fig. 2b, 2d), we readily observed H3.4 occupancy in SgU, which increased at TSS in SgD, both at CGI and non-CGI type promoter genes. Along the developmental trajectory from SgD to rSts, H3.4 became gradually evicted and replaced by H3.3 at TSS of genes in a sequence, histone modification and transcription related manner (Fig. 3a,b, Supplementary Fig. 6a,b).

**Figure 3.**
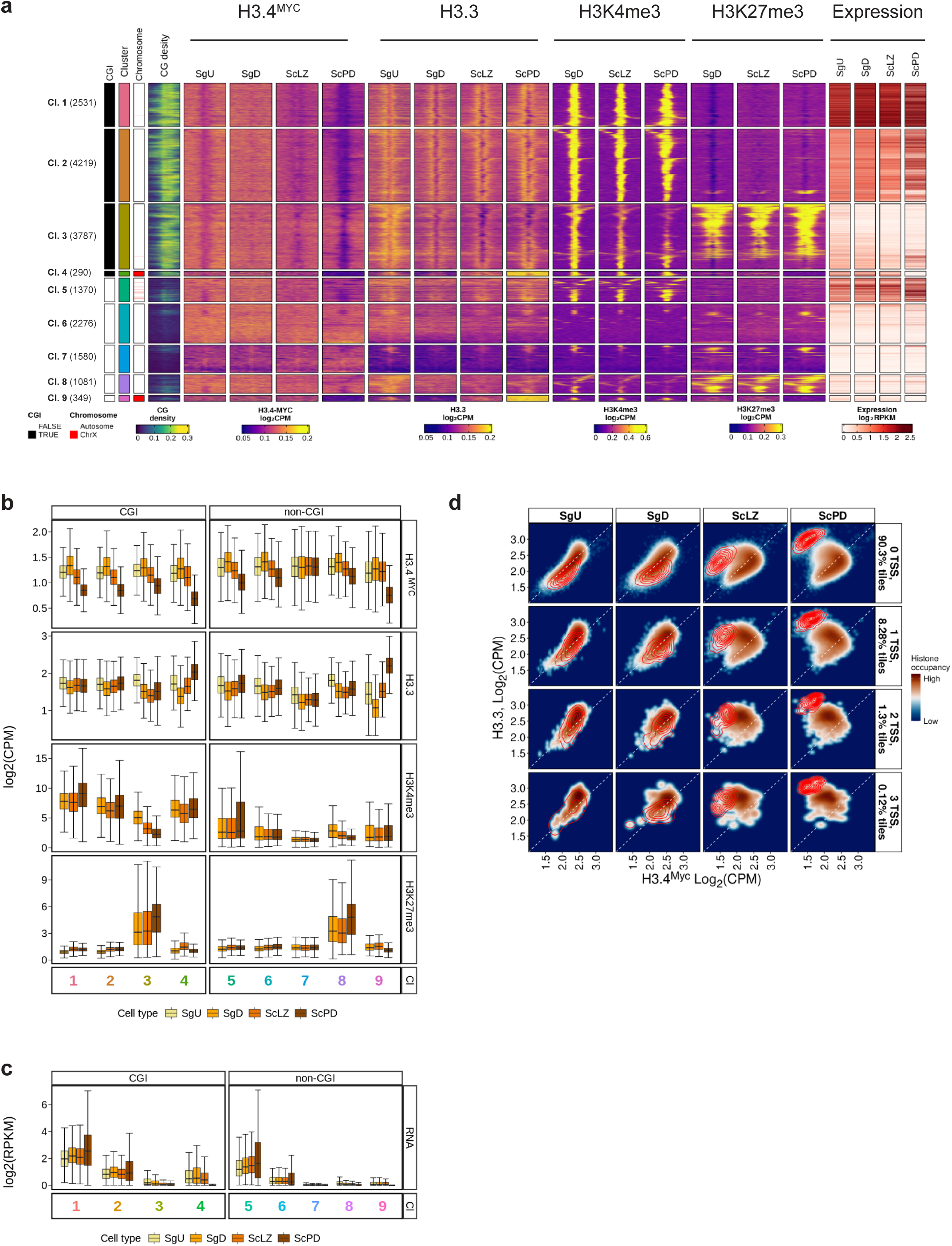
Genome-wide analysis of H3.4^MYC^ and H3.3 occupancy during spermatogenesis. **a**, Heatmap displaying chromatin and transcriptional states at TSS regions (± 5 kb) of CGI- and nonCGI-containing genes measured in FACS-purified SgU, SgD, ScLZ and ScPD cells. Regions were grouped into 9 gene clusters by k-means clustering performed separately for the CGI- and for the non-CGI-type promoters performed on chromatin and RNA variables. The number of gene promoters is displayed for each cluster. CG density (±1 kb around TSS), chromatin and RNA expression levels are indicated in percentage, logCPM and log2RPKM respectively. The ChIP-seq data related to H3K4me3, H3K27me3 in SgD cells, H3K27me3 in ScLZ and ScPD, as well as RNA expression in SgU, SgD and ScLZ cells were obtained from Bocker *et al.*^52^. The RNA-seq data for the ScPD population has been obtained from Gill *et al.*^53^. **b**, Boxplot showing quantification of ChIP-seq data (in log2CPM) in windows ±1 kb around TSS in clusters 1-9 as displayed in panel **a.** The boxes extend from the first quartile (Q1) to the third quartile (Q3) of the data distribution, with a line in the box representing the median. Whiskers indicate variability outside Q1 and Q3, extending to the minimum and maximum values within 1.5 times the interquartile range (IQR), while points outside this range represent outliers. **c**, Boxplot showing quantification of expression of genes (in log2RPKM) belonging to the TSS regions displayed in clusters 1-9 in panel **a**. Boxplot parameters are as described in panel **b**. **d**, Density scatter plots displaying log2CPM counts of H3.4^MYC^ (x-axis) and H3.3 (y-axis) on genomic tiles that contain 0, 1, 2 or 3+ transcription units with TSS in SgU, SgD, ScLZ and ScPD cells. Red contour lines represent the density of tiles located on the X chromosome. The presence of a transcription unit with TSS was defined by an overlap of a given tile with a region ±1 kb around TSS.

We measured the highest H3.4^MYC^ to H3.3 replacement at CGI- and non-CGI-type promoters of actively expressed genes marked by H3K4me3 (Fig. 3a-c, Supplementary Fig. 6a-c, clusters 1, 2 and 5). Moderate levels of replacement were observed at promoters of H3K27me3-marked genes, and of one class of non-CGI promoter genes that are barely expressed (Fig. 3a-c, Supplementary Fig. 6a-c, clusters 3, 6 and 8). Lastly, class 7 with non-CGI promoter genes that are barely expressed displayed hardly any H3.4^MYC^ to H3.3 conversion across any cell types (Fig. 3a-c, Supplementary Fig. 6a-c).

Next, we investigated genome-wide localization of H3.4^MYC^ and H3.3. Hence, we calculated ChIP-seq signals of H3.4^MYC^ and H3.3 in 5 kb non-overlapping genomic tiles that we had classified according to the number of TSS-containing transcription units present and their chromosomal location (Fig. 3d and Supplementary Fig. 6d). On autosomes, genic and intergenic tiles showed comparable ChIP-seq signals for H3.4^MYC^ and H3.3 proteins in SgU and SgD spermatogonia. Upon entry into meiotic prophase and towards haploid round spermatids, however, H3.3 signals progressively increased at the expense of H3.4^MYC^ at many tiles, and such replacement was more pronounced for tiles containing one or more TSS-containing transcription units. The absence of replication-coupled H3.4^MYC^ deposition in post-mitotic cells, even within intergenic regions, is reflected by the progressive replacement by H3.3.

Contrary to autosomal genes, genes localized on the X chromosome displayed remarkably different temporal changes in chromatin composition at their promoters irrespective of their sequence, histone PTM and expression states (Fig. 3a-c, Supplementary Fig. 6a-c, clusters 4 and 9). In SgU cells, X-linked gene promoter regions were packaged by H3.4^MYC^ and H3.3 alike autosomal genes. Then, while H3.4^MYC^ remained present in SgD and early meiotic ScLZ cells, it majorly decreased in ScPD cells, presumably as part of MSCI. In contrast, H3.3 levels already increased in ScLZ, and further to highest levels in ScPD and rSts. Hence, X-linked genes initiate the exchange of H3.4^MYC^ to H3.3-holding nucleosomes at gene promoters already early in meiosis, several days prior to the chromosome-wide exchange observed in mid-pachytene spermatocytes (Supplementary Fig. 7).

As seen at TSS regions, we observed also a replacement of H3.4^MYC^ by H3.3 in X-linked genomic tiles of all four classes already early in ScLZ cells, even prior to the pachytene stage at which X-chromosome wide MSCI-associated eviction of replication dependent nucleosomes has been reported to occur (Fig. 3a)^46,57^. These data underscore the sensitivity of ChIP-seq over immunofluorescence in detecting X-linked nucleosome turnover, as confirmed by the exclusion of H3.4^MYC^ signal along the X and Y chromosomes only in mid-pachytene but not in early-pachytene spermatocytes as seen by imaging (Supplementary Fig. 7).

### H3.4 supports spermatogenesis more proficiently than H3.1

In mouse, *H3f4* carries four non-synonymous substitutions relative to *H3.2* genes. To understand the impact of these amino acid alterations on reproductive fitness, we performed CRISPR/Cas9 gene editing and reverted three of them to those present in *H3.1* (*H3f4^V24A^*; *H3f4^H42R^* or *H3f4^S98A^*) while leaving *H3f4^C96^* intact since this residue is present in H3.1 (Fig. 4a, Supplementary Fig. 8a, 8b). TAU gel electrophoresis confirmed for *H3f4^H3.1/H3.1^* animals the absence of H3.4 and continued presence of H3.1 in male germ cells throughout their development from SgU to eSt (Fig. 4b). Importantly, by expressing H3.1 from the *H3f4* locus, we did not alter global levels of histone H3, as measured by Western blot analysis (Fig. 4c).

**Figure 4.**
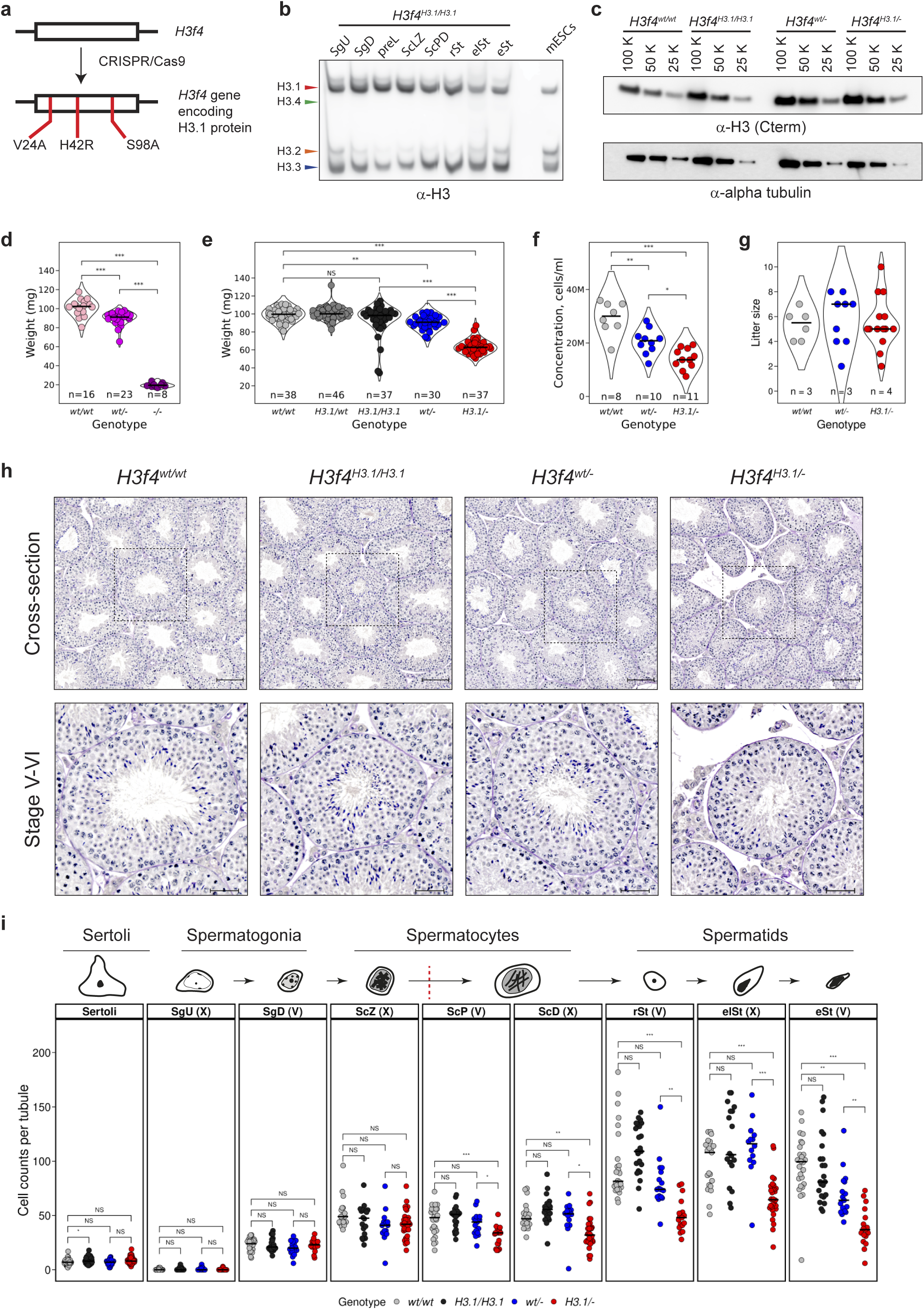
Reduced efficiency of spermatogenesis in *H3f4^H3.1/-^* substitution mice. **a**, Experimental design of CRISPR/Cas9-mediated editing of *H3f4* resulting in the replacement of H3.4-specific amino acids with those present in H3.1. Sequences of sgRNAs and repair template are provided in Supplementary Tables 5 and 6. **b**, Western blot showing the presence of H3.1, H3.2 and H3.3 proteins and absence of H3.4 in germ cells isolated, FACS-purified from *H3f4^H3.1/H3.1^*mice and separated by TAU gel electrophoresis. Mouse embryonic stem cells (mESCs) were used as control. The panH3 antibody recognizing the C-terminus of H3 was used to detect all H3 isoforms. **c**, SDS-PAGE / Western blot titration of H3 proteins present in 100, 50 and 25 thousand rSts isolated and FACS-purified from indicated genotypes. Immuno-based detection was done as described for panel b. Alpha tubulin was used as a loading control. **d**. Violin plot showing testis weights in *H3f4^wt/wt^, H3f4^wt/-^* and *H3f4^-/-^* mice. Each dot is a weight mean of two testes taken from each mouse. Bars represent median values of genotypes. The number of individual measurements (n) is indicated for each genotype. Paired t-test was performed between the indicated genotypes. * P < 0.05, ** P ≤ 0.01, *** P ≤ 0.001, NS – not significant. **e**, Violin plots showing testis weights (mg) of *H3f4^w/wt^, H3f4^H3.1/wt^, H3f4^H3.1/H3.1^, H3f4^wt/-^* and *H3f4^H3.1/-^* mice. Each dot represents the mean weight of two testes isolated from each mouse. Bars represent median values of genotypes. The number of individual measurements (n) is indicated for each genotype. The number of individual males tested (n) is indicated for each genotype. Statistical testing was performed as described for panel d. **f**, Sperm counts (cells/ml) of *H3f4^w/wt^, H3f4^wt/-^* and *H3f4^H3.1/-^* males. Each dot represents sperm counts from an individual male. Bars represent median values of genotypes. The number of individual males tested (n) is indicated for each genotype. Statistical testing was performed as described for panel d. **g**, Comparison of breeding efficiency of *H3f4^w/wt^, H3f4^wt/-^*and *H3f4^H3.1/-^* males. Each dot represents the number of pups born per litter. Bars represent median values for each genotype. The number of males (n) used in breeding tests are indicated for each genotype. Statistical testing was performed as described for panel d. **h**, PAS-haematoxylin-stained testicular cross-sections obtained from *H3f4^w/wt^, H3f4^H3.1/H3.1^, H3f4^wt/-^* and *H3f4^H3.1/-^* mice. Scale bars: 100 µm for cross-section overview and 50 µm for stage V-VI zoom-in. **i**, Quantification of indicated germ cell types at stage V and stage X tubules in *H3f4^w/wt^, H3f4^H3.1/H3.1^, H3f4^wt/-^* and *H3f4^H3.1/-^* mice. Each dot indicates the number of cells counted per tubule of the selected genotype. For each genotype, counts of 3 biological replicates were pooled together for the analysis. For each population, paired t-test was performed between the indicated genotypes. * P < 0.05, ** P ≤ 0.01, *** P ≤ 0.001, NS – not significant

Interestingly, expressing H3.1 instead of H3.4 did not affect spermatogenesis with testicular weights being unchanged between the *H3f4^H3.1/H3.1^* and *H3f4^wt/wt^* males (Fig. 4e). To further challenge the function of *H3f4^H3.1^*, we reduced its gene dosage during spermatogenesis. To this end, we generated a *H3f4* deficient allele (*H3f4^-^,* Supplementary Fig. 9a, 9b) and combined it with the *H3f4^H3.1^* allele (*H3f4^H3.1/-^*). By large, the overall proteins levels of H3.4 and H3.1 in FACS-purified rSts were comparable when being expressed from either one or two alleles (Fig. 4c). Nonetheless, in line with a previous report^42^, we observed gene dosage dependent effects of *H3f4* expression on male germ cell development. Compared to control wild type males, heterozygous *H3f4^wt/-^*males showed a small reduction in testis weight of 10% (P < 0.001) and a ∼25% reduction in sperm counts (P<0.01), but did not lose fertility (Fig. 4f-g). In contrast, homozygous deletion of *H3f4* (*H3f4^-/-^*) caused a severe reduction in testis weight by 80%, extensive degeneration of germ cells and absence of epididymal sperm (Fig. 4d, Supplementary Fig. 9c). When combining the *H3f4^H3.1^* and *H3f4^-^* alleles, the resulting *H3f4^H3.1/-^* males displayed a further decrease in testicular weight by over 30% (P< 0.001) and an additional ∼25% reduction in sperm counts (P<0.05), compared to control heterozygotes (*H3f4^wt/-^*). Nonetheless, *H3f4^H3.1/-^* males were still able to reproduce (Fig. 4g-h, Supplementary Fig. 8d). Together, these data indicate that H3.4 is more proficient than H3.1 in supporting male germ cell development.

### Substitution of H3.4 to H3.1 leads to the reduction of pachytene spermatocytes

Histologically, we next reviewed testicular sections and observed notably reduced numbers of spermatocytes and spermatids, only in *H3f4^H3.1/-^*males (Fig. 4h). To determine quantitatively at what stage germ cell development was affected in *H3f4^H3.1/-^* relative to control males, we developed a partially automated image analysis workflow to classify and count germ cells (Supplementary Fig. 10 and methods). We quantified spermatogonia, spermatocytes and spermatids present at stages V and X of the seminiferous epithelium cycle, in triplicate per genotype (Fig. 4i). While the number of SgU, SgD and early meiotic ScZ cells were comparable between all genotypes, the number of spermatocytes at mid-pachytene (ScP, at stage V) was ∼20% reduced only in *H3f4^H3.1/-^* males compared to *H3f4^wt/wt^* controls (P < 0.05), and *H3f4^H3.1/H3.1^* and *H3f4^wt/-^* genotypes. The number of diplotene spermatocytes (ScD, at stage X) and round, elongating and elongated spermatids were in *H3f4^H3.1/-^*males also reduced, indicating an important role for H3.4 in meiosis and possibly spermatid differentiation. In *H3f4^wt/-^* males, the number of elongated spermatids was ∼40% reduced (P < 0.01), which explains the reductions in testicular weights and sperm counts (Fig. 4e,f). Consistently with a reduced number of meiotic and post-meiotic germ cells, the sizes of seminiferous tubules in *H3f4^H3.1/-^* testes were significantly reduced while the frequencies of stages of the seminiferous epithelium cycle were unaltered between genotypes (Supplementary Fig. 8e,f).

### Meiotic arrest in pachytene spermatocytes expressing H3.1 instead of H3.4

To investigate the cause underlying the reduction of pachytene and diplotene *H3f4^H3.1/-^* spermatocytes, we first performed immunofluorescence experiments on paraffin-embedded testicular sections for cleaved PARP1 (clPARP1), a marker of cells undergoing apoptosis^65^. Indeed, we observed elevated levels of clPARP1 signal in *H3f4^H3.1/-^* spermatocytes throughout the seminiferous tubule cycle, with pachytene cells at stages IV-IX predominantly affected, suggesting a problem in progressing through meiotic prophase (Fig. 5a,b).

**Figure 5.**
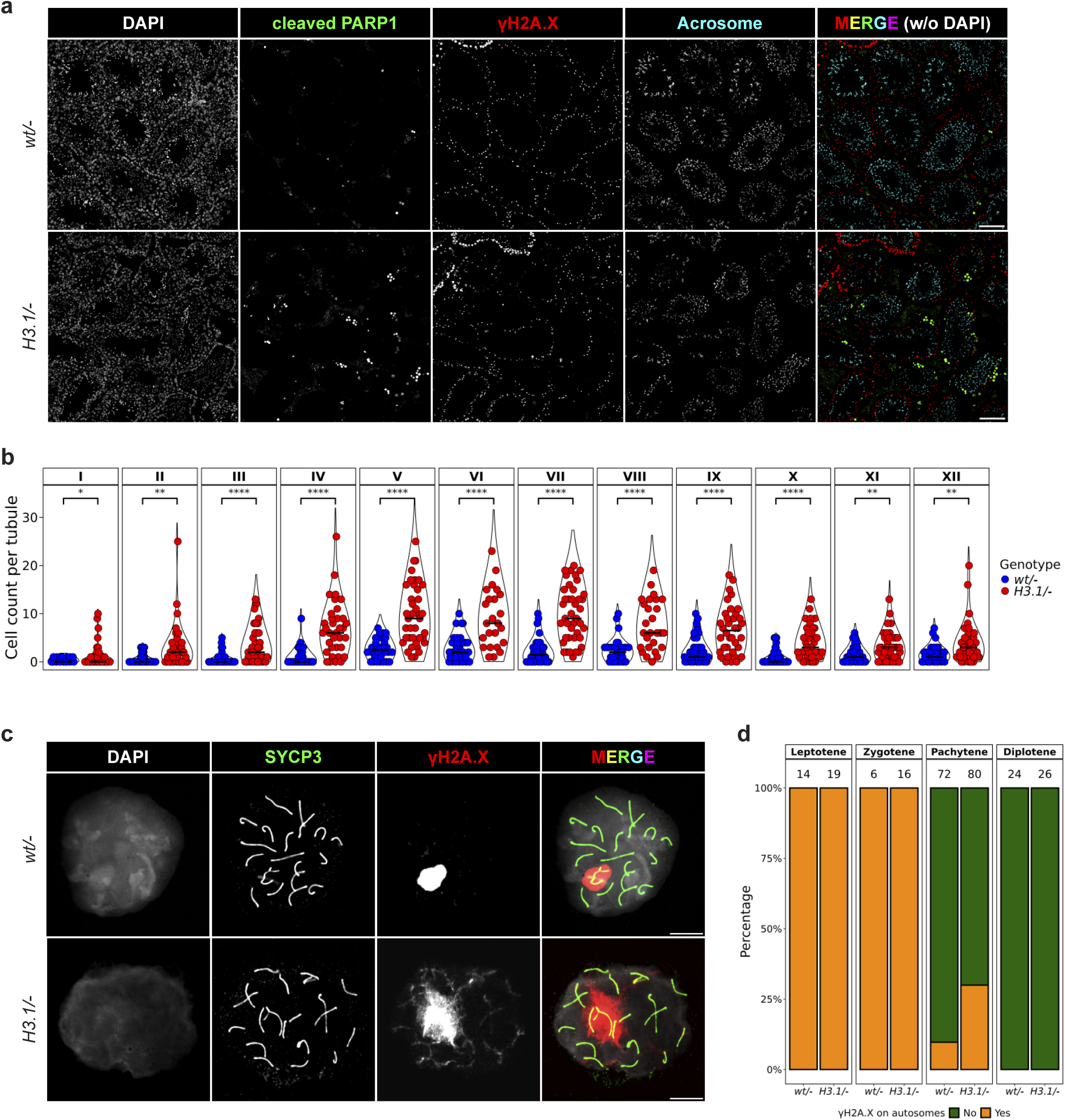
Impairment of meiotic progression in spermatocytes of *H3f4^H3.1/-^* substitution mice. **a**, Cross-section immunofluorescent images comparing cleaved PARP1 signal (green) between *H3f^fwt/-^* and *H3f4^H3.1/-^* mice. γH2A.X (red) and acrosome (turquoise) were used to mark spermatocytes and spermatids, respectively. As an overview, several tubules are shown. Scale bar: 50 µm. **b**, Quantification of testicular germ cells positive for cleaved PARP1 per stage comparing *H3f4^wt/-^* and *H3f4^H3.1/-^* mice. Staging was performed manually as described before^61^. Each dot represents the number of clPARP1-positive cells per tubule. Data from three biological replicates are pooled together. **c**, Meiotic spreads showing the normal and aberrant localization of γH2A.X signal on sex chromosomes and autosomes in pachytene spermatocytes isolated from *H3f4^wt/-^* and *H3f4^H3.1/-^* mice. DAPI is shown in grey, SYCP3 is shown in green, γH2A.X is shown in red. Scale bar: 10 μm. **d**, Bar plot showing the percentage of spermatocytes in prophase I with autosomal staining of γH2A.X, comparing *H3f4^wt/-^* and *H3f4^H3.1/-^* genotypes. A manual assessment of γH2A.X signal on autosomal chromosomes was performed, and the total number of cells analyzedd is indicated above each bar. Data from two biological replicates are pooled together.

To evaluate meiotic progression, we studied the processes of meiotic programmed double strand DNA break formation and repair, and of synapsis between homologous parental chromosomes by staining dried-down nuclear spreads of *H3f4^H3.1/-^* and control spermatocytes for phosphorylated H2A.X (γH2A.X) and SYCP3, a structural component of synaptonemal complexes^66–70^. Remarkably, while we observed comparable patterns of γH2A.X and SCP3 in leptotene and zygotene cells of *H3f4^wt/-^*and *H3f4^H3.1/-^* mice, a significant fraction of *H3f4^H3.1/-^*pachytene-like spermatocytes had elevated γH2A.X labelling of chromatin of homologous autosomes even though their synaptonemal complexes appeared fully synapsed (Fig. 5c, 5d, Supplementary Fig. 11). These data indicate that not all *H3f4^H3.1/-^* spermatocytes manage to process their meiotic DSBs appropriately before their entry into pachytene stage, consistent with the increased cell death of such *H3f4^H3.1/-^* spermatocytes (Fig. 4d). In summary, our data argue that chromatin largely composed of H3.1-containing nucleosomes is less proficient in meiotic DSB repair than H3.4-nucleosome containing chromatin.

### Transcriptional upregulation of Polycomb targets upon H3.4 to H3.1 substitution in meiosis

Beyond the impairment of meiotic progression and associated cell death, we next investigated the possible impact of H3.1 substituted expression on transcriptional regulation in the surviving cells. Therefore, we performed RNA-seq on ScLZ, ScPD, rSt and elSt populations FACS-isolated from *H3f4^H3.1/-^, H3f4^wt/-^* and *H3f4^wt/wt^* males. PCA plot showed a clear separation of cell types, recapitulating the developmental trajectory (Supplementary Fig. 12a). In addition, *H3f4^H3.1/-^* elongating spermatids were well separated from *H3f4^wt/-^* and *H3f4^wt/wt^* elSts in the first dimension suggesting a major change in their gene expression upon H3.4 replacement (Supplementary Fig. 12a). When analyzing the possible transcriptional changes in ScLZ, ScPD and rSt cells preceding spermatid elongation, we observed up to 159 genes up-regulated in these cells, while only a few genes were down-regulated (Fig. 6a and Supplementary Table 4). Interestingly, these three cell types shared several commonly upregulated genes (Fig. 6b), including transcription factors such as *Six2, Foxg1, Nkx6-1*, and a subset of *Hox* genes, as well as genes from the Vomeronasal receptor cluster (Supplementary Fig. 12b).

**Figure 6.**
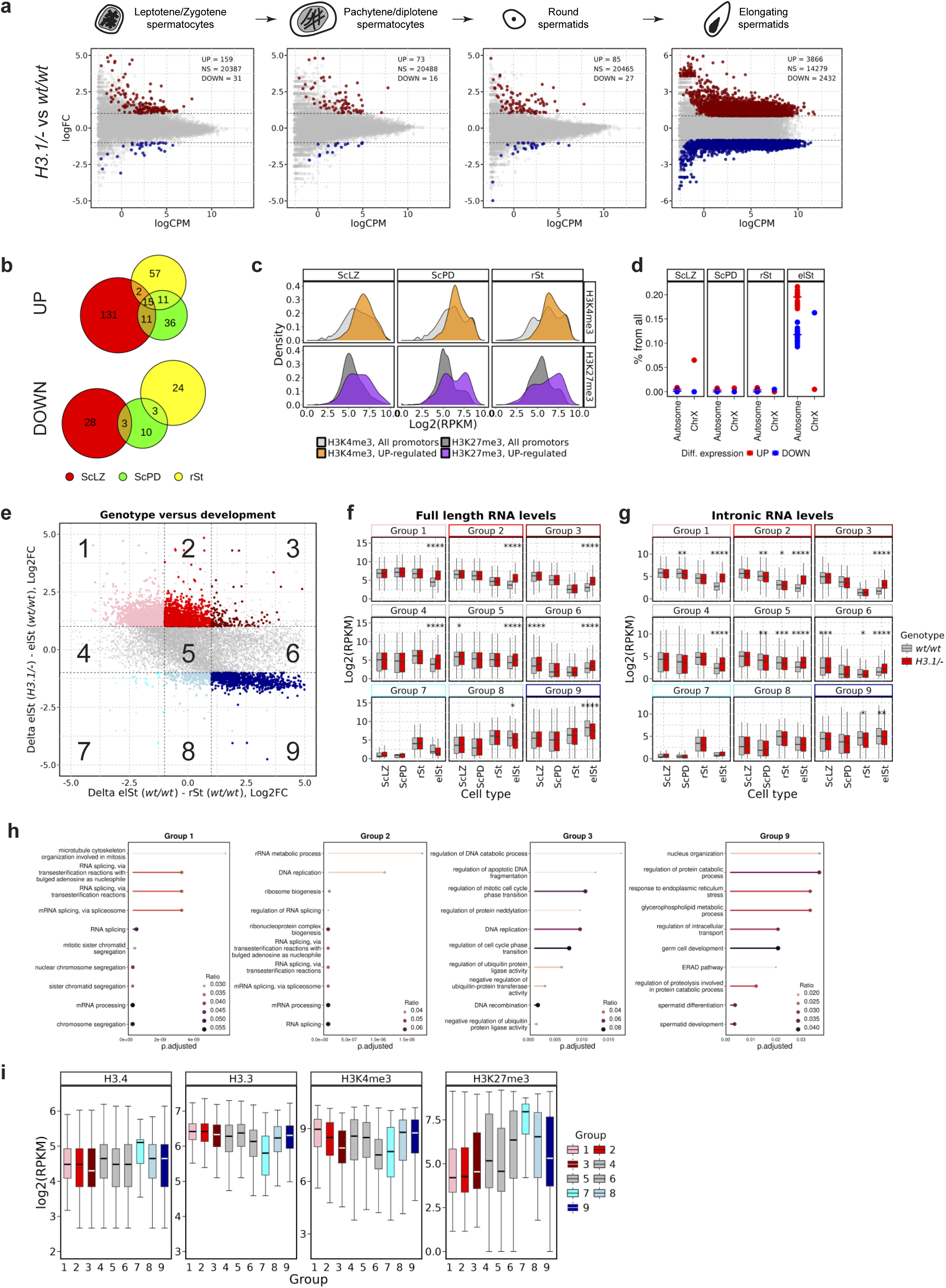
Differential gene expression in spermatocytes and spermatids of *H3f4^H3.1/-^* substitution mice. **a**, Scatter plots showing gene expression log2 fold changes in FACS-isolated ScLZ, ScPD, rSt and elSt in mutant (*H3f4^H3.1/-^*) cells plotted against average gene expression levels in control (*H3f4^wt/wt^*) cells. Data for full-length genic counts is shown. The numbers of significantly upregulated (red) or downregulated (blue) and non-significant (grey) genes are displayed on the top right. EdgeR parameters: min. logFC = 1, max. FDR = 0.05. **b**, Euler diagrams showing the overlap between DEGs in ScLZ, ScPD and rSt cells (panel a). **c**, Density plots showing the distribution of H3K4me3 and H3K27me3 (±1 kb around TSS) for the promoters of upregulated genes (coloured fill) and for promoters of all genes (grey fill). **d**, Dot plot showing the percentage distributions of UP- and DOWN-regulated genes per chromosome in ScLZ, ScPD, rSt and elSt cells. The bars represent the mean values for the selected DEGs. **e**, Scatter plots comparing two contrasts in CGI type promoter genes: genotype and mutation. The x axis represents the log2 fold change (FC) in gene expression during the transition from rSt to elSt in wild type (*H3f4^wt/wt^*) cells (wild-type elSt versus wild-type rSt, max. FDR = 0.05). The y axis represents the log2FC in gene expression in elSt cells caused by the *H3f4^H3.1/-^* mutation (*H3f4^H3.1/-^*versus *H3f4^wt/wt^*, max. FDR = 0.05). Genes UP- and DOWN-regulated in *H3f4^H3.1/-^* mutant cells are highlighted in shades of red and blue, respectively. The DEGs are separated into nine groups, based on the |log2FC| = 1 cutoff in both contrasts (dashed lines). To be included in groups 1-3 and 7-9, only the DEGs were considered. **f**-**g**, Boxplots showing absolute quantification of gene expression in groups 1-9 defined in panel e, comparing wild type (*H3f4^wt/wt^*) and mutant (*H3f4^H3.1/-^*) cells. Panel F shows full-length genic counts, panel g shows intronic counts. The colour code of groups is according to panel e. Paired t-test was performed between the indicated genotypes. * P < 0.05, ** P ≤ 0.01, *** ≤ 0.001, not significant changes are not displayed. **h**, GO term search (GO: Biological Process) for genes in groups 1, 2, 3 and 9 (panel e). **i**, Boxplots displaying wild type rSt ChIP-seq profiles of different chromatin signatures (H3.4^MYC^, H3.3, H3K4me3 and H3K27me3 counted in ±1 kb around TSS) in groups 1-9 defined in panel E. The colour code of groups is according to panel E.

To understand such shared mode of mis-regulation, we investigated chromatin signatures at promoters of the up-regulated genes (±1 kb around TSS) in wild type cells using publicly available ChIP-seq data for H3K4me3 and H3K27me3^52,56^. In all three cell types, we observed an overrepresentation of up-regulated genes among genes marked by H3K27me3, with a majority being bivalently marked by H3K4me3 as well (Fig. 6c, Supplementary Fig. 12b, 12c). Hence, these data suggest that H3.4-containing nucleosomes promote PRC2-mediated repression in male germ cells.

### The *H3f4^H3.1/-^* elongating spermatids transcriptionally resemble the preceding cellular stages with downregulated genes required for sperm maturation

Next, we focused our attention on *H3f4^H3.1/-^* elSts, in which we observed dramatic up- and down-regulation of 3866 and 2432 genes, respectively, compared to *H3f4^wt/wt^* control cells (Fig. 6a, Supplementary Fig. 12a and Supplementary Table 4). Interestingly, barely any X-linked genes were aberrantly upregulated (Fig. 6d), indicating that MSCI and PMSC-mediated gene repression mechanisms were not impaired. Given the prominent role of H3.3-nucleosomes in reprogramming chromatin of sex chromosomes during meiosis (Fig. 3, Supplementary Fig. 6, 7)^46,56,57^, the observed mis-regulation at autosomes only may point to a direct effect of H3.4 substitution by H3.1 on transcription during spermatid elongation.

To better understand the phenomenon, we related the aberrant transcriptional changes observed in *H3f4^H3.1/-^* versus control elSt cells to changes in gene expression that normally occur during the development of rSts into elSt in control mice. Given the differences in H3.4 and H3.3 occupancies at promoters of CGI-type and nonCGI-type genes (Fig. 3, Supplementary Fig. 6), we analyzed these two gene classes separately (Figure 6e, Supplementary Fig. 12d). For both classes, we observed a negative correlation between developmentally programmed expression changes versus changes aberrantly induced in *H3f4^H3.1/-^* elSts, suggesting a defect in spermatid differentiation. Specifically, genes in group 1 that are normally downregulated upon differentiation of rSt into elSt were up-regulated in *H3f4^H3.1/-^* elSts (Fig. 6e, 6f, Supplementary Fig. 12d, 12e). Analysis of intronic reads showed that such genes failed to downregulate their transcription (Fig. 6g, Supplementary Fig. 12f). A similar response was measured for groups 2 & 3 genes that are normally highly expressed in spermatocytes and become downregulated in spermatids. These genes were even aberrantly upregulated in *H3f4^H3.1/-^*elSts (Fig. 6e-g; Supplementary Fig. 12d-f). Gene ontology analysis indicated their involvement in housekeeping functions and meiosis (Fig. 6h), and in spermatid functions (Supplementary Fig. 12g). For genes belonging to groups 8 and 9 that normally sustain or even further increase their expression during nuclear elongation, we observed a down-regulation which likely abrogates at least in part spermatid differentiation, given their roles in spermatid differentiation and sperm function (Fig. 6e-6h; Supplementary Fig. 12d-g). Together, these data demonstrate a dual impact of H3.1 expression at the expense of H3.4, preventing wide-spread changes in transcriptional regulation and programs in elSts.

Finally, we analyzed chromatin signatures of the promoters of differentially expressed genes (±1 kb around TSS) as measured in wild type rSt cells, using ChIP-seq data of H3.4^MYC^ and H3.3 (this study) and public H3K4me3 and H3K27me3^56^. In CGI-type promoter genes, transcriptional upregulation in *H3f4^H3.1/-^*mutants mostly affected genes normally enriched in H3K4me3, the H3.3 variant and being devoid of repressive H3H27me3 (Fig. 6i). Concurrently, downregulated genes are normally marked with H3K27me3 in rSt (Fig. 6i). For non-CGI type promoter genes, the upregulated genes in groups 1-3 have normally higher H3.3 occupancy than unchanged or down-regulated genes, suggestive for the occurrence of higher turn-over of nucleosomes at these genes (Supplementary Fig. 12h). Furthermore, while H3K27me3 levels are indifferent among the groups of non-CGI genes, H3K4me3 levels are normally higher at genes that either failed to become upregulated in *H3f4^H3.1/-^* elongating spermatids. In summary, upon the *H3f4^H3.1/-^* substitution, major transcriptional changes occur at the stage of spermatid elongation. The *H3f4^H3.1/-^*elSt cells have a partial transcriptional profile of the preceding stage due to a failure to downregulate genes that have been active during meiosis or in rSt, or even due to an aberrant reactivation of such genes. At the same time, we measured reduced activation of genes required for spermatid differentiation that normally become highly expressed in elSt cells.

### Amino acids V24 and H42 are most important for the function of H3.4

Finally, we aimed to understand which of the H3.4-specific amino acids are most important to its function. Hence, we generated three additional *H3f4* substitution mouse lines, in which only one of the H3.4-specific amino acids was converted to that of H3.1 (*H3f4^V24A^, H3f4^H42R^* and *H3f4^S98A^*, Fig. 7a and Supplementary Fig.13a,b). As observed for the triple *H3f4^H3.1^* substitution allele (Fig. 4a), none of the single substitution alleles affected testis weights nor breeding efficiencies, if expressed homozygously (Fig. 7b-d). However, when combined with the *H3f4* deficiency allele (*H3f4^-^*), expression of the *H3f4^V24A^* and *H3f4^H42R^* substitutions resulted in decreased testis weights and sperm counts, compared to heterozygous control animals, while the *H3f4^S98A^*mutation did not impair spermatogenesis (Fig. 7b-d). Histological examination of testis sections confirmed the impairment of spermatogenesis, with *H3f4^H42R/-^*mice having the most pronounced effect (Fig. 7e).

**Figure 7.**
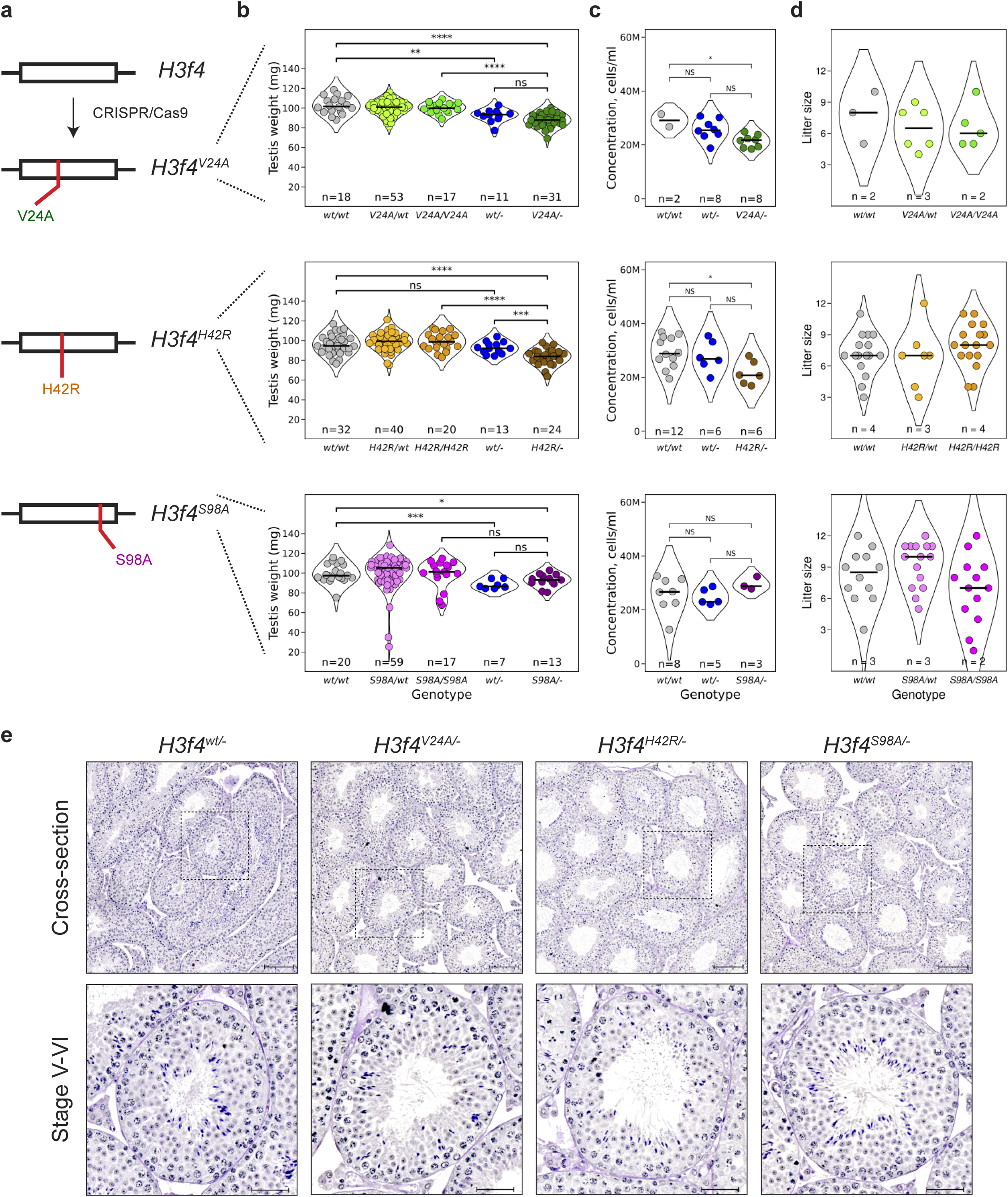
H3.4^V24^ and H3.4^H42^ amino acids contribute to spermatogenesis efficiency. **a**, Experimental design of the creation of *H3f4* gene alleles bearing H3.1-specific substitutions mutations (*H3f4^V24A^, H3f4^H42R^* and *H3f4^S98A^*) using CRISPR/Cas9 gene editing. Sequences of sgRNAs and repair templates are indicated in Supplementary Table 5 and 6, respectively. **b**, Violin plots showing the comparison of testis weights (mg) between *H3f4^wt/wt^, H3f4^mut/wt^, H3f4^mut/mut^, H3f4^wt/-^* and *H3f4^mut/-^* mice, where *H3f4^mut^* represents the *H3f4* allele bearing a H3.1-specific substitutions mutation (*H3f4^V24A^, H3f4^H42R^* or *H3f4^S98A^*). Each dot represents the weight mean of two testes taken from each mouse. The bars represent the median value for each genotype. The number of individual measurements (n) is indicated for each genotype. Paired t-test was performed between the indicated genotypes. * P < 0.05, ** P ≤ 0.01, *** P ≤ 0.001, NS – not significant. **c**, Estimation of sperm counts in *H3f4^wt/wt^, H3f4^wt/-^* and *H3f4^mut/-^* mice. Each dot represents sperm counts from a separate male. The bars represent the median value for each genotype. The number of individual measurements (n) is indicated for each genotype. Statistical testing was performed as described for panel b. **d**, Comparison of breeding efficiency in *H3f4^wt/wt^, H3f4^mut/wt^* and *H3f4^mut/mut^* mice. Each dot represents the number of pups born per litter; the bars represent the median value for each genotype. . The number of males (n) used in breeding tests are indicated for each genotype. Statistical testing was performed as described for panel b. **e**, PAS-haematoxylin staining of cross-section of testes in *H3f4^wt/-^, H3f4^V24A/-^, H3f4^H42R/-^*, and *H3f4^S98A/-^* mice. Scale bars: 100 µm (cross-section overview) and 50 µm (Stage V-VI tubules).

To investigate the impact of individual amino acid substitutions on gene expression, we performed bulk RNA-seq analyses on FACS-isolated rSt and elSt populations from *H3f4^V24A/-^* and *H3f4^H42R/-^*males. While principal component analyses revealed a clear separation of the rSt and elSt populations, it did not discriminate between genotypes (Supplementary Fig. 13c). Direct RNA-seq analyses showed a more pronounced effect of the *H3f4^H42R^* substitution than the *H3f4^V24A^* (Supplementary Fig. 13d), particularly in rSts. Comparison of differentially expressed genes between the *H3f4^H3.1^, H3f4^V24A^* and *H3f4^H42R^* datasets showed a small yet consistent overlap in rSts (Supplementary Fig. 13e), including homeobox genes *Hoxa5, Hoxa6, Six2* and genes from the Vomeronasal receptor cluster. Hence, the individual *H3f4^V24A^* and *H3f4^H42R^* H3.4 substitutions alter gene expression patterns slightly and affect testis weights and sperm counts mildly. Their combination, however, triggers significant responses during meiosis and spermatid elongation, ultimately impairing spermatogenesis. Therefore, we show that the A24V and R42H amino acids substitutions improved the function of H3.4 for male germ cell development.

## Discussion

Through synteny analyses, we identified that among eutherians with analyzable genomes all contain a small *Hist3* gene cluster harboring the *H3-4* locus and few *H2A* and *H2B* genes. Even the genome of elephant, being the eutherian mammal evolutionarily most distantly related to mouse and human, harbors a truncated cluster orthologous to *Hist3*. Moreover, our analyses suggest that *Hist3* cluster orthologs likely originate from an ancestral larger “canonical-like” histone cluster, of which a derivative currently exists in marsupial mammals that contains a *H3.2* gene, genes for all other core histones, CENPA and the linker histone H1.4. Hence, the *Hist3* cluster likely arose within the eutherian lineage after separation from the marsupial lineage over 100 million years ago.

Since then, *H3f4* orthologous genes accumulated more synonymous and non-synonymous mutations than canonical RC *H3.1* and *H3.2* genes situated in large histone clusters and RI *H3.3* genes individually located in the genome. Moreover, multiple acquired substitutions vary between different eutherian lineages suggesting that *H3f4* orthologs are subjected to reduced stabilizing selection compared to canonical RC genes. Given their expression in the germ line, eutherian *H3f4* orthologs genes may therefore been under directional selection, promoting spermatogenesis and/or embryonic development.

In mice, *H3f4* is exceedingly highly expressed in spermatogonia, driving H3.4 deposition already in undifferentiated spermatogonia, and causing an efficient and progressive accumulation of H3.4 throughout the entire genome during spermatogonial amplification until the entry into meiosis, at the expense of the other RC H3.1/H3.2 histones. As we demonstrated with a tagged H3.4^MYC^ allele and alike RC H3.1/H3.2 in somatic cells, H3.4 is successively replaced by H3.3 variant proteins at TSS and in genes following transcription-coupled nucleosome turn-over in proliferating spermatogonia. Upon entry into meiosis, H3.4-baring nucleosomes become further progressively exchanged by H3.3-nucleosomes within these regions as shown before^56^ as well as along the entire sex chromosomes, the later due to MSCI-related chromatin remodeling^46,54,57^.

By generating transgenic animals in which we substituted amino acids that are variant between H3.4 and H3.1, we provide evidence that *H3f4* has likely been under evolutionary directional selection. Our data show that particularly the combination of H3.4^V24^, H3.4^H42^ and H3.4^S98^ substitutions majorly promotes spermatogenesis, enhancing meiotic double strand break repair, gene regulation and germ cell viability. Considering the positive yet limited impact of single amino acid substitutions on germ cell development, we conclude that the H3.4^V24^ and H3.4^H42^ substitutions synergize in some way in improving spermatogenesis. Surprisingly, the positive impact of H3.4 over H3.1 sequence was noticeable only under reduced gene dosage conditions of RC H3 histones, even though we did not detect differences in total H3 levels by Western blot analyses between homozygous or hemizygous expression conditions of H3.4 or H3.1. Our data reveal that progression through meiotic prophase as well as nuclear elongation and condensation are particularly sensitive to H3.4 specific sequence acquisitions, while plain heterozygosity for H3.4 induced only limited cell death. The impact of H3.4^S98A^ is, however, neutral. Interestingly, the ancestral *H3.2* gene in marsupials harbors S98. Moreover, direct DNA sequence comparisons revealed high variability between eutherian and marsupial species as well as among eutherians suggesting reduced selective pressure on the DNA sequence encoding this residue, which is in line with the neutral impact of H3.4^S98^ versus H3.4^A98^ on spermatogenesis.

H3.4^H3.1/-^ males displayed an ∼20% reduction of spermatocytes prior to stage V of the seminiferous epithelium which coincides with persistent labeling of meiotic chromatin by phosphorylated H2A.X on autosomes in pachytene-like spermatocytes. These data point to a defect in repair of meiotically programmed DNA double strand breaks (DSBs), followed by activation of the synapsis checkpoint and cell death^71–73^. Nonetheless, we also observed an increased percentage of spermatocytes marked by cleaved PARP1 at following stages, suggesting persistent or newly formed DNA damage triggering cell death at later time points. Together, these data raise the intriguing possibility that the identity of RC H3 histones at meiotic recombination sites may regulate the formation, processing or repair of DSBs. For example, H3.4 may modulate tri-methylation of H3K4 and H3K36 by PRDM9, thereby controlling efficacy of SPO11-mediated DSB formation^74–76^. Alternatively, H3.4 and H3.1 may differentially regulate possible nucleosome turnover occurring at recombination sites during DSB repair.

Expression of *H3.4^H3.1^* affects gene expression only moderately during spermatogonial proliferation, meiosis and round spermatid differentiation. We predominantly observed preferential upregulation of genes normally bivalently marked by PRC2-mediated H3K27me3 and H3K4me3. These data are consistent with residue H3.4^V24^ being previously shown to promote recognition of tri-methylated H3.4K27 by PHF1 and PHF19, two accessory components of PRC2 complex, stabilizing its chromatin binding and promoting H3K27me3^44,51,77^. Indeed, we observed a clear overlap among genes upregulated in round spermatids isolated from *H3f4^V24A/-^* and *H3.4^H3.1/-^* mice. Many of these genes are normally marked by H3K27me3, supporting an *in vivo* role for H3.4 in promoting PRC2-mediated gene repression.

Surprisingly, we observed thousands of mis-regulated genes during spermatid elongation. On the one hand, our data points to a failure in the timely downregulation or even an aberrant transcriptional activation of genes normally expressed in round spermatids. On the other hand, *H3.4^H3.1^*-expressing elongating spermatids show reduced activation of spermatid differentiation genes normally upregulated during spermatid elongation. The data indicate that particularly elongating spermatids undergoing nucleosome replacement by transition proteins and protamines are sensitive to possible transcriptional regulatory functions of H3.4.

All eutherian *H3f4* orthologs encode for C96 which is absent in marsupial “*H3.2* orthologs”. Hence, C96 represents an ancient substitution mutation acquired early by the ancestral *H3f4* ortholog, like V24. Intriguingly, all RC H3.1 histones, specific to eutherians, encode also for C96, while H3.2 genes, expressed in all eukaryotes including eutherians and marsupials, encode for S96^78^. C96 has been proposed to regulate chromatin condensation by possibly forming intermolecular disulfide bonds between neighboring nucleosomes or between nucleosomes and other chromatin proteins^78^. The significance of C96 in H3.1 and H3.4 proteins for chromatin biology and spermatogenesis remains to be explored.

In human and other eutherians, various substitutions other than V24, C96 and S98, became fixed in *H3f4* orthologs. For human, chimpanzee and bonobo, the M71 and V111 residues regulate stability of nucleosomes^48,49^ and interactions with histone chaperones *in vitro*^50^, analogous to residue H42 in mice^42^. The *in vivo* function of M71 and V111 in regulating spermatogenesis, however, remains unknown. This is particularly relevant given that spermatogenesis in humans is less efficient than in rodents. Likewise, it will be important to assess the *in vivo* relevance of the variant Q134 and F54 residues encoded by genomes of some primate and bovine species, respectively.

The acquired role of *H3f4* orthologs in germ cells is not only determined by their conserved amino acid substitutions but also by their levels of expression relative to those of canonical H3.1 and H3.2 histones. Contrasting to mouse and human in which *H3f4* / *H3-4* is critical to spermatogenesis^42,47^, *H3f4* orthologs underwent loss-of-function mutations in some other species, which can be linked to different reproductive strategies. For example, *H3-4* is truncated in gorilla but is retained in most other primates with annotated genomes. The social reproductive system of gorilla populations is polygynous, which has been associated with reduced sperm competition and reduced testis to body ratios, compared to other primates^79^ and to relaxed selection for hundreds of genes implicated in germ cell development^80^. We also observed pseudogenization or even a complete loss of *H3f4* in four New World monkey species, although here the link to reproductive strategy is unclear. More examples exist in *Carnivora*, where *H3f4* encodes for a functional protein only in a subset of species. In *Felidae*, we identified a truncation of 13 nucleotides within the coding sequence of *H3f4*, resulting in a frameshift mutation (Supplementary Fig. 1c). Similarly, in *Canidae*, another truncation of 25 nucleotides was found in the open reading frame of *H3f4* of several dog breeds and Dingo (Supplementary Fig. 1c). Given that these variants were measured in several independent genome assemblies of evolutionarily close species, their presence is unlikely to be due to sequencing artifacts. The reason why cats and dogs have truncations in *H3-4* is enigmatic. In *Felidae*, polygyny is common and the mean testis to body weight in *Felidae* is among the lowest among all *Carnivora*^81^. *Canidae*, in turn, are characterized by social monogamy, yet with a high testis to body ratio, which has been linked to occasional intra-male competition. In any of these species, the absence of *H3f4* ortholog expression strongly advocates for a role of regular H3.1 and H3.2 proteins in germ cell development, underscoring the opportunistic impact of histone evolution in reproduction.

Several published studies^11,43^ and our own experiments (Supplementary Fig. 4) reveal the detrimental impact of expression of tagged core histones for germ cell development. Whereas such approach is commonly used in biochemical and cell biological experiments, our *in vivo* experiments raise awareness that too high expression levels are detrimental for development.

## Methods

### Phylogenetic analysis of H3 genes in the mammalian lineage

To search for *H3-4* orthologs in the mammalian lineage, we performed a TBLASTN search using the human H3.4 protein sequence as query against 110 mammalian genomes available at Ensembl (release 112). In each species, we looked for BLAST hits that encoded valine 24 (V24) and are located in the *HIST3* gene cluster^34^. If a hit was not annotated as a coding sequence, but otherwise seems to be correct, we manually translated the DNA sequences for this region. Species that possessed truncations, insertions or poor annotation of the putative *H3-4* DNA were not included in the analysis, or where possible, the annotation was adjusted in order to get the full-length protein. The results of this analysis are available in Supplementary Table 1. Pairwise divergence time trees were made using the TimeTree Database (http://www.timetree.org)^82^.

### Analysis of *H3-4* gene paralogues in the selected species

The Comparative Genomics functionality of the Ensembl website (release 112) was used to identify *H3-4* paralogues in human (GRCh38.p14), lemur (Mmur_3.0), mouse (GRCm38.p5), rabbit (OryCun2.0), cow (ARS-UCD1.3), elephant (Loxafr3.0), camel (CamDro2), wallaby (Meug_1.0), Tasmanian devil (mSarHar1.11), opossum (ASM229v1), koala (phaCin_unsw_v4.1) and common wombat (bare-nosed_wombat_genome_assembly). In species with an *H3-4* gene annotated in the genome (human, lemur, rabbit, cow, elephant, camel), *H3-4* was used as a query for the search for paralogues. In the species, where *H3-4* was not annotated but coding the sequence was identified in the genome (rabbit), canonical H3.1 was used as a query for the search for paralogues. For marsupial species, the H3.2S98 sequence was used as a query. The results of this analysis are available in Supplementary Table 3.

The information for Ensembl gene id, number of transcripts, transcript id, number of exons, cross reference to RefSeq and location was obtained using the Ensembl Perl api (release 112). For genes with more than one transcript, the transcript labelled with the flag “Ensembl Canonical” was initially selected for further analysis (Supplementary Table 3). Coding sequences and translations of coding sequences of all genes not annotated as pseudogenes were fetched as FASTA files via the Ensembl Perl api. In the case of rabbit, the genomic location, cDNA and protein sequences were added manually.

### Alignment of DNA and protein sequences

Alignments of DNA and protein sequences were performed with the msa R package^83^ with the default parameters (order = “input”). The first amino acid (methionine) was excluded from the analysis.

### Genome target selection and sgRNA design

The sequences of sgRNAs used in this study are available in Supplementary Table 5. The CRISPR design web tool was used to design sgRNAs. sgRNAs were chosen based on the proximity to the desired cut site (knock out) or insertion site (knock in) and the quality score was calculated as probability of on-target activity minus the sum of probability of off target cutting^84^. A minimum of three sgRNAs per experiment were compared for cutting efficiency *in vitro*.

### Generation of sgRNAs

sgRNAs were generated by one of the following three methods: (1) Oligonucleotides encoding for desired sgRNA (complementary, excluding the PAM and a BbsI restriction site added to the 5’) were annealed, ligated into the pX330-U6-Chimeric_BB-CBh-hSpCas9 plasmid^85^ and transformed into 10-beta Competent E. coli (C3019I, NEB). Individual clones were expanded and miniprep was performed using the NucleoBond Xtra Midi Kit (740410, Macherey Nagel). Positive clones were selected based on Sanger Sequencing analyzed using the CLC Main Workbench. A T7 promoter was added, and *in vitro* transcription was performed using the MEGAshortscript T7 Transcription Kit (AM1354, Thermo Fisher Scientific). sgRNAs were purified using the MEGAclear Transcription Clean-Up Kit (AM1908, Thermo Fisher Scientific). (2) The EnGen sgRNA Synthesis Kit (E3322S, NEB) was used according to the manufacturer’s instructions. In short, the EnGen sgRNA Template Oligo Designer (https://nebiocalculator.neb.com/#!/sgrna, NEB) was used to design oligos for *in vitro* transcription of sgRNAs. Resulting oligos were diluted to a final concentration of 1μM in nuclease-free H2O and mixed with EnGen 2X sgRNA Reaction Mix and EnGen sgRNA Enzyme Mix followed by incubation at 37°C for 2h. sgRNAs were purified using the RNA Clean & Concentrator-25 Kit (R1017, Zymo Research). (3) The Alt-R CRISPR-Cas9 System (IDT) following the manufacturer’s instructions was used. In short, on day of injection, Alt-R CRISPR-Cas9 crRNA and Alt-R CRISPR-Cas9 tracrRNA were mixed, heated to 95°C and left to cool down to RT to anneal.

### *in vitro* cleavage assay to test activity of sgRNAs

All sgRNAs were tested *in vitro* for cutting activity following guidelines by NEB (https://www.neb.com/products/m0386-cas9-nuclease-s-pyogenes) with minor adaptations. In short, sgRNAs were mixed with Cas9 protein (M0386S, NEB) and preincubated at 25°C for 10min. A linearized pGEM-T plasmid (A3600, Promega) containing the *H3f4* sequence +/- 200bp was added and incubated at 37°C for 60min. Resulting fragments were analysed on 2% agarose gels and cutting efficiency was scored based on the amount of uncut product left.

### Homology templates used to generate knock-in mouse models

The sequences of homology repair templates are available in Supplementary Table 6. (1) For homology-directed repair of <100bp, Ultramer DNA Oligonucleotides (IDT) complementary to the strand bound by the sgRNA (Supplementary Table 5) were used. To prevent cutting of the homology template and the newly generated allele, a part of the PAM sequence was altered and/or mismatches close to the PAM sequence were introduced. (2) To generate the *H3f4^H3.1^* allele, a plasmid was designed. The *H3f4* sequence plus 1kb up- and downstream were cloned into the pJET1.2/blunt plasmid (K1231, Thermo Fisher Scientific). The *H3f4* sequence was mutated using the Q5 Site-Directed Mutagenesis Kit (E0554S, NEB) to encode for a H3.1 protein. The resulting plasmid was amplified in 10-beta Competent *E. coli* and purified using the Plasmid Plus Maxi Kit (12963, QIAGEN).

### Preparation of injection mix

sgRNAs were mixed with Cas9 Nuclease (Cas9-TOO-50, Labomics) in IDTE buffer (11-01-02-02, IDT) and incubated at 37°C for 15min. Cas9 mRNA (CAS9MRNA, Sigma) and, pending on whether a knock in was made, the homology template was added. The mix was prepared freshly on the day of injection.

### Mouse husbandry, embryo collection, CRISPR injection and embryo transfer

Animal housing, handling and procedures of mice conformed to the Swiss Animal Protection Ordinance (protocol numbers: 1235, 1236, 2670, 3183 and 3091) and are compliant with the FMI institutional guidelines. Mice were housed in Type II long cages containing aspen bedding on IVC racks in rooms with 15–20 air changes per hour and a 12h light/dark cycle. Temperature was maintained within 20–24 °C with a relative humidity within 45–65%. Food and water were provided ad libitum.

Female mice were super ovulated by injection of 5U PMSG (MSD) followed by 5U hCG (MSD) after 48h. Zygotes were generated by natural mating and harvested at the 1-cell stage. Injections were done into the male pronucleus. On average 25 injected embryos were transferred into each ampulla of pseudo pregnant B6CF1 female mice.

### Sequence validation of mouse models

Genomic DNA of potential founder animals was isolated from ear biopsies and PCR was performed using the Q5 High-Fidelity DNA Polymerase (M0491, NEB). For the *H3f4* knock out mouse model, the following primers were used: 5’-TCCAGAACTCAGGAAAACTATGCC-3’ and 5’-ACTGAGCAGATGCGTTTGGA-3’. To genotype knock-in mouse models, primers spanning the *H3f4* gene were used: 5’-GGCGGACGATTCAGGAAG-3’ and 5’-TCCTGAGGAGACAGGACACC-3’. To exclude random integration of the plasmid in the *H3f4^H3.1^* mouse model, a primer combination with one primer outside and one inside the 5’ homology arm was used: 5’-GGCGGACGATTCAGGAAG-3’ and 5’-ATTTGAGGCCAGAGGGAGTT-3’.

The resulting PCR fragments were ligated into the pJET1.2/blunt vector and amplified in 10- beta Competent *E. coli*. Plasmids were isolated using the NucleoSpin Plasmid Kit (740588, Macherey-Nagel), subjected to Sanger Sequencing and analyzed using the CLC Main Workbench.

### Genotyping of mutant *H3f4* alleles

To identify mutant *H3f4* alleles, the two following strategies were used. (1) Enzyme-based genotyping. Genotyping of mutant *H3f4* alleles (*H3f4^V24A^, H3f4^H42R^, H3f4^S98A^*and *H3f4^H3.1^*) is based on the presence or absence of restriction enzyme cutting site in the *H3f4* gene introduced by the homology repair template. The PCR product was incubated with the restriction enzyme and the restriction fragments were analyzed on agarose gel. (2) Non-enzyme-based genotyping. To genotype the *H3f4* knock-out allele and tagged *H3f4* alleles, the genotyping assay is based only on the size of the PCR product. The sequences of primers and the PCR protocols are indicated in Supplementary Tables 7 and 8, respectively.

### Mass spectrometry analysis of putative H3 bands

To perform the MS analysis on the putative histone bands, we performed TAU gel electrophoresis on acid-extracted histones and stained the gel with Coomassie Brilliant Blue (Supplementary Fig. 3d). The migration patterns of bands corresponding to H3.1, H3.2 and H3.3 was described by Shechter and colleagues^55^. An additional band (band 1) was observed in the testis sample. We excised the putative H3 bands band (band 1 and 2) from testis along with bands from ESCs (band 3 and 4) and digested them using trypsin. An EASY-nLC 1000 Liquid Chromatograph (Thermo Fisher Scientific) and an LTQ Orbitrap Velos hybrid mass spectrometer (Thermo Fisher Scientific) were used to separate peptides. Mascot 2.3 (MATRIX SCIENCE) was used for database search and Progenesis LC-MS (Nonlinear Dynamics) was used for label free quantification. Peptides were normalized using eight H3 peptides conserved among H3 proteins (residues 54-63, 64-69, 73-83, 117-128 and 130-134). The additional testis band (band 1) is 8-fold enriched for the H3.4 peptide compared to the H3.1/2/3 peptide. Conversely, the H3.4 peptide is only detected at very low levels in band 2 (H3.3) and is nearly completely absent in bands 3 and 4 (H3.1 and H3.3), respectively.

### Isolation of H3 isoforms and determination of their relative ratios using TAU gel electrophoresis

To perform tissue-wide profiling of H3 isoforms, small pieces of different tissues were incubated in tissue lysis buffer (15 mM Tris-HCl pH 7.5, 60 mM KCl, 11 mM CaCl2, 5 mM NaCl, 250 mM sucrose, 5 mM MgCl_2_, 0.5 mM DTT, 5 mM sodium butyrate, 0.3 % NP-40 (74385, Fluka) and 1 tablet of cOmplete, EDTA-free Protease Inhibitor Cocktail (000000011873580001, Sigma) and homogenized in a Dounce homogenizer. Nuclei were pelleted by centrifugation and resuspended in 0.2 M HCl, incubated for a minimum of 2h at 4°C, precipitated in trichloroacetic acid and resuspended in H_2_O. Histones from cells were isolated as described previously^55^ with minor modifications. Briefly, pelleted cells were incubated in hypotonic lysis buffer (10 mM Tris-Cl pH 8.0, 1 mM KCl, 1.5 mM MgCl2, 1 mM DTT, 1 tablet of cOmplete, EDTA-free Protease Inhibitor Cocktail) for 50min rotating at 4°C. Nuclei were pelleted, resuspended in 0.4N HCl and incubated rotating o/n at 4°C. Samples were centrifuged at full speed and histone containing supernatants were loaded onto Amicon Ultra-0.5 mL Centrifugal Filters (UFC500324, Millipore) to exchange HCl to H_2_O. Finally, an aliquot was taken and dried in a SpeedVac chamber. The pellet was dissolved in ice cold freshly prepared TAU gel sample buffer (8M urea, 5% acetic acid, 5% β-mercaptoethanol and 1% methyl green), incubated for 30 mins on ice and loaded on the TAU gel.

To perform profiling of H3 isoforms in different populations during spermatogenesis, FACS-isolated cells were collected in Eppendorf tubes in the following numbers: 50K for spermatogonia, 25K for spermatocytes, 100K for rSt and 1M for elSt and eSt populations. The proportions for spermatogonia, spermatocytes and rSt cells were chosen to compensate for the DNA content. elSt and eSt populations were collected 10 times more than rSt, in order to compensate the loss of histones during spermatid elongation^86^. After sorting, cells were pelleted in a swing bucket centrifuge at 300 g for 10 min and the volume of the sample was reduced to 10 µl. The sample was frozen in liquid nitrogen and stored at -80°C until further processing. The sample was thawed on ice and 50 µl of lysis buffer was added (with a final concentration of 10 mM HEPES pH 7.5, 1.5 mM MgCl_2_, 10 mM KCl, 0.05% NP-40, 0.05 mM DTT, 1x of cOmplete, EDTA-free Protease Inhibitor Cocktail (000000011873580001, Sigma), 1 mM PMSF (Roche 10837091001), and 10 mM sodium butyrate). Lysis was performed on ice for one hour and then HCl was added to a final concentration 0.2 M and incubated overnight. For the gel, a 10 µl aliquot was taken and dried in a SpeedVac chamber. Pellets were dissolved in TAU gel sample buffer, as described above. Histone samples were loaded on freshly polymerized 15% gels (15% 60%:0.4% acryl:bisacrylamide, 6M urea/5% acetic acid/0.37% Triton-X100). Proteins were run at 200V at 4°C until the bands of methyl green exit the gel (5% acetic acid was used as a running buffer). Following the electrophoresis, gels were incubated in TAU gel transfer buffer (0.5% acetic acid) for 15 min. Finally, proteins were transferred to the PVDF membrane for 20 min at 500 mA. After the transfer, a standard Western Blotting protocol was used. To visualize all H3 isoforms, we used the anti-H3 antibody that recognized the C-terminal epitope present in all H3 isoforms (Abcam ab1791, 1:10000, Supplementary Table 9). The ratio between the intensities of bands was used to determine the relative ratio of H3 isoforms.

### SDS-PAGE followed by Western Blotting

Histone extracts or whole cell pellets were resuspended in SDS sample buffer (50 mM Tris-HCl pH 6.8, 12.5 mM EDTA, 10% glycerol, 2% SDS, 1% β-mercaptoethanol, 0.02 % bromophenol blue), incubated for 10 min at 94°C and separated on a 15% polyacrylamide gels. Antibodies used in Western blot experiments are indicated in Supplementary Table 9.

### Fixation of tissues and preparation of microscopy sections

Preparation and analysis of histological samples of mouse testis and epididymal tissues was performed as described previously^64^. In brief, freshly isolated testis or epididymis samples were fixed in 5-10 ml of Bouin’s solution (Sigma HT10132) or in 4% PFA in PBS, for the histological and immunofluorescence experiments, respectively. Fixation was performed overnight on a rotating mixer at 4°C. After fixation, samples were washed twice with 70% ethanol. Fixed samples were de-hydrated through a graded series of ethanol solutions (2 × 70%, 80%, 2 × 96% and 3 × 100%) and xylene and finally embedded in paraffin using an automated tissue processing center (TPC 15 Duo, Medite) with standard settings. Sectioning was done at 3 µm thickness using the automatic microtome (HM355S, Thermo Fisher Scientific). Sections were mounted onto Superfrost Plus Adhesion Microscope Slides (J1800AMNZ, Thermo Fisher Scientific) and dried at 37°C overnight.

### Periodic Acid-Schiff and Haematoxylin staining of paraffin-embedded testicular sections

For histological examination, testis sections were deparaffinized by incubating in xylene solution (Sigma 534056) 2 times for 5 mins and then rehydrated in a series of decreasing concentrations of ethanol (2×100%, 95%, 70%, 3 min each) to deionized water. Rehydrated tissues were immersed in Periodic Acid Solution (Sigma, 395132) for 5 min at RT, rinsed several times in deionized water, immersed in Schiff’s Reagent (Sigma 3952016) for 15 min at RT and then rinsed in tap water for 5 min. Samples were counterstained with Mayer’s Haematoxylin Solution (MHS32, Sigma) for 2 mins and rinsed in tap water for 5 mins. Finally, samples were dehydrated in a series of increasing concentrations of ethanol (70%, 95%, 2×100%, 3 min each), cleared in xylenes solution (2 × 3min) and mounted with Permount^TM^ mounting media (ThermoFischer, SP15-100). mage acquisition was performed using motorized automated slide Scanner Zeiss Axioscan Z1 with 20X air objective and analyzed with ZEN blue software (version 2.3, Zeiss).

### Immunofluorescence imaging of paraffin-embedded testicular sections

Testis sections were deparaffinized by incubating in xylene solution (Sigma 534056) 2 times for 5 mins and then rehydrated in a series of decreasing concentrations of ethanol (2×100%, 95%, 70%, 3 min each) to deionized water. Antigen retrieval was performed for 20 min at 94°C in 10 mM sodium citrate buffer (pH 6.0) containing 0.05% Tween-20. Finally, the slides were washed in PBS. Around each sample, a circle was drawn using DAKO hydrophobic pen (Agilent S200230-2), in order to perform staining reactions in smaller volume. Blocking was performed in 30-50 µl of BPS containing 5% horse serum, 1% BSA and 0.05% Tween-20 for 1 hour. Primary antibodies (Supplementary Table 9) were diluted in blocking buffer, added to the samples, and incubated overnight at 4°C. After incubation with primary antibodies, samples were washed with PBS and incubated with secondary antibodies diluted in blocking buffer for 1 hr. Finally, sections were incubated 1:1000 DAPI solution (Sigma D9542) for 10 min and then washed PBS. Mounting was performed using VECTASHIELD Antifade Mounting Medium (H-1000).

Images were obtained using spinning disk confocal scanning unit Yokogawa CSU W1 Dual T2 with 40x/1.3 oil immersion objective. Image analysis was performed with Fiji software^87^ and using the automated image quantification pipeline described below.

### EdU labelling of replicating cells

Intraperitoneal injection of 10 mg/ml EdU solution (Thermo Fischer Scientific A10044) was performed in the proportion of 0.5 g of pure EdU per 10 g of mouse weight. 3 hours after injection, animals were sacrificed, followed by collection, PFA fixation and paraffin-embedding of testes by a standard protocol.

For imaging, testicular sections were prepared, rehydrated, and blocked by a standard protocol. Labelling reaction mix was prepared in the following proportions: 400 µl PBS, 5 µl 100 mM CuSO_4_, 0.5 µl Alexa Fluor 488 Azide (Thermo Fischer Scientific C10632) and 100 µl 500 mM ascorbic acid (added last). After 1 hour of blocking, 30-50 µl of labelling reaction solution was added to samples and incubated for 30 min at room temperature in dark chamber. After the completion of reaction, samples were washed several times with blocking buffer and processed into a standard immunofluorescence imaging protocol.

### Preparation of meiotic spreads

Meiotic spreads were prepared as described previously^88,89^. Prior to staining, slides were washed in PBS, in order to remove the excess of PFA.

### Whole-mount immunofluorescence of testicular tubules

After decapsulation of testes from adult mice, seminiferous tubules were gently detangled and washed 2-3 times with PBS, to remove the interstitial cells. Then, the tubules were fixed with 4% PFA in PBS for 1 hour at 4°C. After fixation, the tubules were washed 3 times in PBS and then 3 times in PBS containing 0.04% Tween 20 (PBST) in a cell strainer at RT, each wash for 10 mins. Then, the tubules were dehydrated through a graded series of 25%, 50%, 75% methanol containing PBST and 100% methanol at 4°C for 7 mins, respectively. The samples were stored at -80°C for later usage. At the time of observation, the samples were rehydrated through a graded series of 75%, 50%, 25% methanol containing PBST for 7 mins at 4°C and then washed 3 times with PBST for 10 mins each at RT. Rehydrated tubules were blocked in PBST containing 4% Normal Donkey Serum (Abcam ab7475) for 1 hour at RT. After blocking, the tubules were incubated with primary antibodies for 3 hours at RT. After this, the tubules were washed 3 times for 10 mins in blocking buffer followed by incubation with secondary antibodies for 2 hours at RT. Finally, tubules were incubated 1:1000 DAPI solution (Sigma D9542) for 10 mins and then washed 3 times with PBST for 10 mins. For imaging, tubules were oriented between a microscope slide and a coverslip separated by a 0.12 mm thick SecureSeal™ Imaging Spacers (Grace Bio Labs), with PBST used as a mounting media.

Images were obtained using spinning disk confocal scanning unit Yokogawa CSU W1 Dual T2 with 63x/1.3 oil immersion objective. Image analysis was performed with Fiji software^87^.

### Quantitative analysis of immunofluorescence data

The quantitative analysis of the immunofluorescence images is performed in five consecutive steps: (1) Projecting, stitching and binning of acquired image tiles; (2) Segmentation of testes tubules; (3) Extraction of testes tubules (4) Nuclei segmentation in extracted testes tubules and (5) Cell classification. In detail: *(1) Projecting, stitching and binning:* The raw input data consists of 3D stacks with 10% overlap in XY for all four acquired channels. These 3D tiles a first maximum intensity projected (MIP) along Z. Then the illumination correction is applied to the DAPI channel for which the illumination field is obtained by averaging all 3D stacks of the DAPI channel for a given acquisition, computing the maximum intensity projection along Z, and fitting a 2D Gaussian. The MIPs are stitched with the Fiji Image Stitching plugin^90^. The stitched image is average binned by a factor of 2 and saved. Additionally, the 8 times averaged binned version is saved for segmentation. *(2) Segmentation of testicular tubules:* Testes tubules were segmented on the 8 times average binned DAPI channel with a custom StarDist (v0.7.3)^91^ model. For the training four images where manually annotated with Napari (https://zenodo.org/records/8076424). The ground truth data consists of 14 training patches and 2 validation patches containing a total of 752 annotated tubules. The StarDist network uses default parameters except for n_rays = 128 and grid = (4, 4). It was trained for 400 epochs, with 100 steps per epoch, an initial learning rate of 0.0003 and a training batch size of 8. Training was performed on nVidia Tesla V100 GPU with 32GB of memory. *(3) Extraction of testicular tubules:* The StarDist model for tubule segmentation predicts a star-convex polygon centred on the tubules. The predicted polygons are upscaled from 8x binning to 2x binning and refined such that the label boundary is not cutting through nuclei. *(4) Nuclei segmentation in extracted testes tubules:* A second StarDist model was trained for the nuclei segmentation on the 2 times binned DAPI channel. The ground truth data was manually annotated with Napari and split into 12 training and 4 validation images. The StarDist network uses default parameters except for n_rays = 64 and gird = (1, 1). It was trained for 400 epochs, with 100 steps per epoch, an initial learning rate of 0.0003 and a training batch size of 4. Training was performed on an nVidia Tesla V100 GPU with 32GB of memory. *(5) Cell classification:* Segmented cell nuclei from tubules of stages of interest were classified using Ilastik software^92^.

### Fluorescence Activated Cell Sorting (FACS)

SgU, SgD, ScLZ and ScPD cells used for ultra-low native ChIP-seq experiments (ULI-NChIP-seq, Figure 3), we used a previously published protocol based on surface markers of spermatogonial populations (Supplementary Table 9)^64^.

To isolate ScPD and rSt cells used for conventional ChIP-seq experiments (Supplementary Figure 6) and ScLZ, ScPD, rSt and elSt cells for RNA-seq (Figure 6a), we used a previously published protocol based on dual staining of DNA with Hoechst 33342 and SYTO 16^89^.

### Sperm counting

After euthanizing a mouse, two cauda epididymis were isolated, with surrounding fat carefully removed, and squeezed in total of 1000 µl PBS. The suspension was filtered through 30 um mesh. From this filtrate, an aliquot of 50 µl was diluted in 1000 µl of water, to immobilize cells. From this solution, 10 µl were taken to manually count cells in a cell counter chamber.

### Antibodies

All primary and secondary antibodies used in this study are indicated in Supplementary Table 9.

### RNA sequencing

FACS-isolated populations of ScLZ, ScPD, rSt and elSt cells were pelleted at 2000g for 10 min at 4°C. Total RNA was extracted from cell pellets using the RNeasy Mini Kit (Qiagen 74104) according to manufacturer’s instructions. DNA removal was performed using on-column the RNase-Free DNase Set (Qiagen, 79254). RNA concentration was determined with Qubit RNA HS Assay Kit (Q32852). Quality of RNA was determined using Fragment Analyzer system. Libraries were prepared with the Illumina Stranded total RNA-seq protocol according to manufacturer’s instructions. Quality control of libraries was performed with Agilent 2100 Bioanalyzer System. After passing the QC, sequencing was performed on Illumina NovaSeq platform (50 bp, paired end).

### Chromatin Immunoprecipitation followed by sequencing (ChIP-seq)

To profile the H3 isoforms in early cell populations (SgU, SgD, ScLZ and ScPD), we followed the previously published ultra-low ChIP protocol^63^. Its adaptation to FACS-isolated spermatogonial populations with the detailed workflow was described in our previous study^64^. In brief, to profile the H3 isoforms, chromatin from 5K SgU or 10K SgD, ScLZ or ScPD cells were used with 0.5 µl of anti-H3.3 (Cosmo Bio CE-040B) or 0.5 µl anti-Myc (Abcam ab9132) antibodies per sample. ULI-ChIP-seq libraries were prepared from the eluted ChIP DNA using the NEBNext Ultra II DNA Library Prep Kit for Illumina (NEB E7645L), as described before^64^. Libraries were sequenced in 50-bp paired-end mode with Illumina NovaSeq 6000 using the manufacturer’s instructions.

ChIP-seq in post-replicative cell populations (ScPD and rSt) was performed as follows. Chromatin from FACS-isolated ScPD and rSt populations was prepared by incubating cells for 20 min on ice in lysis buffer (50 mM Tris-HCl (pH 8.0), 150 mM NaCl, 1% Triton X-100, 0.2% sodium deoxycholate, 5 mM CaCl_2_) containing 5 U MNase (Roche Nuclease S7 10107921001) per 1M cells. After incubation on ice, chromatin was digested for 10 min at 37°C. The MNAse reaction was stopped by adding EGTA to 50 mM final concentration. The IP reactions were performed overnight at 4°C on a rotator in the following proportions. To profile H3 isoforms, chromatin from 50K ScPD or 200K rSt was incubated with 5 µl of antibodies (anti-MYC: Abcam ab9132, anti-H3.3: Cosmo Bio CE-040B). To profile H3K4me3 and H3K27me3 in ScPD cells, chromatin from 1200K ScPD was incubated with 3 µl of antibodies (anti-H3K4me3: Millipore 17-614, anti-H3K27me3: Cell Signaling #9733). Per sample, 25 µl of 1:1 mix of protein A:protein G Dynabeads (Thermo Fisher Scientific, 10001D and 10003D) was washed once with 25 µl lysis dilution buffer (50 mM Tris-HCl (pH 8.0), 150 mM NaCl, 1% Triton X-100, 0.2% sodium deoxycholate, 50 mM EGTA), then added to chromatin-antibody mix and incubated on a rotator for 4 h at 4°C. The Chromatin-Antibody-Dynabeads complexes were washed twice with 200 µl of RIPA buffer (140 mM NaCl, 10 mM Tris-HCl (pH 8.0), 0.1% Sodium deoxycholate, 0.1% SDS, 1% Triton X-100, 1 mM EDTA), once with 200 µl high salt buffer (140 mM NaCl, 10 mM Tris-HCl (pH 8.0), 0.1% Sodium deoxycholate, 0.1% SDS, 1% Triton X-100, 1 mM EDTA), once with 200 µl LiCl wash buffer (250 mM LiCl, 0.5% NP40, 0.5% deoxycholate, 1 mM EDTA, 10 mM Tris-HCl, (pH 8.0)), and once with 200 µl TE buffer (10 mM Tris-HCl (pH 8.0), 1 mM EDTA). Finally, chromatin was eluted and digested for 1 h at 63°C in Elution buffer (10 mM Tris-HCl (pH 8.0), 1 mM EDTA, 0.1% SDS, 300 mM NaCl). DNA from eluted material was purified by 2X volumes of Ampure XP DNA purification beads (Beckman Coulter, A63881) according to manufacturer’s instructions and resuspended in 9 µl of nuclease-free water. The eluted ChIP DNA was used to prepare libraries using the NEB Ultra protocol according to the manufacturer’s instructions. The libraries were sequenced in a 50 pb single end mode on the Illumina HiSeq 2500 platform, according to the manufacturer’s instructions.

### Analysis of publicly available genomic data

ChIP-seq data profiling H3K4me3 and H3K27me3 in wild type SgD cells and H3K27me3 in wild type ScLZ and ScPD cells prepared with ULI-ChIP method were downloaded from public GEO database GSE214682^52^. ChIP-seq data profiling H3K4me3 and H3K27me3 in wild-type rSt cells were obtained from the GEO database GSE42629^56^. RNA-seq data from wild type SgU, SgD and ScLZ cells was downloaded from public GEO database GSE214682^52^. RNA-seq from wild-type ScLZ, ScPD, rSt, elSt and eSt populations were downloaded from the public GEO database GSE214316^53^.

### Computational analysis of ChIP-seq data

ChIP-seq data were processed with TrimGalore (version 0.6.2) to perform adaptor trimming and remove low quality reads with settings (-clip_R1 2 --stringency 3). The trimmed reads were aligned to the mouse genome build mm10 using STAR (version 2.7.10a^93^) with settings (--alignIntronMin 1 –alignIntronMax 1 --alignEndsType EndToEnd --alignMatesGapMax 1000 --outFilterMatchNminOverLread 0.85). The BAM files were de-duplicated using SAMtools (version 1.10) with standard settings and used for the downstream analysis.

To profile the occupancy of the H3 isoforms genome-wide, the mouse genome from the BSgenome.Mmusculus.UCSC.mm10 assembly was partitioned into 10 kb non overlapping tiles. Tiles overlapping with blacklisted regions^94^ were excluded from the downstream analysis. The tiles were classified as containing a TSS if an overlap with ±1 kb around genic TSS (TxDB.Mmusculus.UCSC.mm10.knownGene) was reported. Further, the uniquely mapped reads were counted on genomic regions with the qCount function (mapqMin = 225L) from the QuasR package^95^ and correction for reads originating from chrX was performed. log2CPM values were calculated for each sample and normalized within each ChIP experiment (i.e., libraries from different cell types but the same antibody were normalized to each other) using normalizeBetweenArrays function from the Limma package^96^ (method = “cyclicloess”, cyclic.method = “fast”).

To profile the H3 isoforms and the H3 modifications at gene promoters, genic TSS coordinates were extracted as described above and reads mapping to ±5 kb around the TSS were calculated with the qProfile function (binSize = 201L) and log2CPM values were calculated. To normalize these data, additionally, qProfile quantification (binSize = 201L) was performed on 10 kb genomic tiles that do not contain TSS and do not overlap with chrX. For these tiles, log2CPM values were calculated and tiles that commonly possess read counts below Q25 and above Q75 per ChIP were defined. These regions were added to the genic TSS, and normalization was performed on the merged data as described above. After normalization, non-TSS regions were removed, and downstream analysis was performed on normalized TSS data only. In Figure 3A and Figure S6A, clustering was done on rowSums values of normalized log2CPM qProfile data combined with RNA expression data. The resulting data frame was scaled and clustered using the functions hclust (method = “ward.D”) and dist (method = “euclidean”) from the stats R package (version 4.2.1). The order of rows in cluster was defined with Rtsne function (dims = 1) on aggregated qProfile data. For heatmap plotting, normalized data was visualized using the ComplexHeatmap R package^97^.

### Computational analysis of RNA-seq data

RNA-seq data were processed with TrimGalore (version 0.6.2) to perform adaptor trimming and remove low quality reads with settings (-clip_R1 2 --stringency 3). The trimmed reads were aligned to the mouse genome build mm10 using STAR (version 2.7.10a^93^) with settings (--alignIntronMin 1 –alignIntronMax 1 --alignEndsType EndToEnd --alignMatesGapMax 1000 --outFilterMatchNminOverLread 0.85). The BAM files were de-duplicated using SAMtools (version 1.10) with standard settings and used for the downstream analysis. Quantification of gene expression (from TxDb.Mmusculus.UCSC.mm10.knownGene Bioconductor annotation) was performed with QuasR R package^95^.

Genes with at least 1 count per million (CPM) in three biological replicates were used in downstream analysis. The EdgeR R package^98^ was used for statistical analysis of differential gene expression between the following comparison pairs: (1) *H3f4^wt/-^* and *H3f4^H3.1/-^*; (2) *H3f4^wt/-^* and *H3f4^V24A/-^*; (3) *H3f4^wt/-^* and *H3f4^H42R/-^*. CPM values were normalized using trimmed mean of M values (TMM) normalization using function calcNormFactors(method = “TMM”). Generalized Linear Model (GLM) was fit using genotypes as covariates. The following statistical significance cutoffs were used: |log2FC |> 1 and false discovery rate-adjusted (FDR) < 0.05. Gene Ontology (GO) search was performed using clusterProfiler R package^99^.

## Acknowledgements

We thank all Peters lab members for the critical reading of this manuscript and helpful input, and specifically Dr. Gaspa-Toneu, Dr. Gill and Dr. Ozonov for help with wet lab experiments and computational analysis of sequencing data, respectively. We thank Dr. Stadler for advice on ChIP-seq data analysis. We thank Dr. Plantard, Dr. Gelman and the facility for advanced imaging and microscopy (FAIM) for support with microscopy experiments. We thank Dr. Smallwood and the genomics facility for their help with NGS library preparations and sequencing. We thank Dr. van der Vlag and Dr. Heyting for sharing antibodies. This work supported by funding from the Novartis Research Foundation and the Swiss National Science Foundation (31003A_146293 and 31003A_172873).

## Author contributions

P.A.K., P.B., C.-Y.L. and A.H.F.M.P. designed experiments. P.A.K. and H.-R.H. performed evolutionary analysis of the *H3* genes. P.B., C.-Y.L. and J.-F.S. generated *H3f4* mouse models by CRISPR/Cas9 editing. H.K. performed FACS experiments. C.-Y.L. and P.A.K. performed TAU gel experiments. C.-Y.L. performed MS experiments. P.A.K., G.F. and S.C. performed genomic experiments. P.A.K. analyzed genomic data. P.A.K. performed imaging experiments. T.-O.B. developed the imaging quantification pipeline, with input from P.A.K. P.A.K., P.B., and A.H.F.M.P. wrote the manuscript, with input from all authors. A.H.F.M.P. conceived and supervised the overall project.

## Conflict of interest

The authors declare no conflict of interest.

## Notes

### Competing Interest Statement

The authors have declared no competing interest.

