## Supplementary Figures 1-13 and legends for "The eutherian-specific histone H3.4 promotes germ cell development and reproductive fitness"

### Supplementary Figures and Supplementary Table Legends

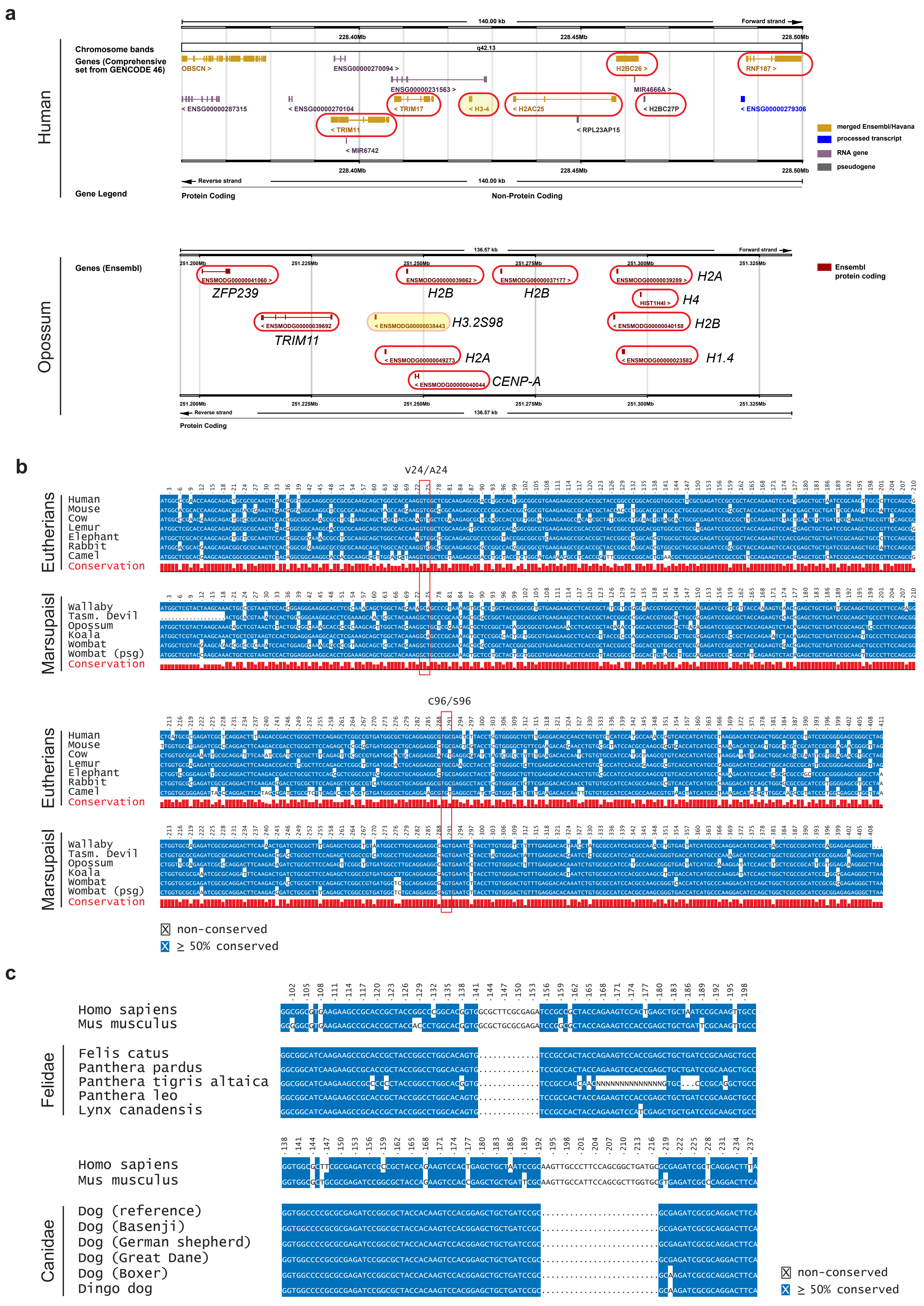
a
Human
Chromosome bands
Genes (Comprehensive set from GENCODE 46)
Gene Legend
140.00 kb
Forward strand
228.40Mb
228.45Mb
228.50Mb
q42.13
OBSCN >
ENSG00000270094 >
ENSG00000231563 >
H2BC26 >
RNF187 >
MIR4666A >
< ENSG00000287315
< ENSG00000270104
< TRIM17
< H3-4
< H2AC25
< H2BC27P
< ENSG00000279306
< TRIM11
< RPL23AP15
< MIR6742
Reverse strand
Protein Coding
Non-Protein Coding
merged Ensembl/Havana
processed transcript
RNA gene
pseudogene
Opossum
Genes (Ensembl)
136.57 kb
251.200Mb
251.225Mb
251.250Mb
251.275Mb
251.300Mb
251.325Mb
ENSMODG00000041060 >
ZFP239
ENSMODG00000039862 >
H2B
ENSMODG00000037177 >
H2B
ENSMODG00000039289 >
H2A
HIST1H4I >
H4
< ENSMODG00000039692
TRIM11
< ENSMODG00000038443
H3.2S98
< ENSMODG00000040158
H2B
< ENSMODG00000049273
H2A
< ENSMODG00000023582
H1.4
< ENSMODG00000040044
CENP-A
Ensembl protein coding
b
V24/A24
Eutherians
Human
Mouse
Cow
Lemur
Elephant
Rabbit
Camel
Conservation
Marsupials
Wallaby
Tasm. Devil
Opossum
Koala
Wombat
Wombat (psg)
Conservation
C96/S96
non-conserved
≥ 50% conserved
c
Homo sapiens
Mus musculus
Felidae
Felis catus
Panthera pardus
Panthera tigris altaica
Panthera leo
Lynx canadensis
Canidae
Dog (reference)
Dog (Basenji)
Dog (German shepherd)
Dog (Great Dane)
Dog (Boxer)
Dingo dog
non-conserved
≥ 50% conserved

**Supplementary Figure 1. Sequence and locus conservation of the H3 proteins in the selected mammalian species**

**a**, Ensembl genome browser snapshots of the *H3-4* locus and neighbouring genes in human (top) and opossum (bottom). Genes surrounded by red lines correspond to genes presented in Fig. 1B. For opossum, gene symbols have been manually added. Links to human and opossum genome browser views are:

https://may2024.archive.ensembl.org/Homo_sapiens/Location/View?db=core;r=1:228360000-228500000
https://may2024.archive.ensembl.org/Monodelphis_domestica/Location/View?r=2:251195710-251332276

**b**, Alignment of coding sequences of *H3-4* orthologs in selected placental and marsupial species. Codons for V24/A24 and C96/S96 are highlighted in red.

**c**, Alignment of parts of coding sequences of *H3-4* orthologs in species of *Felidae* (top) and *Canidae* (bottom) families. Human *H3-4* and mouse *H3f4* sequences are given as reference.

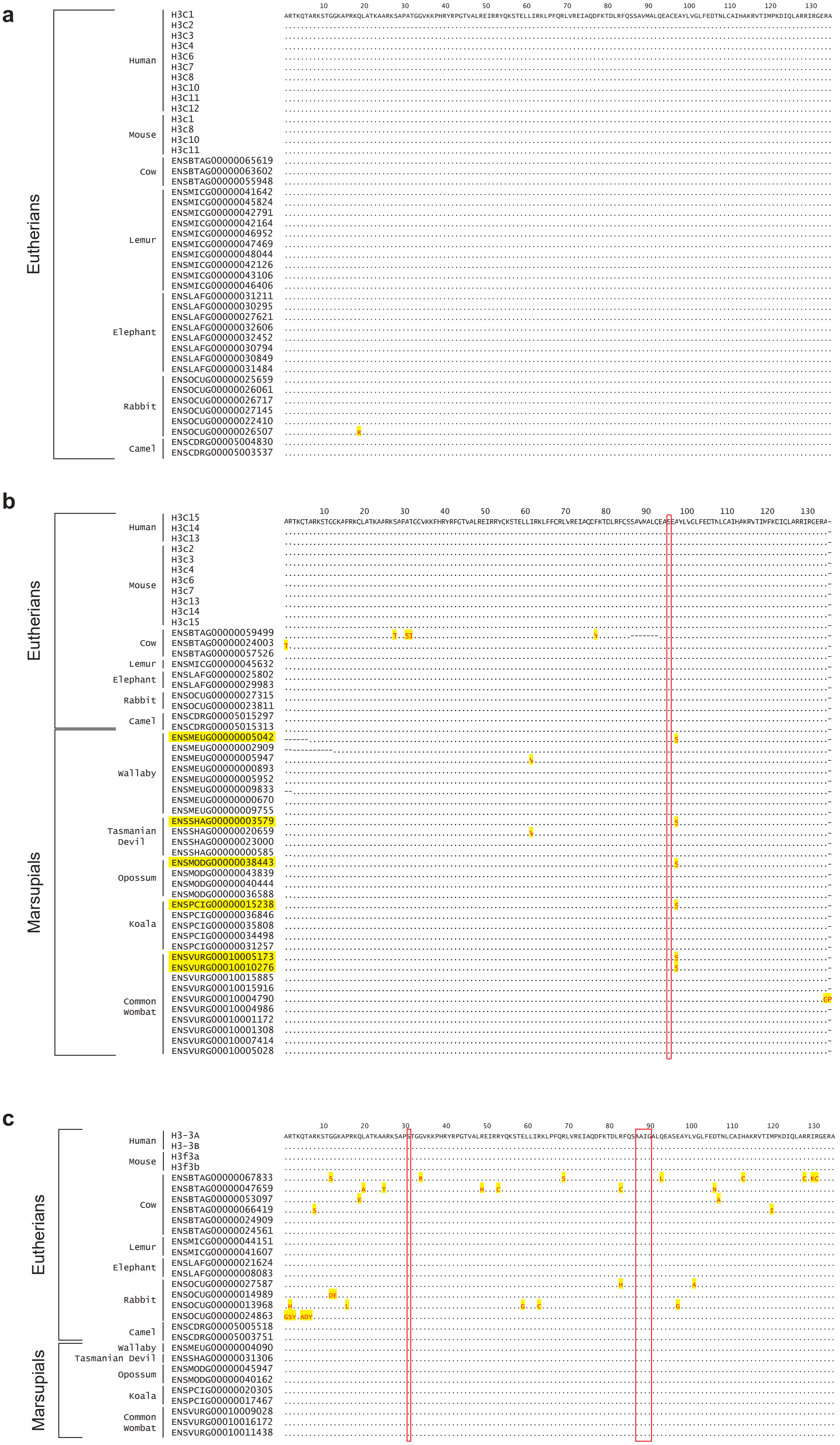

a
Eutherians
Human
H3C1
H3C2
H3C3
H3C4
H3C6
H3C7
H3C8
H3C10
H3C11
H3C12
Mouse
H3c1
H3c8
H3c10
H3c11
Cow
ENSBTAG00000065619
ENSBTAG00000063602
ENSBTAG00000055948
Lemur
ENSMICG00000041642
ENSMICG00000045824
ENSMICG00000042791
ENSMICG00000042164
ENSMICG00000046952
ENSMICG00000047469
ENSMICG00000048044
ENSMICG00000042126
ENSMICG00000043106
ENSMICG00000046406
Elephant
ENSLAFG00000031211
ENSLAFG00000030295
ENSLAFG00000027621
ENSLAFG00000032606
ENSLAFG00000032452
ENSLAFG00000030794
ENSLAFG00000030849
ENSLAFG00000031484
Rabbit
ENSOCUG00000025659
ENSOCUG00000026061
ENSOCUG00000026717
ENSOCUG00000027145
ENSOCUG00000022410
ENSOCUG00000026507
Camel
ENSCDRG00005004830
ENSCDRG00005003537
10 20 30 40 50 60 70 80 90 100 110 120 130
ARTKQTARKSTGGKAPRKQLATKAARKSAPATGGVKKPHRYRPGTVALREIRRYQKSTELLIRKLPFQRLVREIAQDFKTDLRFQSSAVMALQEACEAYLVGLFEDTNLCAIHAKRVTIMPKDIQLARRIRGERA
b
Eutherians
Human
H3C15
H3C14
H3C13
Mouse
H3c2
H3c3
H3c4
H3c6
H3c7
H3c13
H3c14
H3c15
Cow
ENSBTAG00000059499
ENSBTAG00000024003
ENSBTAG00000057526
Lemur
ENSMICG00000045632
Elephant
ENSLAFG00000025802
ENSLAFG00000029983
Rabbit
ENSOCUG00000027315
ENSOCUG00000023811
Camel
ENSCDRG00005015297
ENSCDRG00005015313
Marsupials
Wallaby
ENSMEUG00000005042
ENSMEUG00000002909
ENSMEUG00000005947
ENSMEUG00000000893
ENSMEUG00000005952
ENSMEUG00000009833
ENSMEUG00000000670
ENSMEUG00000009755
Tasmanian Devil
ENSSHAG00000003579
ENSSHAG00000020659
ENSSHAG00000023000
ENSSHAG00000000585
Opossum
ENSMODG00000038443
ENSMODG00000043839
ENSMODG00000040444
ENSMODG00000036588
Koala
ENSPCIG00000015238
ENSPCIG00000036846
ENSPCIG00000035808
ENSPCIG00000034498
ENSPCIG00000031257
Common Wombat
ENSVURG00010005173
ENSVURG00010010276
ENSVURG00010015885
ENSVURG00010015916
ENSVURG00010004790
ENSVURG00010004986
ENSVURG00010001172
ENSVURG00010001308
ENSVURG00010007414
ENSVURG00010005028
10 20 30 40 50 60 70 80 90 100 110 120 130
c
Eutherians
Human
H3-3A
H3-3B
Mouse
H3f3a
H3f3b
Cow
ENSBTAG00000067833
ENSBTAG00000047659
ENSBTAG00000053097
ENSBTAG00000066419
ENSBTAG00000024909
ENSBTAG00000024561
Lemur
ENSMICG00000044151
ENSMICG00000041607
Elephant
ENSLAFG00000021624
ENSLAFG00000008083
Rabbit
ENSOCUG00000027587
ENSOCUG00000014989
ENSOCUG00000013968
ENSOCUG00000024863
Camel
ENSCDRG00005005518
ENSCDRG00005003751
Marsupials
Wallaby
ENSMEUG00000004090
Tasmanian Devil
ENSSHAG00000031306
Opossum
ENSMODG00000045947
ENSMODG00000040162
Koala
ENSPCIG00000020305
ENSPCIG00000017467
Common Wombat
ENSVURG00010009028
ENSVURG00010016172
ENSVURG00010011438
10 20 30 40 50 60 70 80 90 100 110 120 130
ARTKQTARKSTGGKAPRKQLATKAARKSAPSTGGVKKPHRYRPGTVALREIRRYQKSTELLIRKLPFQRLVREIAQDFKTDLRFQSAAIGALQEASEAYLVGLFEDTNLCAIHAKRVTIMPKDIQLARRIRGERA

**Supplementary Figure 2. Sequences of H3.1, H3.2 and H3.3 in selected placental and marsupial mammals**

**a-c**, Amino acid sequence alignments of H3.1 (**a**), H3.2 (**b**) and H3.3 (**c**) proteins in selected placental and marsupial species. Marsupial *H3.2* genes that are orthologous to placental *H3-4* are highlighted in yellow.

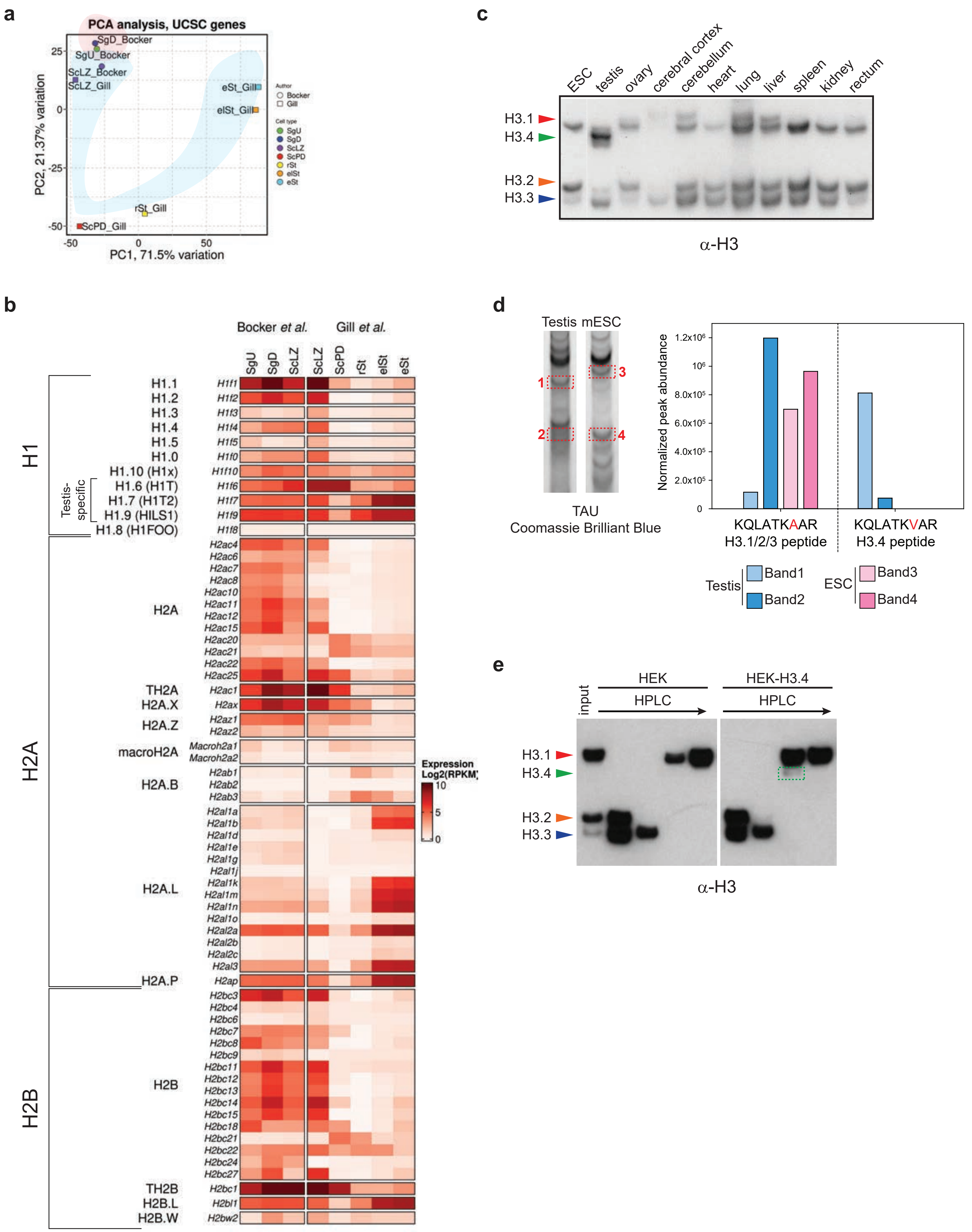

a
PCA analysis, UCSC genes
SgD_Bocker
SgU_Bocker
ScLZ_Bocker
ScLZ_Gill
eSt_Gill
elSt_Gill
rSt_Gill
ScPD_Gill
PC2, 21.37% variation
PC1, 71.5% variation
Author
Bocker
Gill
Cell type
SgU
SgD
ScLZ
ScPD
rSt
elSt
eSt
b
Bocker et al.
Gill et al.
SgU
SgD
ScLZ
ScLZ
ScPD
rSt
elSt
eSt
H1
H1.1
H1.2
H1.3
H1.4
H1.5
H1.0
H1.10 (H1x)
H1.6 (H1T)
H1.7 (H1T2)
H1.9 (HILS1)
H1.8 (H1FOO)
Testis-specific
H1f1
H1f2
H1f3
H1f4
H1f5
H1f0
H1f10
H1f6
H1f7
H1f9
H1f8
H2A
H2A
H2ac4
H2ac6
H2ac7
H2ac8
H2ac10
H2ac11
H2ac12
H2ac15
H2ac20
H2ac21
H2ac22
H2ac25
TH2A
H2ac1
H2A.X
H2ax
H2A.Z
H2az1
H2az2
macroH2A
Macroh2a1
Macroh2a2
H2A.B
H2ab1
H2ab2
H2ab3
H2A.L
H2al1a
H2al1b
H2al1d
H2al1e
H2al1g
H2al1j
H2al1k
H2al1m
H2al1n
H2al1o
H2al2a
H2al2b
H2al2c
H2al3
H2A.P
H2ap
H2B
H2B
H2bc3
H2bc4
H2bc6
H2bc7
H2bc8
H2bc9
H2bc11
H2bc12
H2bc13
H2bc14
H2bc15
H2bc18
H2bc21
H2bc22
H2bc24
H2bc27
TH2B
H2bc1
H2B.L
H2bl1
H2B.W
H2bw2
Expression Log2(RPKM)
10
5
0
c
ESC
testis
ovary
cerebral cortex
cerebellum
heart
lung
liver
spleen
kidney
rectum
H3.1
H3.4
H3.2
H3.3
α-H3
d
Testis
mESC
1
2
3
4
TAU
Coomassie Brilliant Blue
Normalized peak abundance
KQLATKAAR
H3.1/2/3 peptide
KQLATKVAR
H3.4 peptide
Testis
Band1
Band2
ESC
Band3
Band4
e
input
HEK
HPLC
HEK-H3.4
HPLC
H3.1
H3.4
H3.2
H3.3
α-H3

**Supplementary Figure 3. The expression of H3 isoforms in mouse tissues**

**a**, Principal component analysis (PCA) comparing RNA-seq expression of UCSC-annotated genes in different cell types[52,53]. The RNA-seq libraries of the two independent studies recapitulate the respective developmental trajectories during spermatogenesis

**b**, Heatmap showing mRNA expression H1, H2A and H2B encoding genes during spermatogenesis. The names of histone genes are according to the current nomenclature[34]. Abbreviations: SgU, SgD, ScLZ, ScPD, rSt, elSt and eSt refer to populations of FACS-sorted undifferentiated and differentiating spermatogonia, leptotene/zygotene and pachytene/diplotene stage spermatocytes and round, elongating and elongated spermatids.

**c**, Western blot showing H3.1, H3.4, H3.2 and H3.3 proteins separated by TAU gel electrophoresis in extracts of mouse embryonic stem cells and various tissues isolated from adult mice. A panH3 antibody recognizing the C-terminus of H3 to detect all H3 isoforms.

**d**, Mass spectrometry analysis of histones isolated from testes and mESC. We separated histones by TAU gel electrophoresis and stained them with Coomassie Brilliant Blue. We excised histone protein containing bands and subjected them to mass spectrometry. We quantified normalized peak abundances corresponding to the H3.1/H3.2/H3.3 (KQLATK$A_{24}$AR, left) common peptide and the H3.4 (KQLAT$V_{24}$AR, right) specific peptide in the four excised bands. The diagnostic amino acid variants between the two peptides are highlighted in red. Band 1 contains: H3.4, band 2: H3.3, band 3: H3.1 and band 4: H3.3.

**e**, Proteins blots showing the migration characteristics of H3.4 relative to other H3 proteins separated by TAU gel electrophoresis. We prepared histone extracts from control HEK293 cells or HEK293 cells expressing H3.4. We obtained different H3 protein containing fractions by reverse phase HPLC separation and subjected them to TAU gel electrophoresis.

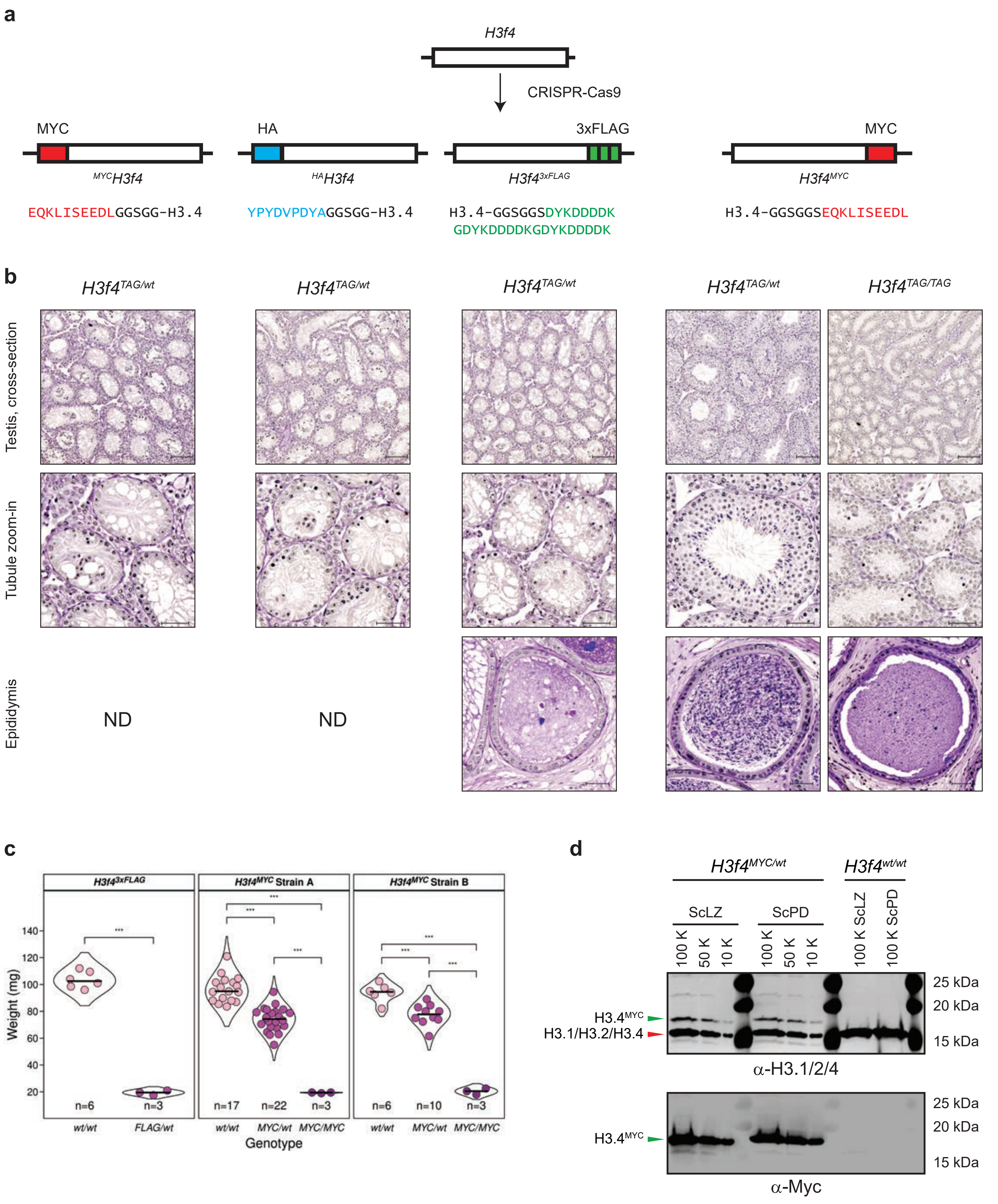
a
H3f4
CRISPR-Cas9
MYC
HA
3xFLAG
MYC
MYCH3f4
HAH3f4
H3f4 3xFLAG
H3f4 MYC
EQKLISEEDLGGSGG-H3.4
YPYDVPDYAGGSGG-H3.4
H3.4-GGSGGSDYKDDDDK
GDYKDDDDKGDYKDDDDK
H3.4-GGSGGSEQKLISEEDL
b
H3f4 TAG/wt
H3f4 TAG/wt
H3f4 TAG/wt
H3f4 TAG/wt
H3f4 TAG/TAG
Testis, cross-section
Tubule zoom-in
Epididymis
ND
ND
c
H3f4 3xFLAG
H3f4 MYC Strain A
H3f4 MYC Strain B
Weight (mg)
n=6
n=3
n=17
n=22
n=3
n=6
n=10
n=3
wt/wt
FLAG/wt
wt/wt
MYC/wt
MYC/MYC
wt/wt
MYC/wt
MYC/MYC
Genotype
d
H3f4 MYC/wt
H3f4 wt/wt
ScLZ
ScPD
100 K
50 K
10 K
100 K
50 K
10 K
100 K ScLZ
100 K ScPD
25 kDa
20 kDa
15 kDa
H3.4 MYC
H3.1/H3.2/H3.4
α-H3.1/2/4
H3.4 MYC
α-Myc

**Supplementary Figure 4. CRISPR/Cas9 approaches to tag the endogenous *H3f4* gene**

**a**, Schematic overview of the four different transgenic mouse lines generated by CRISPR/Cas9-mediated tagging of the endogenous *H3f4* gene. The sequences of protein tags are displayed by different colours and linker sequences in black. Sequences of sgRNAs and repair templates are provided in Supplementary Table 5 and 6.

**b**, PAS-haematoxylin-stained histological cross-sections of testis and epididymal tissues obtained from mice bearing tagged *H3f4* genes. Scale bars: 100 µm (testis, cross-section overview) and 50 µm (tubule zoom-in and epididymis). ND: not determined.

**c**, Violin plots showing testicular weights of control, heterozygous and homozygous transgenic mice of the $H3f4^{3xFLAG}$ strain and the two independent $H3f4^{MYC}$ strains. In the violin plots, each dot represents the mean weight of two testes taken from each mouse. The number (n) of mice per genotype are indicated. Paired t-test was performed between the indicated genotypes: *** $P \leq 0.001$.

**d**, SDS-PAGE Western blot analysis of $H3.4^{MYC}$ expression in FACS-purified ScLZ and ScPD cells of wild type $H3f4^{wt/wt}$ and heterozygous $H3f4^{MYC/wt}$ mice. Blots were probed with antibodies recognizing H3.1/H3.2/H3.4 (upper panel) and MYC (lower panel) epitopes. Bands corresponding to the tagged $H3.4^{MYC}$ and untagged H3.1, H3.2 and H3.4 proteins are indicated by arrows.

**a**

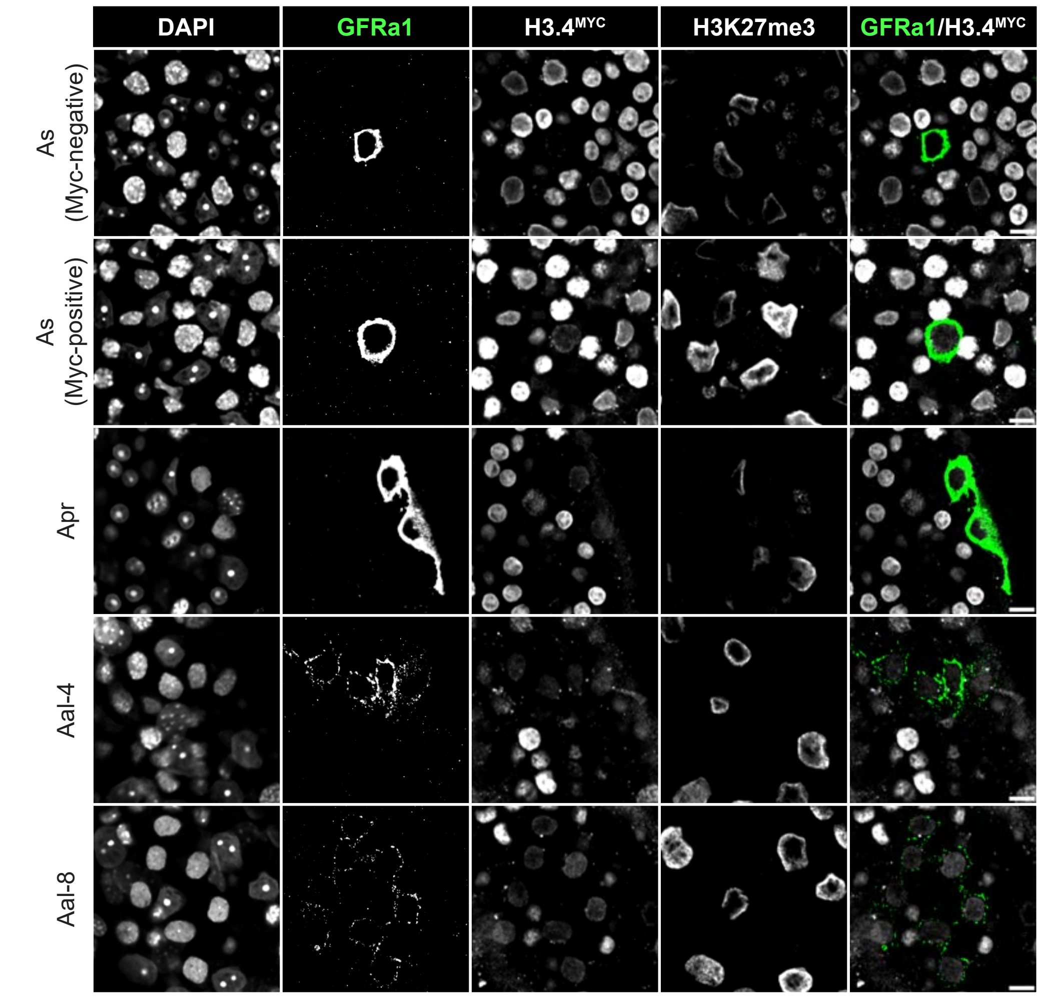

DAPI
GFRa1
H3.4MYC
H3K27me3
GFRa1/H3.4MYC
As (Myc-negative)
As (Myc-positive)
Apr
Aal-4
Aal-8


**b**

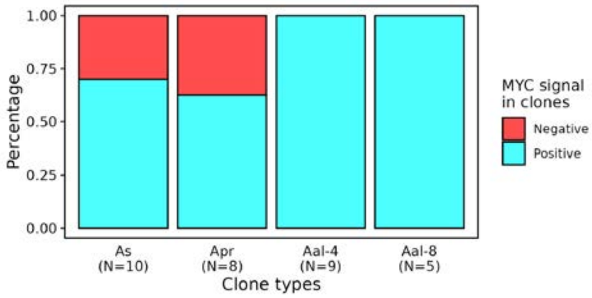

1.00
0.75
0.50
0.25
0.00
Percentage
As (N=10)
Apr (N=8)
Aal-4 (N=9)
Aal-8 (N=5)
Clone types
MYC signal in clones
Negative
Positive

**Supplementary Figure 5. H3.4-Myc is expressed in undifferentiated spermatogonia**

**a**, Whole-mount immunofluorescence images showing H3.4$^{MYC}$ expression in different clones of GRFα1-positive spermatogonia with different chain length: $A_s$ (single), $A_{pr}$ (paired), $A_{al\text{-}4}$ (chain of 4 aligned cells) and $A_{al\text{-}8}$ (chain of 8 aligned cells). For $A_s$, an example of a clone positive or negative for H3.4$^{MYC}$ are displayed. Scale bar: 10 μm.

**b**, Stacked bar plot showing the proportion of GRFα1-positive spermatogonia clones ($A_s$, $A_{pr}$, $A_{al\text{-}4}$ and $A_{al\text{-}8}$) positive or negative for H3.4$^{MYC}$ staining. n corresponds to the number of clones assessed.

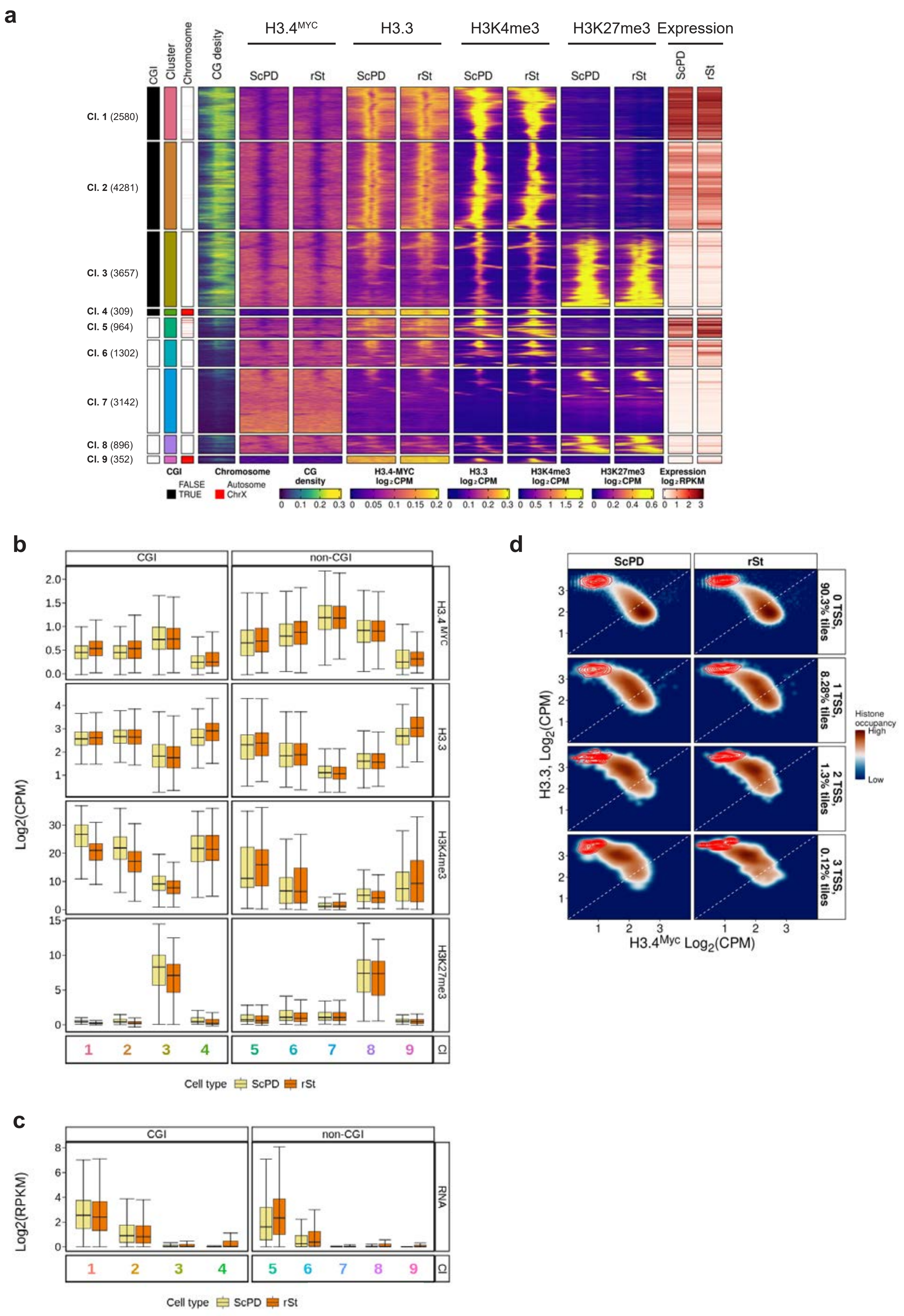
a
H3.4MYC
H3.3
H3K4me3
H3K27me3
Expression
CGI
Cluster
Chromosome
CG desity
ScPD
rSt
Cl. 1 (2580)
Cl. 2 (4281)
Cl. 3 (3657)
Cl. 4 (309)
Cl. 5 (964)
Cl. 6 (1302)
Cl. 7 (3142)
Cl. 8 (896)
Cl. 9 (352)
CGI
FALSE
TRUE
Chromosome
Autosome
ChrX
CG density
H3.4-MYC log2CPM
H3.3 log2CPM
H3K4me3 log2CPM
H3K27me3 log2CPM
Expression log2RPKM
b
CGI
non-CGI
H3.4 MYC
H3.3
H3K4me3
H3K27me3
Log2(CPM)
Cl
Cell type
ScPD
rSt
c
CGI
non-CGI
RNA
Log2(RPKM)
Cl
Cell type
ScPD
rSt
d
ScPD
rSt
0 TSS, 90.3% tiles
1 TSS, 8.28% tiles
2 TSS, 1.3% tiles
3 TSS, 0.12% tiles
H3.3, Log2(CPM)
H3.4Myc Log2(CPM)
Histone occupancy
High
Low

**Supplementary Figure 6. Genome localization of H3 isoforms in ScPD and rSt cells**

**a**, Heatmap displaying chromatin and transcriptional states at TSS regions (± 5 kb) of CGI- and nonCGI-containing genes measured in FACS-purified ScPD and rSt cells. Regions were grouped into 9 gene clusters by k-means clustering performed separately for the CGI- and for the non-CGI-type promoters performed on chromatin and RNA variables. The number of gene promoters is displayed for each cluster. CG density (±1 kb from TSS), chromatin and RNA expression levels are indicated in percentage, logCPM and log2RPKM respectively. The ChIP-seq for H3K4me3 and H3K27me3 data in rSt were obtained from Erkek *et al.*[56] and RNA expression data in ScPD and rSt from Gill *et al.*[53]

**b**, Boxplot showing quantification of ChIP-seq data (in log2CPM) in windows ±1 kb around TSS in clusters 1-9 as displayed in panel **a.** The boxes extend from the first quartile (Q1) to the third quartile (Q3) of the data distribution, with a line in the box representing the median. Whiskers indicate variability outside Q1 and Q3, extending to the minimum and maximum values within 1.5 times the interquartile range (IQR), while points outside this range represent outliers.

**c**, Boxplot showing quantification of expression of genes (in log2RPKM) belonging to the TSS regions displayed in clusters 1-9 in panel **a**. Boxplot parameters are as described in panel **b**.

**d**, Density scatter plots displaying log2CPM counts of H3.4$^{MYC}$ (x-axis) and H3.3 (y-axis) on genomic tiles that contain 0, 1, 2 or 3+ transcription units with TSS in ScPD and rSt cells. Red contour lines represent the density of tiles located on the X chromosome. The presence of a transcription unit with TSS was defined by an overlap of a given tile with a region ±1 kb around TSS.

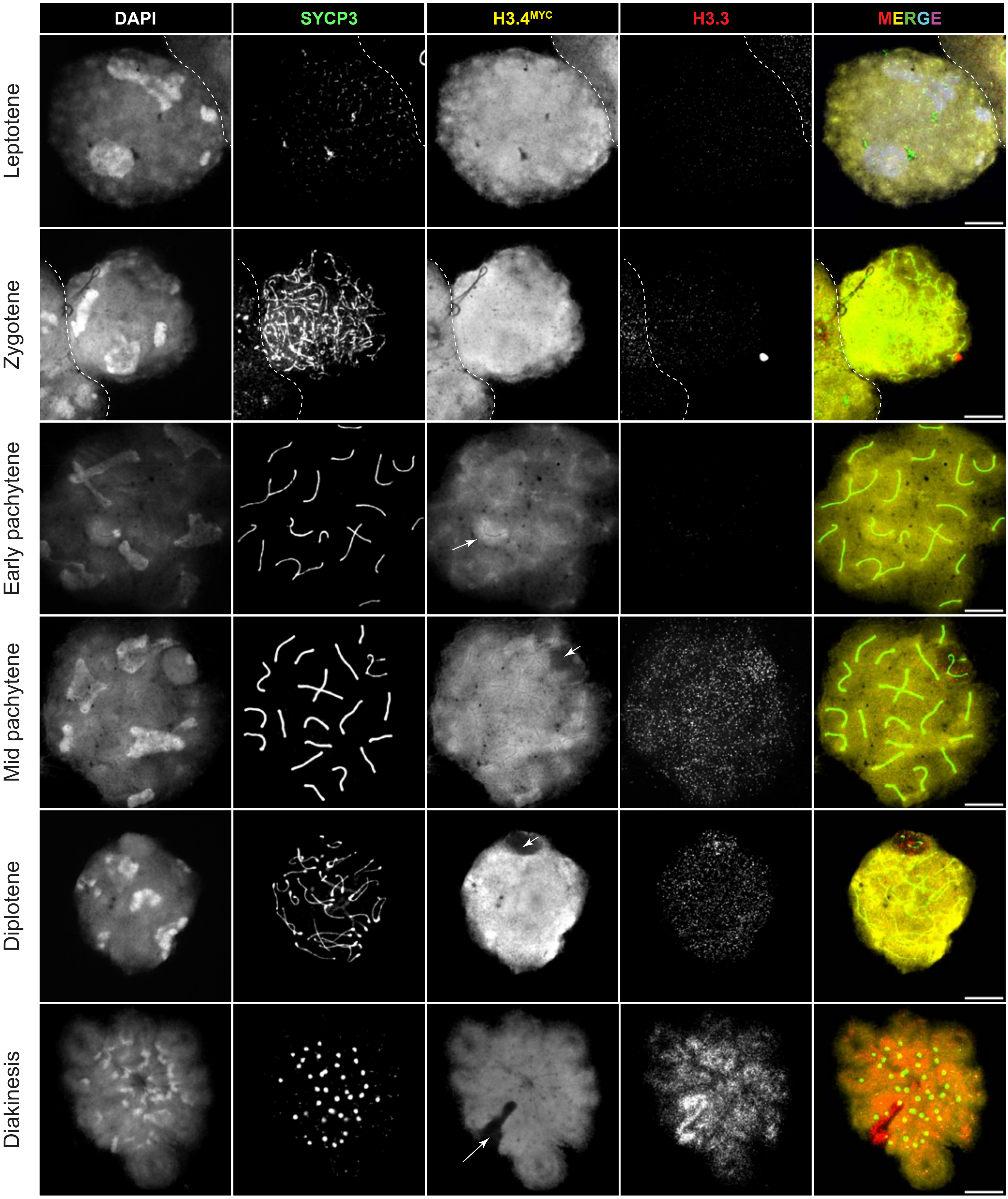
DAPI
SYCP3
H3.4MYC
H3.3
MERGE
Leptotene
Zygotene
Early pachytene
Mid pachytene
Diplotene
Diakinesis

**Supplementary Figure 7. H3.4 is evicted from the X and Y chromosomes in mid-pachytene spermatocytes**

Meiotic spreads showing temporal dynamics of H3.4$^{MYC}$ and H3.3 proteins during progression through meiotic prophase I. The X and Y chromosomes undergoing the eviction of H3.4 are indicated by a white arrow.

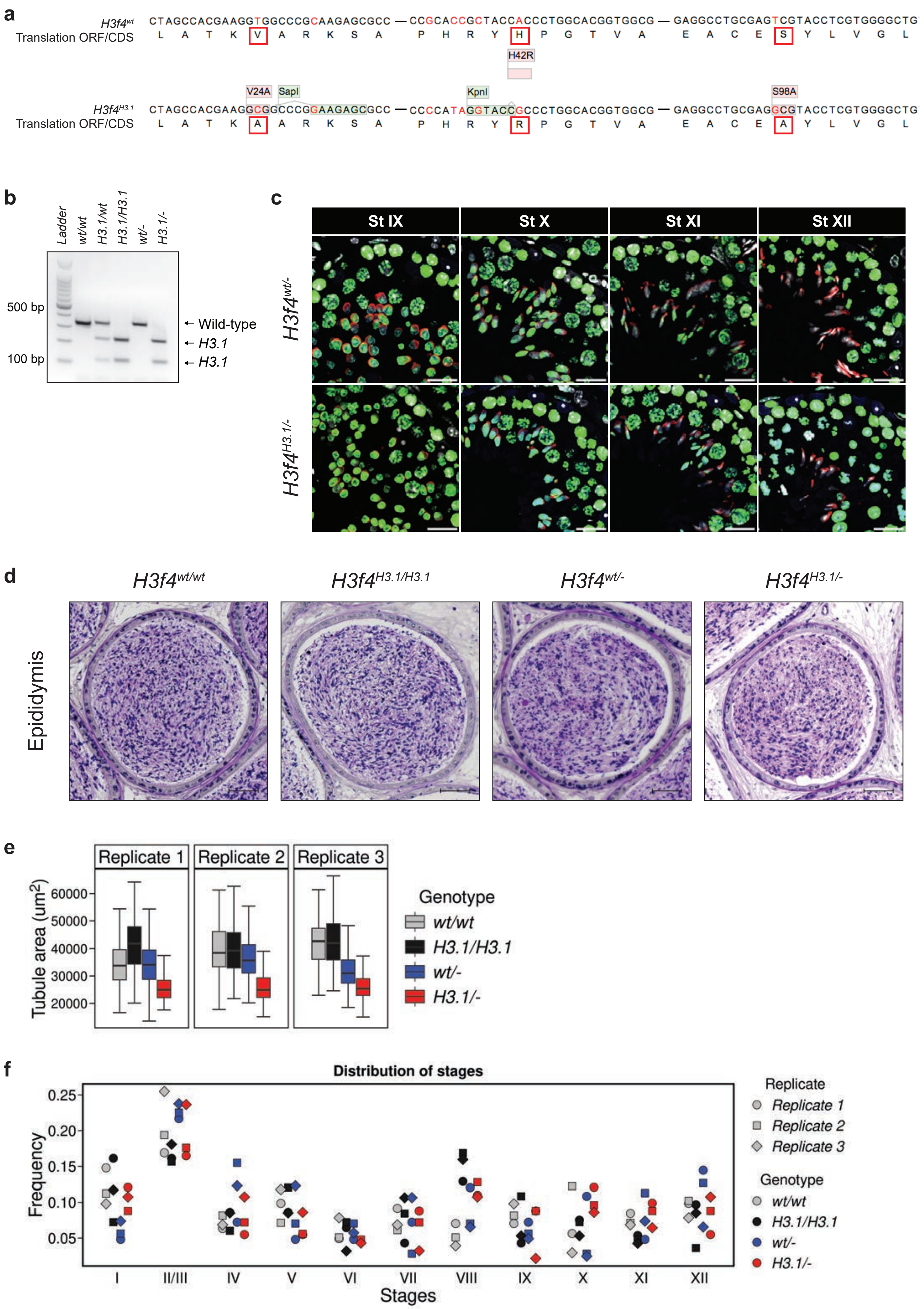
a
H3f4wt
CTAGCCACGAAGGTGGCCCGCAAGAGCGCC — CCGCACCGCTACCACCCTGGCACGGTGGCG — GAGGCCTGCGAGTCGTACCTCGTGGGGCTG
Translation ORF/CDS
L A T K V A R K S A P H R Y H P G T V A E A C E S Y L V G L
H42R
V24A
SapI
KpnI
S98A
H3f4H3.1
CTAGCCACGAAGGCGGCCCGGAAGAGCGCC — CCCCATAGGTACCGCCCTGGCACGGTGGCG — GAGGCCTGCGAGGCGTACCTCGTGGGGCTG
Translation ORF/CDS
L A T K A A R K S A P H R Y R P G T V A E A C E A Y L V G L
b
Ladder
wt/wt
H3.1/wt
H3.1/H3.1
wt/-
H3.1/-
500 bp
100 bp
← Wild-type
← H3.1
← H3.1
c
St IX
St X
St XI
St XII
H3f4wt/-
H3f4H3.1/-
d
H3f4wt/wt
H3f4H3.1/H3.1
H3f4wt/-
H3f4H3.1/-
Epididymis
e
Replicate 1
Replicate 2
Replicate 3
Tubule area (um2)
60000
50000
40000
30000
20000
Genotype
wt/wt
H3.1/H3.1
wt/-
H3.1/-
f
Distribution of stages
Frequency
0.25
0.20
0.15
0.10
0.05
I
II/III
IV
V
VI
VII
VIII
IX
X
XI
XII
Stages
Replicate
Replicate 1
Replicate 2
Replicate 3
Genotype
wt/wt
H3.1/H3.1
wt/-
H3.1/-

**Supplementary Figure 8. Generation of the *H3f4$^{H3.1}$* allele by CRISPR/Cas9 gene editing**

**a**, Schematic representation of the *H3f4$^{wt}$* wild type and *H3f4$^{H3.1}$* modified alleles, with corresponding H3.4 to H3.1-specific amino acid substitutions (in red boxes). Variant positions in DNA sequences encoding H3.4 and H3.1 are indicated in red, while relevant restriction sites are shown in light green. Abbreviations: V24A, Valine 24 to Alanine; H42R, Histidine 42 to Arginine; S98A, Serine 98 to Alanine. CDS, coding sequence; ORF, open reading frame.

**b**, PCR genotyping results of the *H3.1* mouse model. The following genotypes are shown: *H3f4$^{wt/wt}$*, *H3f4$^{H3.1/wt}$*, *H3f4$^{H3.1/H3.1}$*, *H3f4$^{wt/-}$* and *H3f4$^{H3.1/-}$*. Primers and PCR protocols used for genotyping are provided in Supplementary Table 7 and 8. The PCR products were digested with KpnI into a 338bp restriction fragment for the wild type allele and two bands of 222 bp and 116 bp for the *H3f4$^{H3.1}$* allele. A hundred base pair ladder is shown for size comparison.

**c.** Representative immunofluorescence images of stage IX to XII seminiferous tubules, co-stained for DAPI and with an antibody that recognized H3.1, H3.2 and H3.4 (#ab34)[62]. Scale bar: 20 µm.

**d**, PAS-haematoxylin staining of caudal epidydimal tissues obtained from *H3f4$^{wt/wt}$*, *H3f4$^{H3.1/H3.1}$*, *H3f4$^{wt/-}$* and *H3f4$^{H3.1/-}$* mice. Scale bar: 50 µm.

**e**, Boxplots showing the area of segmented seminiferous tubules used in the quantification analysis shown in Fig. 4h. Data for three biological replicates per genotype are shown.

**f**, Distribution of stages of the seminiferous tubule cycle in testes of 3 mice per genotype to identify stage V and X tubules for further automated cell quantification analysis (Fig. 4h). Staging was manually performed as described before[61]. Each dot represents the frequency of a given stage per biological replicate.

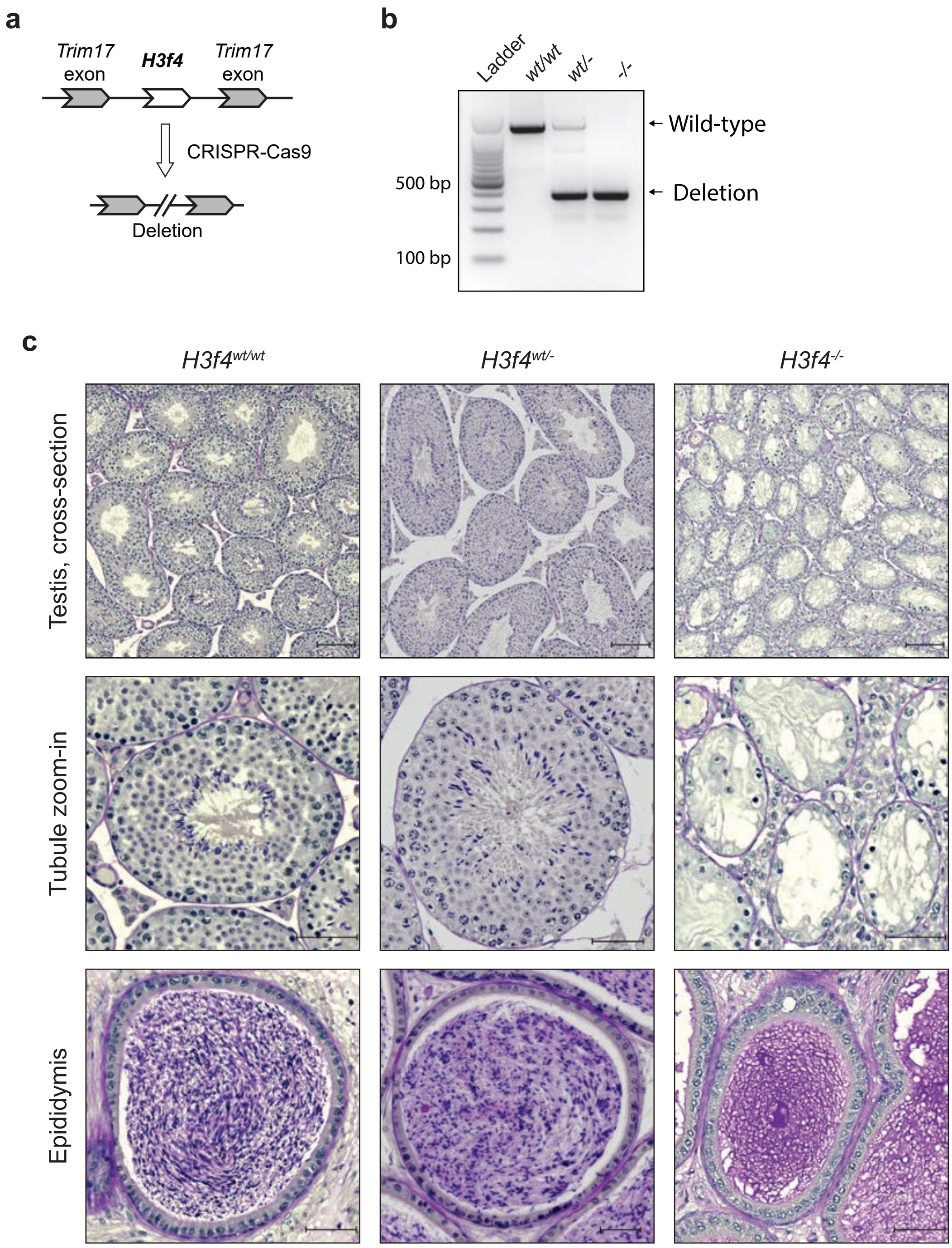

a
Trim17 exon
H3f4
Trim17 exon
CRISPR-Cas9
Deletion
b
Ladder
wt/wt
wt/-
-/-
500 bp
100 bp
Wild-type
Deletion
c
H3f4wt/wt
H3f4wt/-
H3f4-/-
Testis, cross-section
Tubule zoom-in
Epididymis

**Supplementary Figure 9. Deletion of the *H3f4* gene leads to infertility**

**a**, CRISPR/Cas9 strategy to delete the *H3f4* gene. Sequences of sgRNAs are indicated in Supplementary Table 5.

**b**, Genotyping results of mice with wild-type ($H3f4^{wt/wt}$), heterozygous ($H3f4^{wt/-}$) and *H3f4* knock-out ($H3f4^{-/-}$) genotypes. PCR using primers (Supplementary Tables 7 and 8) that bind up- and downstream of the deletion shows fragment sizes of 1180bp (wild-type allele) and 330bp (deleted allele). Ladder in multiples of hundred base pairs is shown for size comparison.

**c**, PAS-haematoxylin-stained histological cross-sections of testes and epididymal tissues obtained from $H3f4^{wt/wt}$, $H3f4^{wt/-}$ and $H3f4^{-/-}$ mice. Scale bars: 100 µm (cross-section overview) and 50 µm (tubule zoom-in and epididymis).

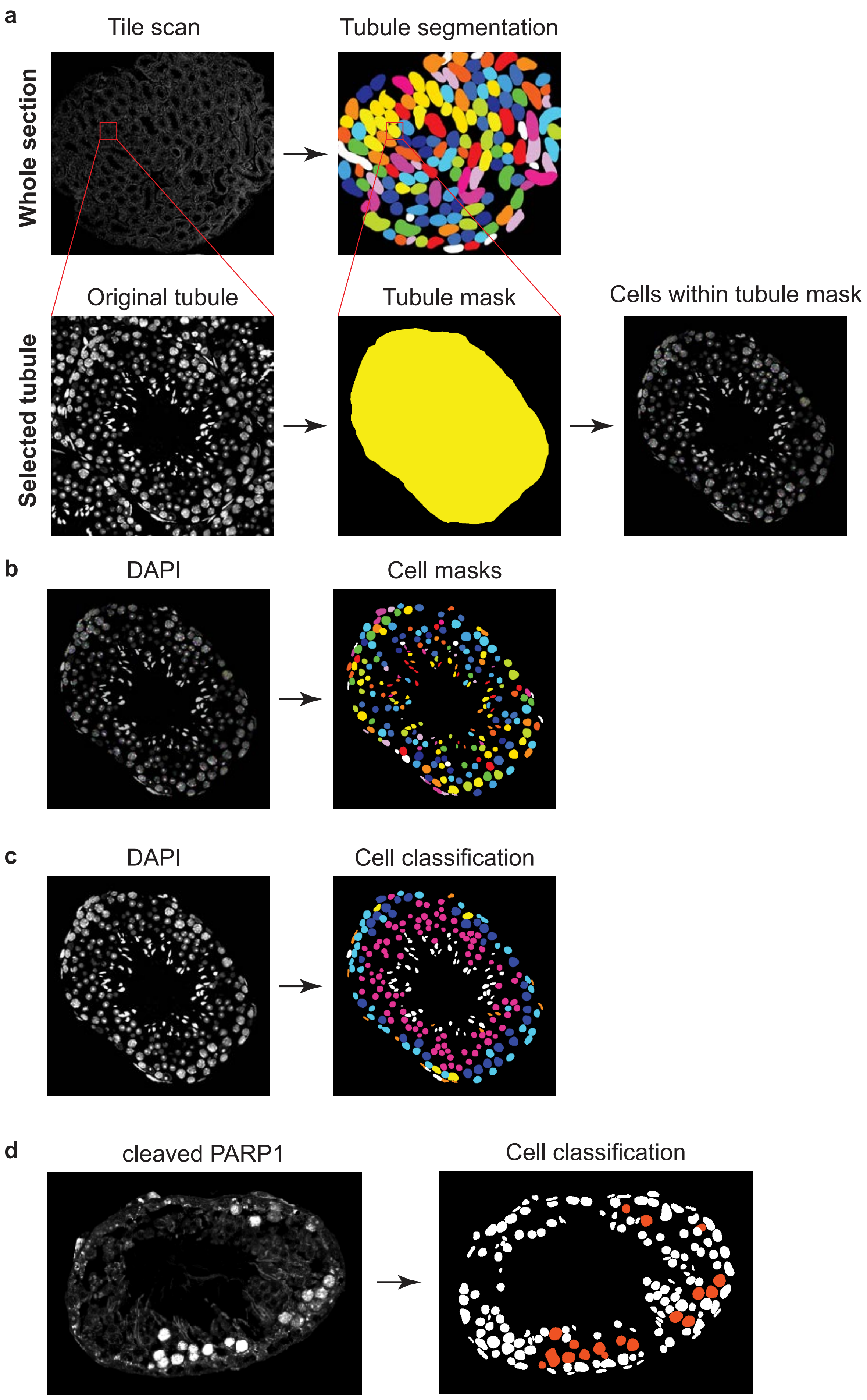

a
Tile scan
Tubule segmentation
Whole section
Original tubule
Tubule mask
Cells within tubule mask
Selected tubule
b
DAPI
Cell masks
c
DAPI
Cell classification
d
cleaved PARP1
Cell classification

**Supplementary Figure 10. The pipeline for automated quantification of male germ cells**

**a**, Example of tubule segmentation on an 8-binned stitched scan (DAPI channel used as an input). Raw DAPI and tubule masks are shown for the whole scan (upper panels) and for a selected tubule example (bottom).

**b**, Example of StarDist-based cell segmentation after manual curation of cell shapes. DAPI images were used as input information for cell segmentation.

**c,** Example of a multi-type cell classification results performed by Ilastik software[92] on a single channel (DAPI) channel. Ground truth annotation was used to assign classes to cells in a learning phase. Different colours represent cell different cell types.

**d**, Example of a single-type cell classification results performed by Ilastik software[92] on a single channel (cleaved PARP1) channel. Ground truth annotation was used to assign classes to cells in a learning phase.

a

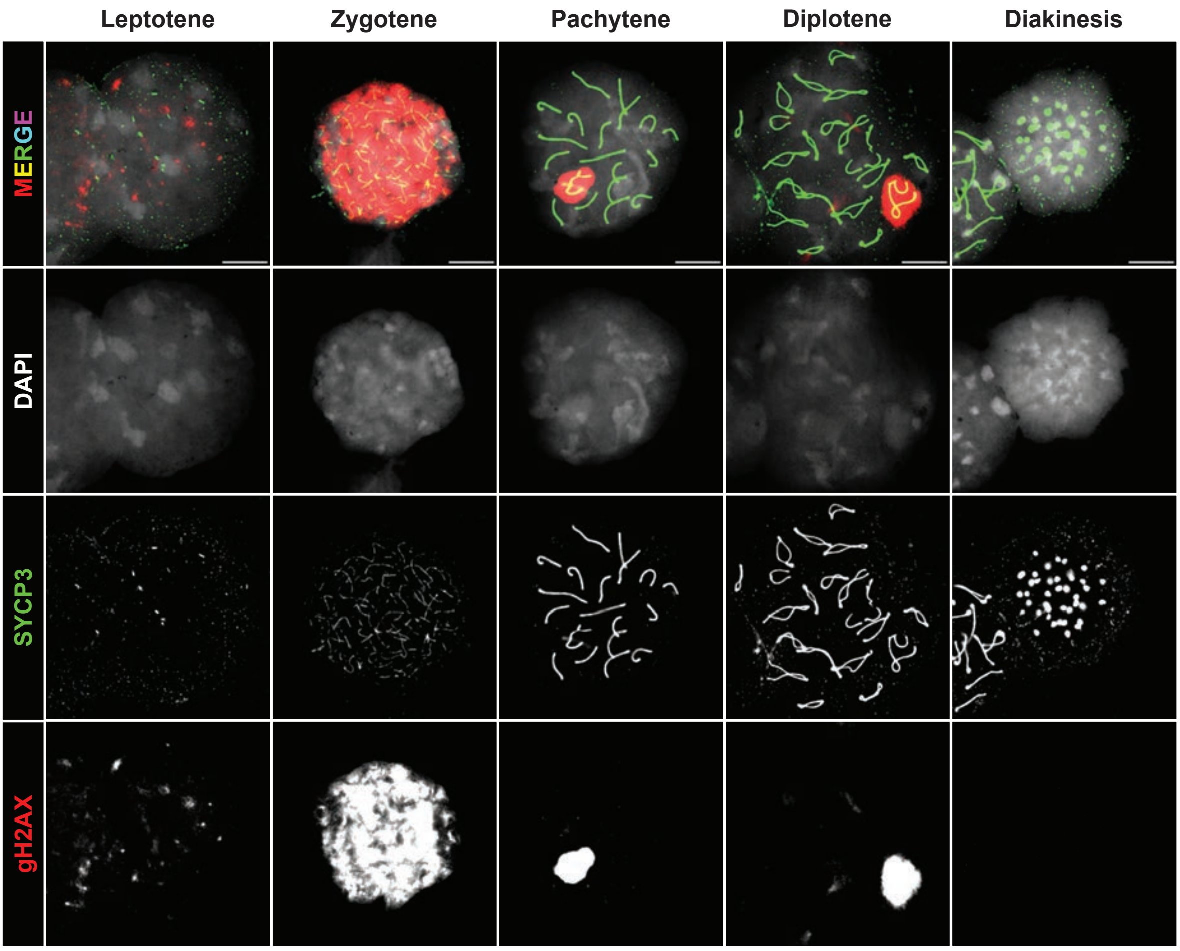
Leptotene
Zygotene
Pachytene
Diplotene
Diakinesis
MERGE
DAPI
SYCP3
gH2AX

b

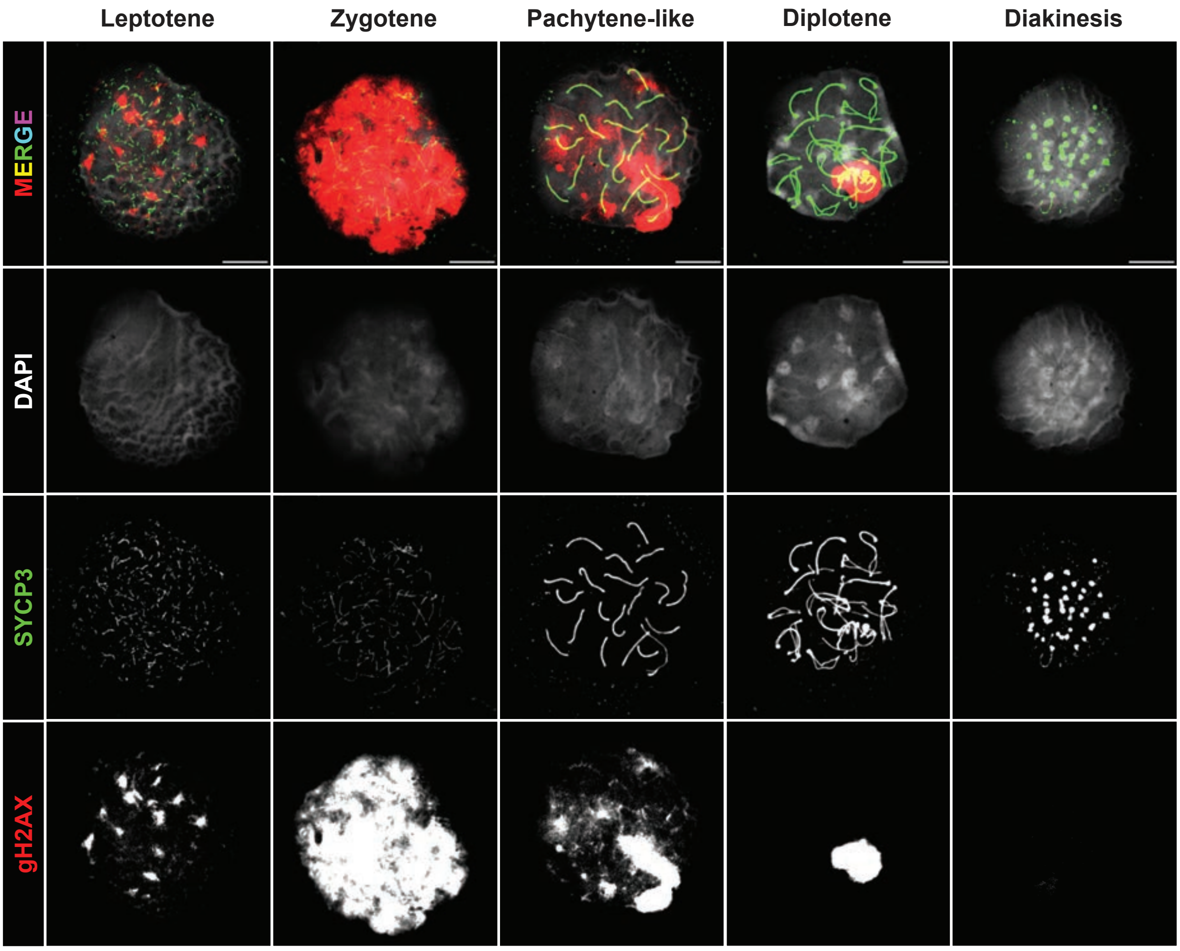
Leptotene
Zygotene
Pachytene-like
Diplotene
Diakinesis
MERGE
DAPI
SYCP3
gH2AX

**Supplementary Figure 11. Developmental course of meiotic prophase in *$H3f4^{wt/-}$* and *$H3f4^{H3.1/-}$* mice.**

**a-b,** Examples of cells in meiotic prophase aligned according to their developmental trajectories in *$H3f4^{wt/-}$* (a) and *$H3f4^{H3.1/-}$* (b) mice. Representative images of leptotene, zygotene, pachytene (pachytene-like) and diplotene spermatocytes are shown, together with cells in diakinesis. Scale bar: 10 μm.

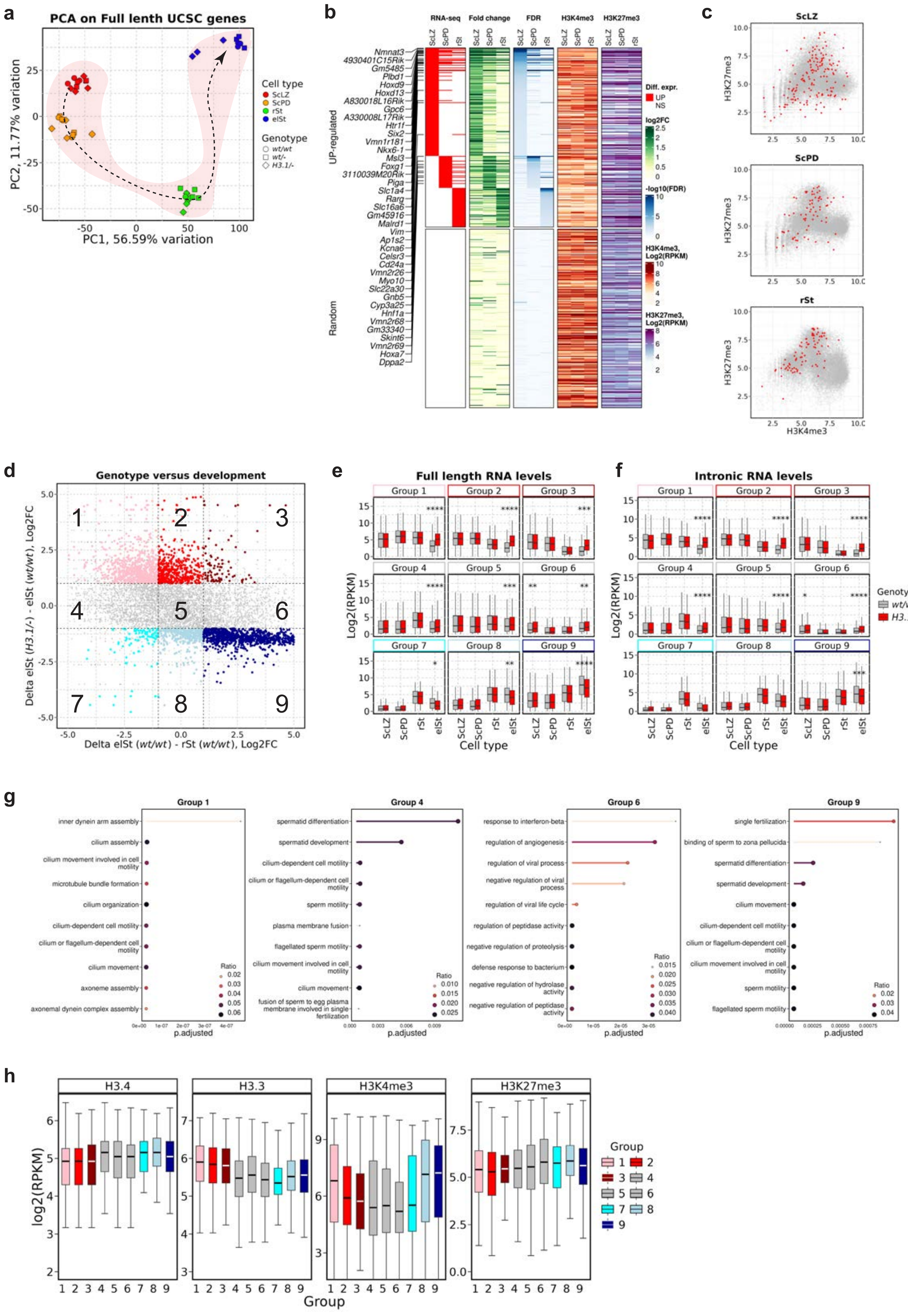
a
PCA on Full lenth UCSC genes
PC1, 56.59% variation
PC2, 11.77% variation
Cell type
ScLZ
ScPD
rSt
elSt
Genotype
wt/wt
wt/-
H3.1/-
b
RNA-seq
Fold change
FDR
H3K4me3
H3K27me3
UP-regulated
Random
Diff. expr.
UP
NS
log2FC
-log10(FDR)
H3K4me3, Log2(RPKM)
H3K27me3, Log2(RPKM)
c
ScLZ
ScPD
rSt
H3K27me3
H3K4me3
d
Genotype versus development
Delta elSt (H3.1/-) - elSt (wt/wt), Log2FC
Delta elSt (wt/wt) - rSt (wt/wt), Log2FC
e
Full length RNA levels
f
Intronic RNA levels
Log2(RPKM)
Cell type
Genotype
wt/wt
H3.1/-
g
Group 1
Group 4
Group 6
Group 9
p.adjusted
Ratio
h
H3.4
H3.3
H3K4me3
H3K27me3
log2(RPKM)
Group

**Supplementary Figure 12. Major transcriptional changes in elongating spermatids in *H3f4*$^{H3.1/-}$ mice.**

**a**, Principal component analysis (PCA) of RNA expression data (full length genic counts on UCSC-annotated genes), comparing biological replicates and genotypes in samples that were used in the study. The developmental trajectory is shown with the arrow.

**b**, Heatmap showing H3K4me3 and H3K27me3 ChIP-seq counts (±1 kb around TSS) on the promoters of UP-regulated genes in ScLZ, ScPD and rSt cells (Fig. 6a). A randomly selected cohort of genes is added below (same number of genes as for the upregulated). The EdgeR results (significance, fold-change and FDR) are shown for each gene. The genes commonly upregulated in at least two cell populations are indicated on the left. For panels b and c, H3K27me3 in ScLZ and ScPD was obtained from Bocker *et al.*[52], H3K4me3 and H3K27me3 in rSt was obtained from Erkek et al.[56]

**c**, Scatter plots showing the enrichment of H3K4me3 versus H3K27me3 at the promoters (±1 kb around TSS) of all genes (grey) and of the genes upregulated in ScLZ, ScPD and rSt cells (Fig. 6a) (red).

**d**, Scatter plots comparing two contrasts in non-CGI type promoter genes: genotype and mutation. The x axis represents the log2 fold change (FC) in gene expression during the transition from rSt to elSt in wild type (*H3f4*$^{wt/wt}$) cells (wild-type elSt versus wild-type rSt, max. FDR = 0.05). The y axis represents the log2FC in gene expression in elSt cells caused by the *H3f4*$^{H3.1/-}$ mutation (*H3f4*$^{H3.1/-}$ versus *H3f4*$^{wt/wt}$, max. FDR = 0.05). Genes UP- and DOWN-regulated in *H3f4*$^{H3.1/-}$ mutant cells are highlighted in shades of red and blue, respectively. The DEGs are separated into nine groups, based on the |log2FC| = 1 cutoff in both contrasts (dashed lines). To be included in groups 1-3 and 7-9, only the DEGs were considered.

**e-f**, Absolute quantification of gene expression in groups 1-9 defined in panel d, comparing wild type (*H3f4*$^{wt/wt}$) and mutant (*H3f4*$^{H3.1/-}$) cells. Panels e and f show full-length genic and intronic counts, respectively.

**g**, GO term search (GO: Biological Process) for genes in groups 1, 4, 6 and 9 (panel D).

**h**, Boxplots displaying wild type rSt ChIP-seq profiles of different chromatin signatures (H3.4$^{MYC}$, H3.3, H3K4me3 and H3K27me3 counted in ±1 kb around TSS) in groups 1-9 defined in panel D. The colour coding of groups is according to panel D.

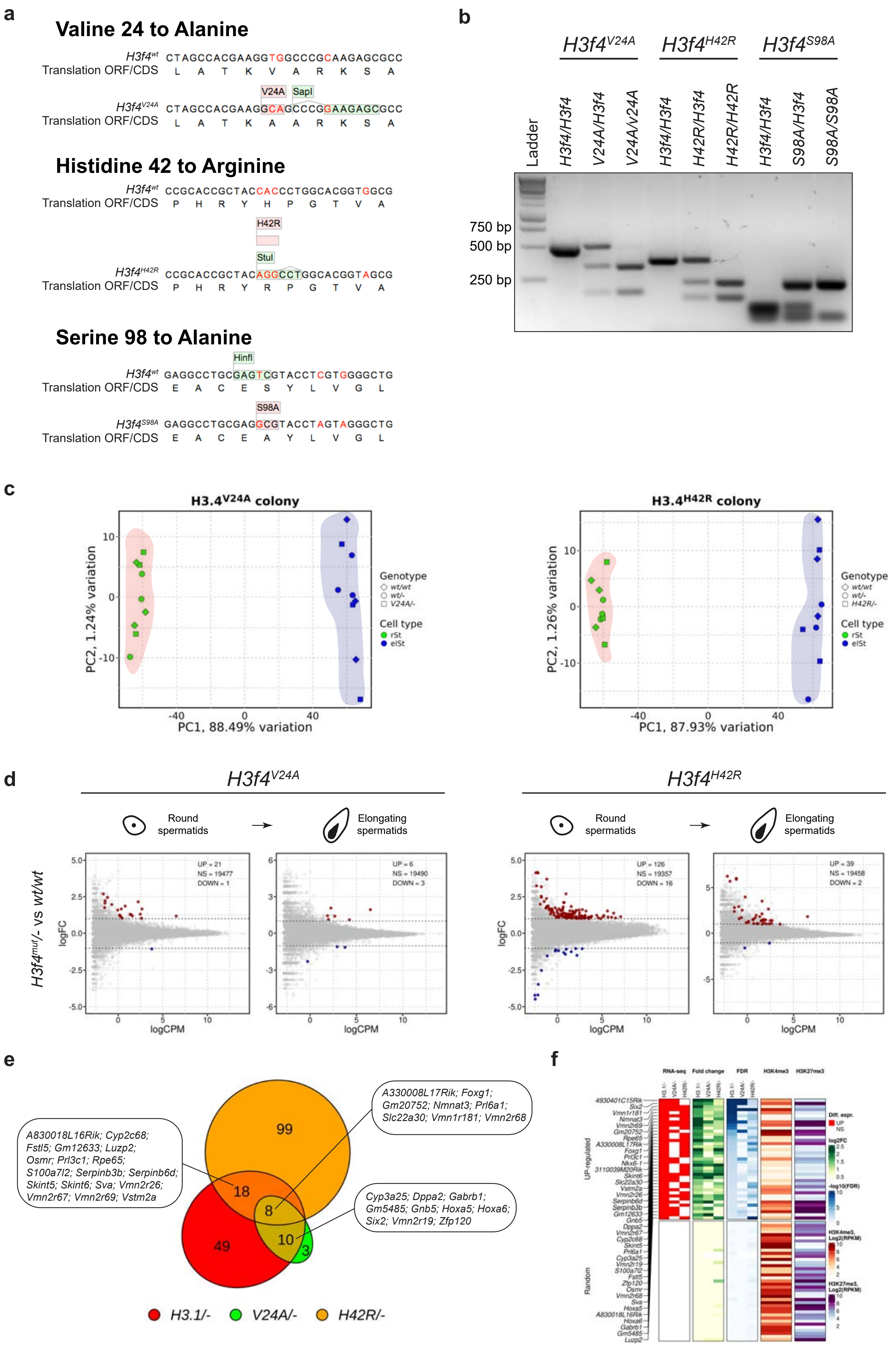
a
Valine 24 to Alanine
H3f4wt CTAGCCACGAAGGTGGCCCGCAAGAGCGCC
Translation ORF/CDS L A T K V A R K S A
V24A
SapI
H3f4V24A CTAGCCACGAAGGCAGCCCGGAAGAGCGCC
Translation ORF/CDS L A T K A A R K S A
Histidine 42 to Arginine
H3f4wt CCGCACCGCTACCACCCTGGCACGGTGGCG
Translation ORF/CDS P H R Y H P G T V A
H42R
StuI
H3f4H42R CCGCACCGCTACAGGCCTGGCACGGTAGCG
Translation ORF/CDS P H R Y R P G T V A
Serine 98 to Alanine
HinfI
H3f4wt GAGGCCTGCGAGTCGTACCTCGTGGGGCTG
Translation ORF/CDS E A C E S Y L V G L
S98A
H3f4S98A GAGGCCTGCGAGGCGTACCTAGTAGGGCTG
Translation ORF/CDS E A C E A Y L V G L
b
H3f4V24A
H3f4H42R
H3f4S98A
Ladder
H3f4/H3f4
V24A/H3f4
V24A/V24A
H3f4/H3f4
H42R/H3f4
H42R/H42R
H3f4/H3f4
S98A/H3f4
S98A/S98A
750 bp
500 bp
250 bp
c
H3.4V24A colony
PC2, 1.24% variation
PC1, 88.49% variation
Genotype
wt/wt
wt/-
V24A/-
Cell type
rSt
elSt
H3.4H42R colony
PC2, 1.26% variation
PC1, 87.93% variation
Genotype
wt/wt
wt/-
H42R/-
Cell type
rSt
elSt
d
H3f4V24A
H3f4H42R
Round spermatids
Elongating spermatids
H3f4mut/- vs wt/wt
logFC
logCPM
UP = 21
NS = 19477
DOWN = 1
UP = 6
NS = 19490
DOWN = 3
UP = 126
NS = 19357
DOWN = 16
UP = 39
NS = 19458
DOWN = 2
e
99
18
8
49
10
3
A830018L16Rik; Cyp2c68; Fstl5; Gm12633; Luzp2; Osmr; Prl3c1; Rpe65; S100a7l2; Serpinb3b; Serpinb6d; Skint5; Skint6; Sva; Vmn2r26; Vmn2r67; Vmn2r69; Vstm2a
A330008L17Rik; Foxg1; Gm20752; Nmnat3; Prl6a1; Slc22a30; Vmn1r181; Vmn2r68
Cyp3a25; Dppa2; Gabrb1; Gm5485; Gnb5; Hoxa5; Hoxa6; Six2; Vmn2r19; Zfp120
H3.1/-
V24A/-
H42R/-
f
RNA-seq
Fold change
FDR
H3K4me3
H3K27me3
UP-regulated
Random
Diff. expr.
UP
NS
log2FC
-log10(FDR)
H3K4me3, Log2(RPKM)
H3K27me3, Log2(RPKM)

**Supplementary Figure 13. Individual contributions of H3.4-specific amino acids to the support of spermatogenesis.**

**a**, Schematic representation of *H3f4* point substitution alleles. Variant positions in DNA sequence between wild type *H3f4* (up) and point mutation alleles (down) are indicated in red. For each mouse model, the DNA sequence of the mutated position within *H3f4* is shown and the translation is shown below. Changes in amino acid sequence are indicated in light red and relevant restriction sites are shown in light green. Changes in DNA sequence that are not annotated are silent mutations that were added to prevent cutting of the sgRNA after successful recombination. Abbreviations: CDS, coding sequence; ORF, open reading frame.

**b**, Genotyping results of wild-type, heterozygous and homozygous mice from colonies with H3.1-specific point mutations. Fragments were PCR amplified (Supplementary Tables 7 and 8) and digested with the following restriction enzymes: SapI for *H3f4*$^{V24A}$; StuI for *H3f4*$^{H42R}$; HinfI for *H3f4*$^{S98A}$ allele. Fragment sizes after nuclease digestion: SapI: 516 bp (*H3f4*$^{wt/wt}$), 340 bp + 176 bp (*H3f4*$^{V24A/V24A}$); StuI: 384 bp + 27 bp (shorter band not shown) (*H3f4*$^{wt/wt}$), 227 bp + 157 bp + 27 bp (shorter band not shown) (*H3f4*$^{H42R/H42R}$); HinfI: 125 bp + 107 bp + 76 bp (*H3f4*$^{wt/wt}$), 230 bp + 76 bp (*H3f4*$^{S98A/S98A}$). Ladder with indicated size (bp) is shown for comparison on the left.

**c**, Principal component analysis (PCA) on USCS-annotated genes comparing RNA-seq data obtained from FACS-isolated rSt and elSt cells with *H3f4*$^{wt/wt}$, *H3f4*$^{wt/-}$, *H3f4*$^{V24A/-}$ and *H3f4*$^{H42R/-}$ genotypes.

**d**, Scatter plots showing gene expression log2 fold changes in FACS-isolated rSt and elSt mutant cells plotted against average gene expression levels in *H3f4*$^{wt/wt}$ cells. Contrasts: *H3f4*$^{V24A/-}$ versus *H3f4*$^{wt/wt}$ and *H3f4*$^{H42R/-}$ versus *H3f4*$^{wt/wt}$. The numbers of significantly upregulated (red) or downregulated (blue) genes are displayed on the top right and genes names of the most highly significant differentially expressed genes are shown. EdgeR parameters: min. logFC = 1, max. FDR = 0.05.

**e**, Euler diagrams showing the overlap of DEGs in rSt cells in three different mutants: *H3f4*$^{H3.1/-}$ (Fig. 6a), *H3f4*$^{V24A/-}$ and *H3f4*$^{H42R/-}$ (Supplementary Fig. 13d).

**f**, Heatmap showing H3K4me3 and H3K27me3 ChIP-seq counts[56] (±1 kb around TSS) on the promoters of commonly UP-regulated genes in *H3f4*$^{H3.1/-}$, *H3f4*$^{V24A/-}$and *H3f4*$^{H42R/-}$ rSt cells. A randomly selected cohort of genes is added below (same number of genes as for the upregulated). The EdgeR results (significance, fold-change and FDR) are shown for each gene. The genes commonly upregulated in at least two cell populations are indicated on the left.

### Supplementary Table Legends

#### Supplementary Table 1. Mammalian orthologs of human *H3-4*

Orthologs of the human *H3-4* gene found in the genomes of 110 placental and marsupial species available in ENSEMBL (release 112). The first three rows represent the sequences of mouse H3.1, H3.2 and H3.3 used as a reference for Fig. 1a. For species with known *H3-4* orthologs, the corresponding coordinates, type and sequence (amino acid and nucleotide) are provided. For species with no *H3-4* orthologs, only the Latin name and TaxID are provided. Only functional genes were included for the alignment of protein sequences; pseudogenes were discarded. For each orthologous sequence that is excluded from the analysis, explanation is provided in the “Notes” column.

#### Supplementary Table 2. The organization of the *HIST3* cluster in marsupials

The genes within the marsupial *HIST3* locus are indicated and the links to the ENSEMBL are provided accordingly. Abbreviations: str – strand; psg – pseudogene; C. wombat – common wombat. The order of genes is according to the ENSEMBL genome browser.

#### Supplementary Table 3. Paralogs of *H3-4* in the selected placental and marsupial species

Paralogues of the *H3-4* identified in the selected seven placental (human, mouse, elephant, cow, lemur, rabbit, camel) and five marsupial (wallaby, Tasmanian Devil, opossum, koala, common wombat) species. For each gene, their ENSEMBL ID, coordinates, strand and symbol (if available) are indicated. The column “Keep” indicates, whether a paralogue was taken into the downstream analysis, or not. NoTr – number of transcripts. For the details, please see the Methods section.

#### Supplementary Table 4. Differentially genes due to H3.4 to H3.1 substitution

Genes that are differentially expressed in $H3f4^{H3.1/-}$ mutants in FACS-isolated ScLZ, ScPD, rSt and elSt populations, compared to wild type ($H3f4^{wt/wt}$) mice. Abbreviations, according to edgeR: LR – likelihood ratio, FDR – false discovery rate.

**Supplementary Table 5. Sequences of sgRNAs used for *H3f4* gene editing**

Sequences of sgRNAs used for *H3f4* gene editing and the corresponding alleles. Synthesis protocols are according to the Methods section.

**Supplementary Table 6. Sequences of homology templates used for *H3f4* gene editing**

Table showing the sequences of homology repair templates, their types and methods used for purification. The corresponding alleles are indicated.

**Supplementary Table 7. Sequences of oligonucleotides used for genotyping**

Table showing the names and sequences of oligonucleotides used in the study, as well as purposes of their use.

**Supplementary Table 8. Strategies used for genotyping of *H3f4* mutant alleles**

Table showing the PCR protocols used for the genotyping of *H3f4* mutant alleles.

**Supplementary Table 9. Antibodies used in the study**

Table showing the primary and secondary antibodies used in this study. The dilutions for each assay are indicated accordingly.
